# Development of approaches and identifying limitations towards functional yeast surface display of FLS2

**DOI:** 10.1101/2025.11.25.690463

**Authors:** Benedikt Dolgikh, Samantha Schulte, Daniel R. Woldring

## Abstract

Pattern recognition receptors such as FLAGELLIN SENSING 2 (FLS2) are central to plant immunity and attractive targets for engineering broader detection of bacterial phytopathogens for application in pest management (and diagnostics) in food crops, and sustainable agriculture practices. However, evaluating numerous FLS2 variants for altered pathogen sensing specificity directly in plants is slow and low throughput, and have been seldom optimized for heterologous display systems. Here, we established conditions that enabled *Arabidopsis thaliana* FLS2 ectodomain expression on the surface of *Saccharomyces cerevisiae* and evaluated binding to its cognate ligand, flg22. We show how yeast high-mannose glycosylation of the FLS2 ectodomain contributes to inefficient folding and loss of detectable flg22 binding in standard yeast surface display conditions. Substitutions at all N-glycosylation motifs compromised surface expression, indicating that some glycosylation is required for trafficking. We tuned the extent of glycosylation using tunicamycin, an N-linked glycosylation inhibitor, in combination with thermal stress to modulate ER quality control. Under these conditions, we observed a reproducible subpopulation of cells with improved flg22 binding despite reduced overall expression, and we confirmed flg22 selectivity against non-FLS2 proteins using both flow cytometry and magnetic bead-based enrichment. Guided by structural modeling of high-mannose glycans on the FLS2 ectodomain, we then substituted asparagines at selected N-glycan sites to serine. We identified a key glycan site variant, N388S, which lies proximal to the flg22 binding interface and increased the binding population size under stress conditions. Binding assays against FLS2 variants and a reported non-binding variant affirmed that FLS2 selectivity was specific to FLS2 display, showing that all variants maintained low affinity interaction with flg22. Together, these results point to FLS2 display conditions, not only glycosylation state, as an underlying limitation to detect true flg22 interactions which will require more sensitive approaches to confidently resolve true binding populations.

**For Table of Contents Use Only:** 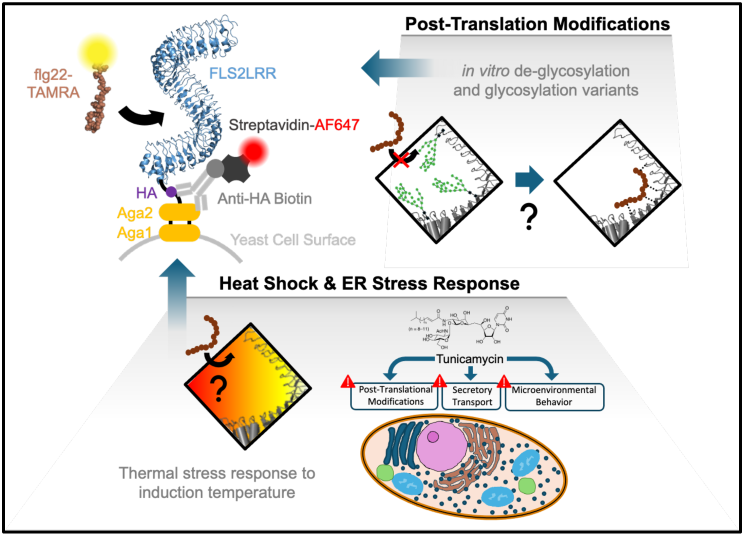

## Introduction

Global climate change and rapid pathogen evolution threaten sustainable agriculture by increasing disease pressure on major crops (*1–3*). For instance, Huanglongbing (citrus greening), caused by the bacterial phytopathogen *Candidatus Liberibacter asiaticus (CLas)*, has contributed to a ∼90% decline in Florida citrus production since the mid-2000s (*4–6*). It is widely supported that the efficiency of plant defense response to bacterial phytopathogens depends on plant host recognition sensitivity to flagellin pathogen-associated molecular patterns (PAMPs)(*7,8*). In the case of orange defense response against notable bacterial phytopathogens, weakened response to *CLas* and *Xanthomonas citri subsp. citri* is mainly attributed to PAMP sequence variation which weaken the recognition sensitivity of orange FLAGELLIN SENSING 2 (FLS2)(*9–12*). As a key cellular gatekeeper to recognize and initiate plant defense signaling, the pattern recognition receptor (PRR) FLS2 is a first line of defense against many bacterial pathogens. This leucine-rich repeat receptor-like kinase perceives conserved PAMPs in bacterial flagellin (flg22), initiating pattern-triggered immunity via BAK1 co-receptor recruitment and downstream signaling (*11,13–16*).

FLS2-flg22 recognition exhibits clear signatures of co-evolution: single amino acid changes in flg22 sequences can abolish perception by specific FLS2 variants, allowing pathogens to evade detection, whereas compensatory mutations in specific FLS2 orthologs restore or broaden responsiveness to previously unrecognized flg22 peptides (*13,17–20*). This tunability makes FLS2 a powerful candidate for protein engineering aimed at expanding the spectrum of detectable bacterial phytopathogens.

High-throughput engineering of FLS2 variants in planta remains challenging. Even with advances in CRISPR-Cas and nanoparticle-based delivery (*21,22*), generating and phenotyping transgenic plants typically supports libraries of only 10^2^-10^3^ variants over several months (*23*). Heterologous hosts (bacteria, insect cells, or mammalian cells) offer alternative routes for protein production, but often lack plant-like glycosylation or are not optimized for large-scale surface display campaigns (*24–30*). Yeast (*Saccharomyces cerevisiae* and *Pichia pastoris*) occupy a useful middle ground: they are eukaryotic, genetically tractable, and support yeast surface display platforms that routinely screen >10⁷ variants in weeks (*31–35*). However, FLS2 is an unusually large, heavily glycosylated receptor, and prior attempts to display its ectodomain on yeast failed to yield detectable ligand binding (*36*). Non-optimized binding conditions and differences in protein folding between plants and yeast, such as during endoplasmic reticulum quality control (ERQC) and N-linked glycosylation (*28,30,37–41*), may complicate functional expression. Thus, we set out to address the previous shortcomings in establishing a functional FLS2 ectodomain display system.

Here, we systematically investigate how yeast N-linked glycosylation and ER stress influence surface display and ligand binding of the *A. thaliana* FLS2 ectodomain. We show that FLS2 expressed on the yeast surface is extensively modified with high-mannose glycans, that complete removal of N-glycosylation motifs impairs trafficking, and that partial inhibition of glycosylation combined with thermal stress yields a subpopulation of cells with weak, but detectable flg22-positive display phenotype. Guided by structural modeling of high-mannose glycans on the FLS2 ectodomain, we identify the N388 glycan site, which is proximal to the flg22 binding interface, to be particularly disruptive to the flg22-positive display phenotype. These parameters provide clear suggestions for investigating an enrichment-compatible flg22-binding phenotype towards achieving functional yeast surface display of FLS2.

## Results and Discussion

### FLS2 is surface displayed but lacks detectable flg22 binding under standard yeast display conditions

As a first step toward developing a yeast display workflow for FLS2 engineering, we displayed the *A. thaliana* FLS2 ectodomain on the surface of *S. cerevisiae* as an Aga2 fusion (**Figure S1–S2**), hereafter termed FLS2LRR. Upon induction, flow cytometry showed that a high fraction of cells (>50%) were expression-positive, and Western blot analysis confirmed production of the FLS2LRR fusion protein (**Figure 1A,B; Figure S3**). However, despite this detectable surface expression, we did not observe flg22 binding under standard induction and labeling conditions (**Figure S3**). Thus, baseline yeast display supported FLS2LRR expression but did not produce a detectable ligand-binding receptor population.

**Figure 1.**
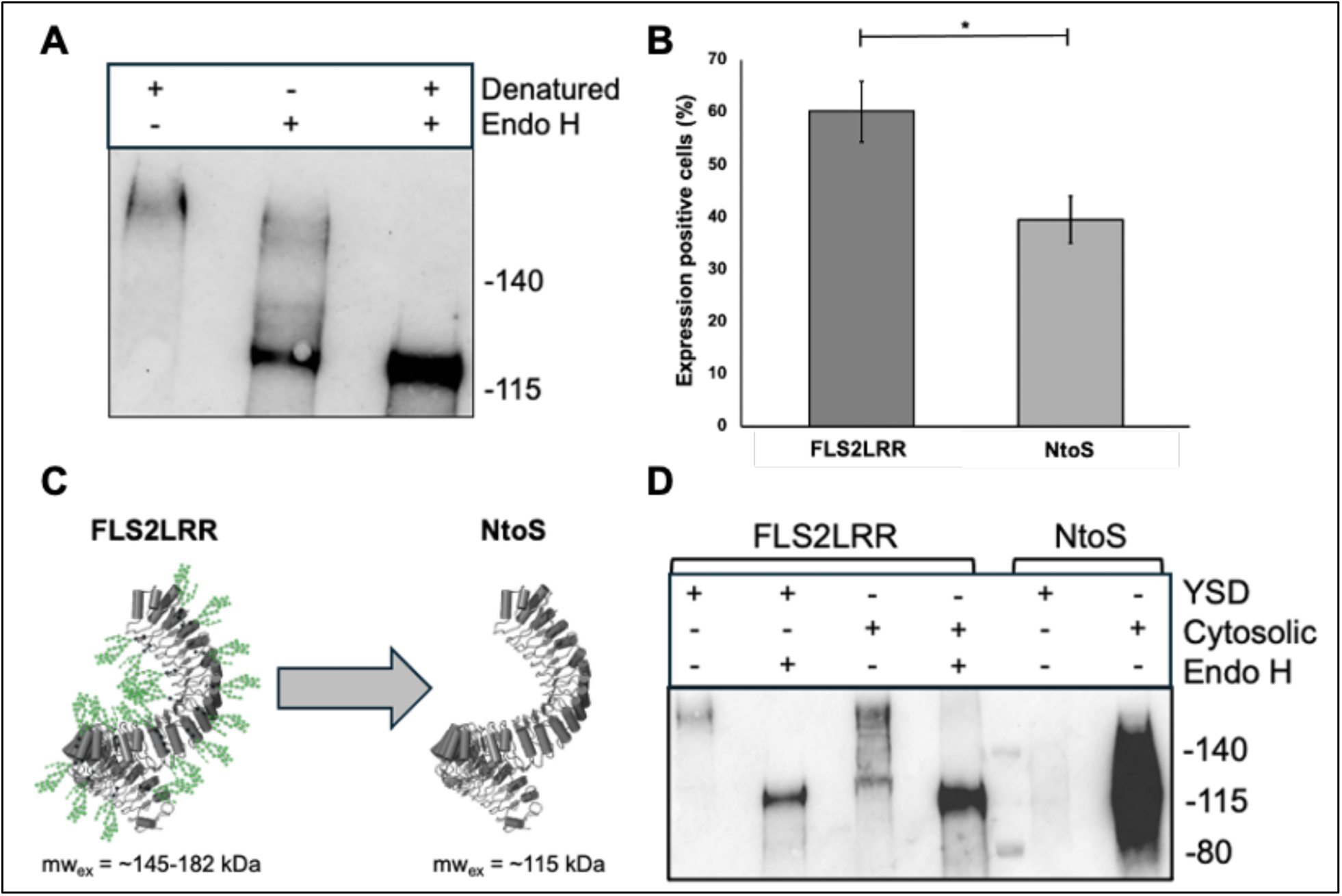
Yeast N-linked glycosylation supports maximal surface expression. (A) Western blot analysis of Aga2-HA-FLS2 (FLS2LRR) after Endo H treatment under denaturing or non-denaturing conditions attributes large kDa shift to yeast-specific glycosylation. (B) The mean percentage of FLS2LRR and NtoS populations with surface display expression. (C) Diagram of expected mass change for non-glycosylated Aga2-HA-FLS2 (NtoS) with N-linked glycan associated asparagines (22 sites) mutated to serine. (D) Western blot analysis of surface displayed (YSD) or cell lysate extracted (cytosolic) FLS2LRR and NtoS with or without denaturing Endo H treatment. For western blots, FLS2LRR and NtoS are detected using rat anti-HA-biotin (3F10) and streptavidin-647 antibodies. Cytometry data (C) shown is the mean and standard error for n=4 biological replicates per group (each bar) where p-value of * = < 0.05.

We next asked whether yeast-specific protein processing could contribute to the disconnect between surface expression and ligand binding. FLS2 orthologs contain numerous conserved N-linked glycosylation motifs (NXS/T, where X ≠ Pro)(*42*) (**Figure S4**). In plants, N-linked glycans are often complex or hybrid structures containing xylose and/or fucose, whereas S. cerevisiae predominantly produces high-mannose N-glycans (**Figure S5**)(*43*). Because these host-specific glycosylation differences could alter FLS2 folding, trafficking, or ligand accessibility, we examined the glycosylation state of yeast-displayed FLS2LRR.

The FLS2 ectodomain contains 22 putative N-linked glycosylation sites, and high-mannose occupancy at these sites would be expected to increase the apparent molecular weight by approximately 30–67 kDa (**Figure S6**), consistent with hyperglycosylation observed for other heterologous glycoproteins expressed in S. cerevisiae (*44*). To test whether FLS2LRR was modified by yeast-type N-glycans, we treated FLS2LRR display extracts with endoglycosidase H (Endo H) under denaturing and non-denaturing conditions. Under denaturing conditions, glycans should be broadly accessible to Endo H, whereas incomplete digestion under non-denaturing conditions would suggest that some glycan sites are less accessible in the folded or partially folded displayed protein (*45*).

Western blot analysis showed that untreated FLS2LRR migrated as a broad species above 140 kDa, whereas Endo H treatment under denaturing conditions shifted the protein to a sharper band near 115 kDa (**Figure 1A**). This large molecular-weight shift is consistent with extensive Endo H-sensitive high-mannose glycosylation of FLS2LRR in yeast. Endo H treatment under non-denaturing conditions produced an intermediate migration pattern, consistent with partial glycan removal and limited accessibility of some glycan sites. These results indicate that yeast-displayed FLS2LRR is heavily modified by high-mannose N-glycans. Together with the lack of detectable flg22 binding under standard conditions, these data suggest that baseline yeast display produces a surface-expressed but ligand-binding-deficient FLS2LRR population.

### Removal of all predicted N-glycosylation motifs reduces FLS2 surface display and increases intracellular retention

Because FLS2LRR was heavily modified by Endo H-sensitive high-mannose glycans in yeast yet lacked detectable flg22 binding under standard display conditions, we next asked whether reducing N-glycosylation could improve functional display. Specifically, we tested whether removing predicted N-glycosylation motifs would preserve FLS2 expression and trafficking while potentially reducing glycan-dependent effects on folding or ligand accessibility.

To identify a substitution strategy likely to preserve the FLS2 ectodomain structure, we evaluated alternative residues at all 22 predicted N-glycosylation-site asparagines. Candidate substitutions were guided by amino acid substitution frequencies (BLOSUM62 (*46*)) and modeled using AlphaFold2 Multimer (*47*) (**Figure S7A**). AF2-predicted structures were then analyzed using Rosetta stability calculations (*48–50*). Among the substitutions evaluated, replacing all 22 glycosylation-site asparagines with serines produced the smallest predicted deviation from the wild-type structure (**Figure S7B**). Based on this analysis, we generated an FLS2 ectodomain Aga2 fusion in which all 22 predicted N-glycosylation-site asparagines were substituted with serine, hereafter termed NtoS (**Figure 1C**). These may be advantageous compared to asparagine-to-aspartate substitutions described in the literature (*37*), given that serine physiochemical properties are more suitable for maintaining continuous hydrogen bonding to preserve fold at LRR motifs (*51*).

We compared surface expression of FLS2LRR and NtoS by flow cytometry. Across biological replicates (n = 3), the mean fraction of expression-positive cells decreased from 60.2±9.9% for FLS2LRR to 39.4±7.7% for NtoS (**Figure 1B**). Thus, removal of all predicted N-glycosylation motifs did not abolish display, but it reduced the fraction of cells with detectable surface expression. Because productive surface display requires folding, secretion, and retention of the Aga2 fusion at the cell surface (*52,53*), this reduction is consistent with impaired folding and/or trafficking of the NtoS construct.

Western blot analysis of surface-extracted and intracellular fractions further supported this interpretation. Although both FLS2LRR and NtoS were detected in intracellular fractions after induction, a larger fraction of NtoS protein was retained intracellularly compared with FLS2LRR (**Figure 1D**). This pattern suggests that removal of all predicted ectodomain N-glycosylation motifs increases intracellular retention and reduces efficient surface trafficking.

Together, these results indicate that complete removal of predicted N-glycosylation motifs compromises, but does not eliminate, FLS2 surface display in yeast. These data support a model in which some degree of N-glycosylation promotes productive folding and/or trafficking of Aga2–FLS2, whereas excessive or yeast-specific high-mannose glycosylation may still interfere with detectable ligand binding. This tradeoff motivated us to test whether intermediate glycosylation states, rather than complete glycan-site removal, could improve functional display (**Figure S8**).

### Partial glycosylation inhibition and thermal stress increase weak flg22-TAMRA-positive populations

#### Tunicamycin shifts FLS2LRR toward lower-molecular-weight glycoforms

Because FLS2LRR was heavily modified by high-mannose N-glycans in yeast and lacked detectable flg22 binding under standard conditions, we next asked whether reducing N-glycosylation could alter the flg22-TAMRA-positive population. Prior studies suggest that FLS2 glycosylation is important for receptor biogenesis in planta, but that reduced glycosylation is not necessarily incompatible with flg22 recognition if the receptor is properly folded (*37,38,54*). For example, Sun et al. reported that the majority FLS2 glycosylation-site variants, ranging from single to octuple glycan-site substitutions, retained flg22 responsiveness in planta, although reduced responsiveness in some mutants could reflect altered expression, trafficking, or binding (*37*). Similarly, the FLS2–flg22–BAK1 complex has been structurally characterized after enzymatic de-glycosylation of recombinantly expressed protein (*54*). Together, these findings imply that reduced glycosylation should not inherently disrupt flg22 binding as long as overall folding is preserved or achieved prior to de-glycosylation.

Because FLS2LRR is displayed on the yeast surface, enzymatic partial de-glycosylation is difficult to control. We therefore used tunicamycin, referred to as “Tm” in some figures, to perturb N-linked glycosylation during induction. Tunicamycin inhibits ALG7 early in lipid-linked oligosaccharide biosynthesis, reducing the availability of glycan precursors for oligosaccharyltransferase-mediated N-glycosylation (*55,56*). Importantly, tunicamycin is a global perturbation that can affect glycoprotein synthesis, ER stress, cell physiology, and viability (*57–59*); therefore, its effects cannot be attributed solely to changes at specific FLS2 glycosylation sites. We induced FLS2LRR-expressing cells in the presence of 0, 0.2, or 2 µg/mL tunicamycin and analyzed display extracts by Western blot. Increasing tunicamycin concentration shifted FLS2LRR toward lower-molecular-weight species, with 2 µg/mL producing the most pronounced shift (**Figure S9A**). This result is consistent with reduced N-glycosylation of FLS2LRR, although the specific occupied glycan sites and degree of site-to-site heterogeneity were not resolved in this assay.

#### Tunicamycin increases the fraction of flg22-TAMRA-positive FLS2LRR cells relative to non-FLS2 controls

Because tunicamycin globally perturbs glycoprotein biosynthesis and could increase nonspecific peptide binding (*55,60,61*), we compared flg22-TAMRA signal in FLS2LRR-displaying cells with two unrelated surface-displayed proteins, Aga2–HA–HaloTag and Aga2–HA–RIXI. This comparison allowed us to ask whether tunicamycin caused a general increase in flg22-TAMRA-positive cells or whether the increase was greater in the FLS2LRR-displaying population.

Across independent biological replicates (n = 3), the mean fraction of flg22-TAMRA-positive FLS2LRR cells increased from 0.2% without tunicamycin to 5.4% and 11.2% at 2 µg/mL tunicamycin in two replicate sets (**Figure 2A,B**). HaloTag- and RIXI-displaying cells also showed some increase in flg22-TAMRA-positive events, but the magnitude was not statistically significant, increasing from 2.6% to 3.4% and from 3.3% to 5.8%, respectively. Thus, tunicamycin increased flg22-TAMRA-positive events more strongly in FLS2LRR-displaying cells than in non-FLS2 controls.

**Figure 2.**
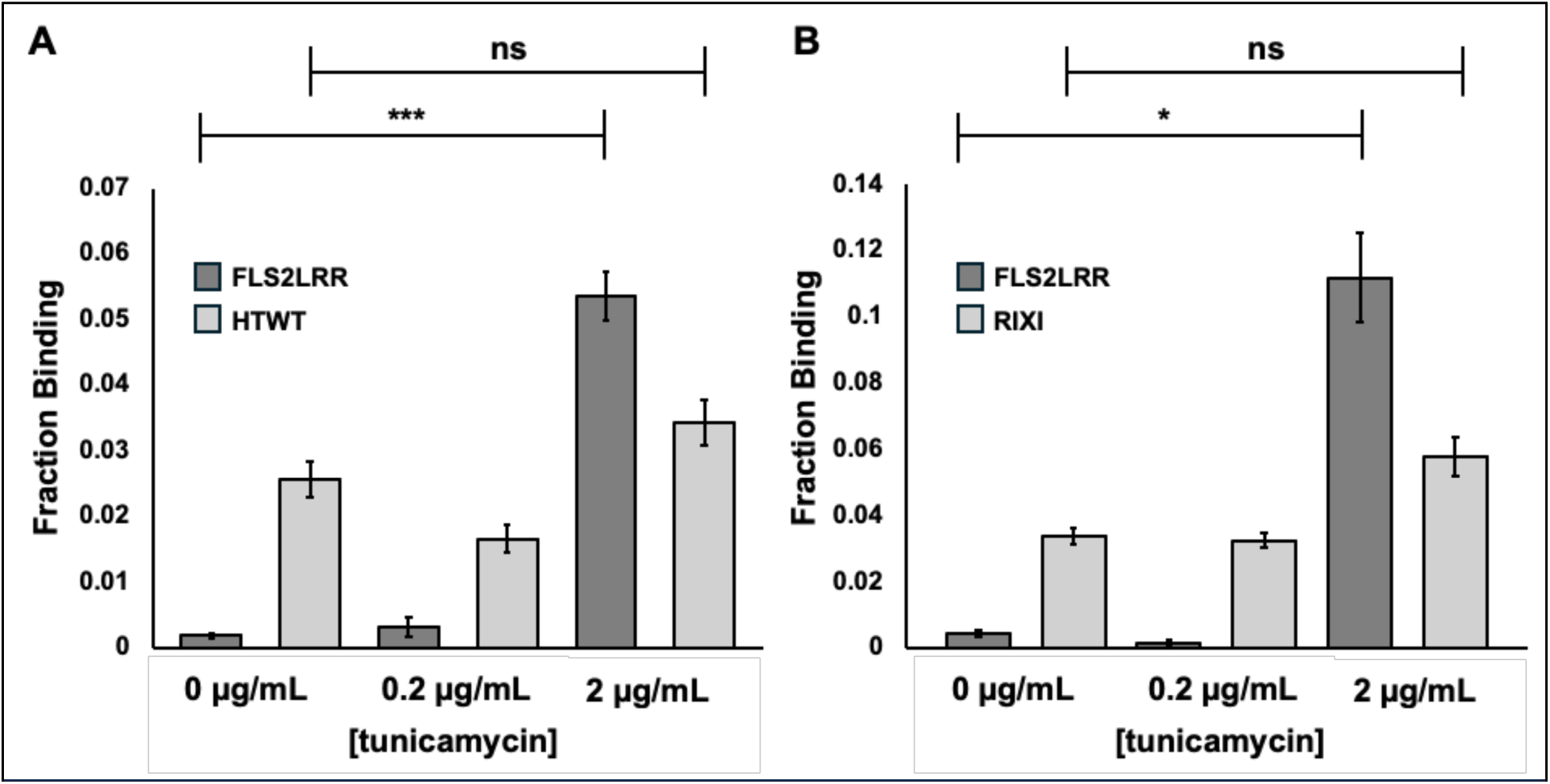
flg22 is selective for FLS2LRR even after tunicamycin treatment during induction. Fraction of FLS2LRR or HTWT (A) or RIXI (B) expressing cells that bind flg22 upon induction with increasing concentrations of tunicamycin. Cytometry data (A,B) shown are the mean and standard error for n=3 biological replicates after background subtraction (see methods) per group (each bar) with Bonferroni Correction factor of 0.025; * < 0.025, ** < 0.0125, *** < 0.00625. HTWT: HaloTag, Engineered *R. rhodochrous* haloalkane dehalogenase (UNIPROT ID: P0A3G2, DHAA_RHORH; PDB ID: 5Y2X). RIXI: Rice GH11 Xylanase Inhibitor (UNIPROT ID: Q7GCM7, XIP_ORYSJ).

These data support the interpretation that FLS2LRR display contributes to the increased flg22-TAMRA-positive population under tunicamycin treatment. However, because the non-FLS2 controls also showed measurable flg22-TAMRA signal and tunicamycin globally affects yeast physiology, these data should be interpreted as evidence for a relative increase in FLS2LRR-associated flg22-TAMRA positivity, rather than definitive proof of receptor-specific flg22 binding. The replicate-to-replicate variation in the subset of 22 possible occupied N-glycan sites observed at 2 µg/mL tunicamycin is consistent with heterogeneous glycosylation and/or stress responses influencing the fraction of receptors capable of binding flg22 (**Figure S8B**), but the current data do not directly identify which FLS2 glycan sites are occupied in each condition. Future site-specific glycosylation analysis and additional biological replication will be needed to define the relationship between glycan occupancy, expression, and ligand-responsive display.

#### Thermal stress and tunicamycin have construct-dependent effects on flg22-TAMRA-positive populations

Tunicamycin treatment also intersects with ER stress and unfolded protein response pathways, which can alter recombinant protein folding, secretion, and cell-surface properties (*58,62,63*). To test whether an additional stress condition could influence FLS2 display, we combined tunicamycin with induction at 37 °C. Heat shock responses in yeast can increase chaperone expression and alter protein homeostasis, which may affect folding or trafficking of difficult-to-express proteins (*64,65*). Because both FLS2LRR and NtoS showed substantial intracellular retention under standard conditions (**Figure 1D**), we asked whether tunicamycin, heat, or their combination would alter the fraction of cells with detectable flg22-TAMRA signal.

We induced FLS2LRR- and NtoS-displaying cells under four conditions: original no stress, tunicamycin alone (2 µg/mL), heat alone (37 °C), and combined tunicamycin plus heat, hereafter referred to as additive stress. For FLS2LRR, the fraction of flg22-TAMRA-positive cells increased from 0.4% under no stress to 3.0% with heat alone, 13.4% with tunicamycin alone, and 13.8% with additive stress (**Figure 3A**). For NtoS, heat alone produced the strongest increase, from 5.4% to 14.7%, whereas tunicamycin alone had a smaller effect (**Figure 3B**). These results indicate that FLS2LRR and NtoS respond differently to stress conditions: FLS2LRR is most strongly affected by tunicamycin-containing conditions, whereas NtoS benefits most from heat alone.

**Figure 3.**
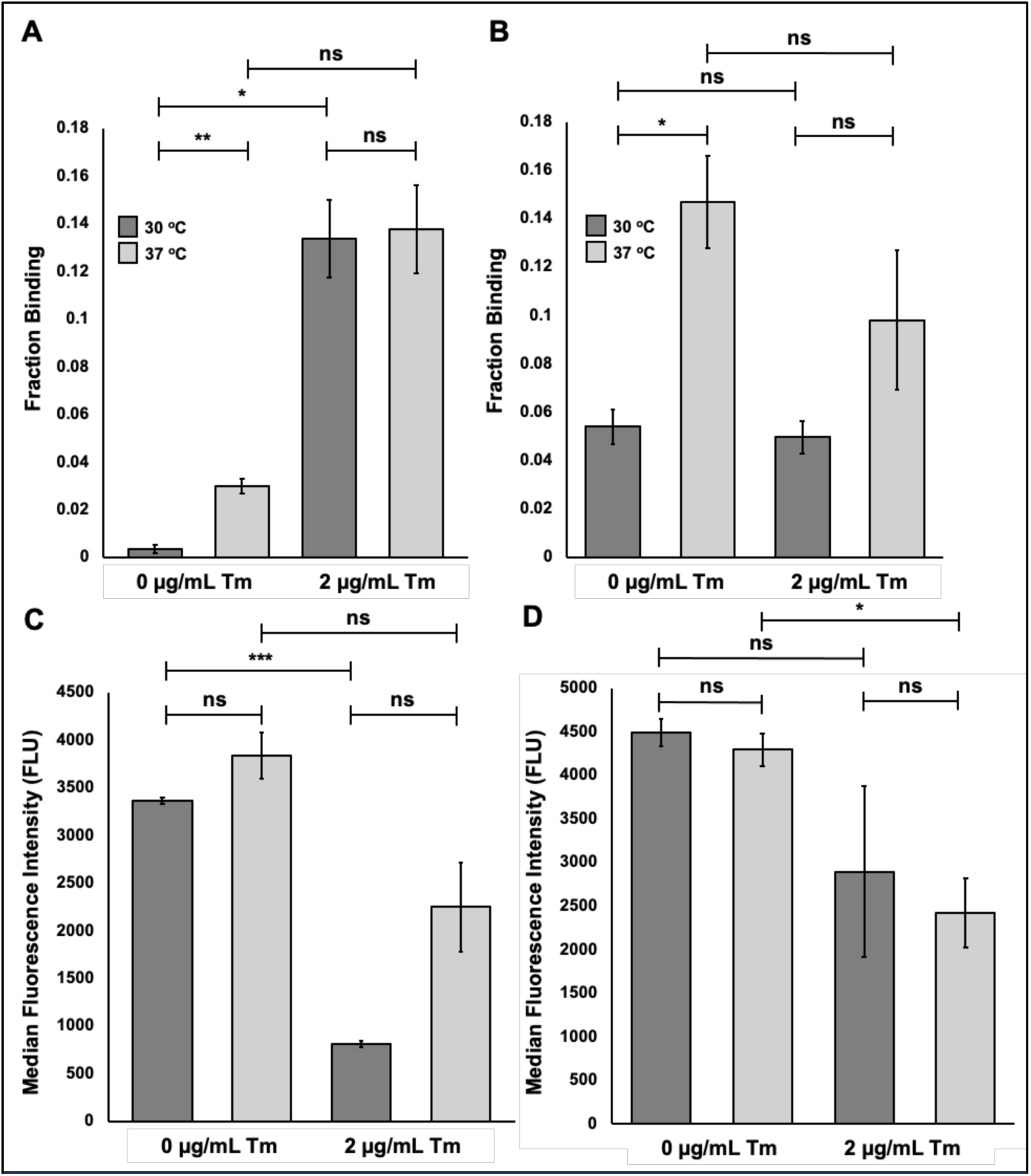
Tunicamycin and thermal stress impact FLS2LRR flg22 binding differently than NtoS. The mean fraction of background subtracted binding/expression positive FLS2LRR (A) and NtoS (B) populations after induction without stress, with tunicamycin, thermal stress, or both. The means of median fluorescence intensity from total surface expression+ve FLS2LRR (C) and NtoS (D) populations after induction with tunicamycin, thermal stress or both. Cytometry data (A-D) shown is the mean and standard error for n=3 biological replicates after background subtraction (see methods) per group (each bar) with Bonferroni Correction factor of 0.0125; * < 0.0125, ** < 0.00313, *** < 0.00078.

These increases in flg22-TAMRA-positive cells occurred alongside reduced expression. Median fluorescence intensity for FLS2LRR expression decreased approximately 4.2-fold upon tunicamycin treatment (**Figure 3C**), and NtoS expression decreased approximately 1.9-fold under additive stress (**Figure 3D).** Thus, stress conditions increased the fraction of cells scored as flg22-TAMRA-positive while reducing overall surface expression, consistent with a tradeoff between expression level and the emergence of a ligand-responsive or ligand-associated subpopulation. Density plots further showed that NtoS double-positive events were detectable under fewer stress conditions, whereas FLS2LRR showed the clearest double-positive population under additive stress (**Supplementary Information 5a-b**).

Although NtoS showed strong flg22-TAMRA positivity under heat stress, complete removal of all predicted N-glycosylation motifs reduced surface expression and may be less useful as an engineering scaffold if glycosylation is required for productive expression or function in planta. Häweker et al showed that complete de-glycosylation led to intracellular retention and heavily reduced trafficking after introducing high concentrations of tunicamycin (10 ug/mL) in planta (*38*). Therefore, we focused subsequent experiments on FLS2LRR and partially glycosylated FLS2LRR-derived variants, with the goal of identifying conditions or glycan-site substitutions that increase the flg22-TAMRA-positive population while preserving more of the native glycosylation framework necessary for trafficking in planta.

#### Concentration-dependent flg22-TAMRA signal reveals assay background and limits quantitative affinity interpretation

To determine whether the flg22-TAMRA-positive populations reflected saturable receptor-ligand binding, we measured concentration-dependent flg22-TAMRA signal across a range of ligand concentrations. Chinchilla et al. reported IC50 values of 5–10 nM for competition of radiolabeled flg22 by unlabeled flg22 in plant membrane preparations (*66*). In contrast, our flow cytometry experiments required substantially higher flg22-TAMRA concentrations, suggesting that binding in the yeast-display format, if present, was much weaker and/or confounded by background signal. Because high ligand concentrations can increase weak or nonspecific interactions in yeast surface display assays (*67*), we included the FLS2LRR-S390K variant as a negative-control FLS2 construct based on prior evidence that this mutation disrupts flg22 binding (*68*).

We induced FLS2LRR, NtoS, and FLS2LRR-S390K under no-stress or additive-stress conditions and incubated cells with 0.1, 1, 3, 10, 50, and 100 µM flg22-TAMRA before flow cytometry. Under no-stress conditions, none of the constructs showed a strong concentration-dependent increase in MFI. Under additive stress, however, all three constructs showed increasing MFI with increasing flg22-TAMRA concentration (**Figure 4**). Importantly, none of the curves reached a clear plateau, even at 100 µM flg22-TAMRA. Moreover, the S390K negative-control construct also produced a concentration-dependent signal under additive stress.

**Figure 4.**
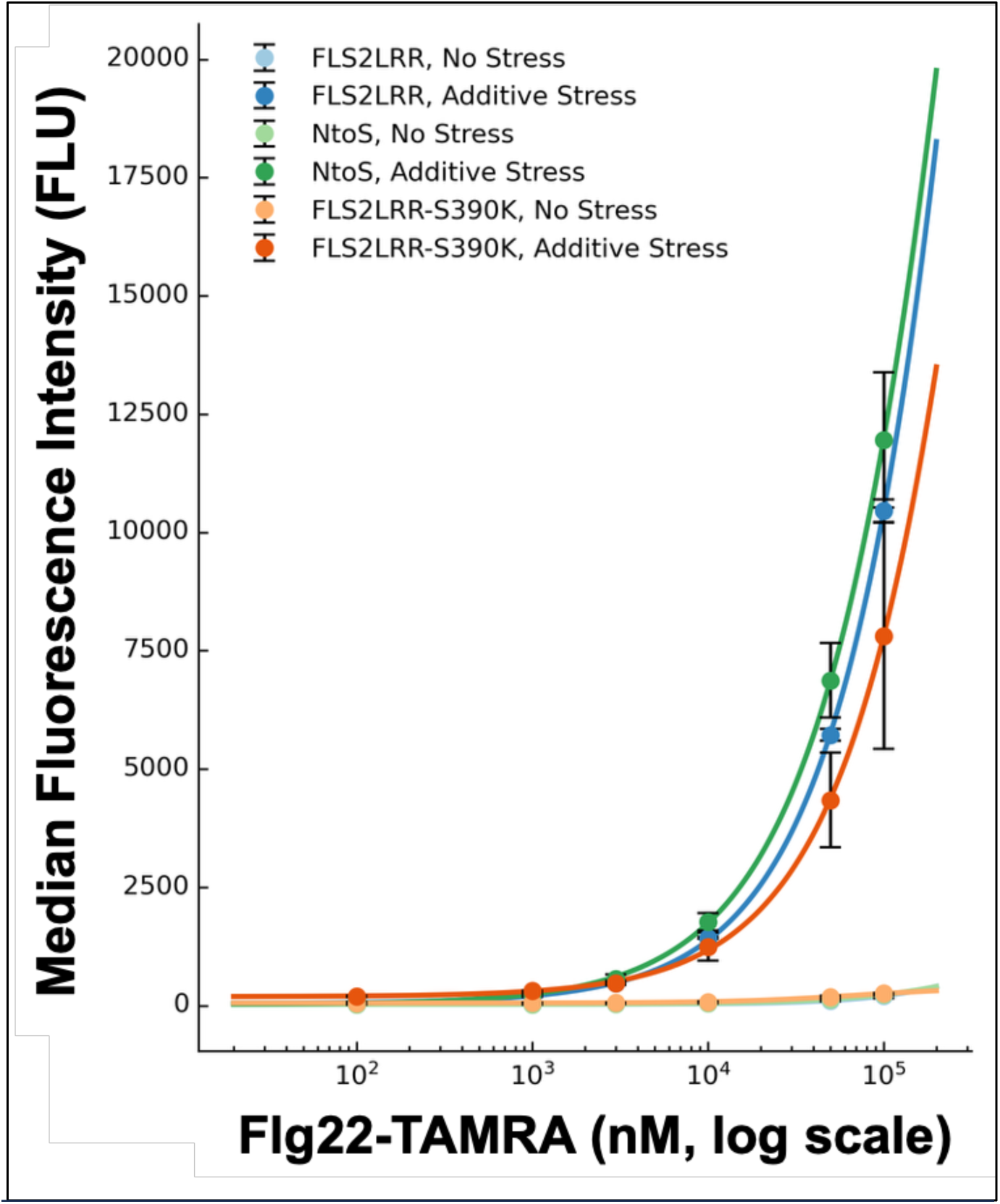
On the surface of yeast, FLS2LRR and NtoS show weak affinity for flg22-TAMRA. The binding affinity of FLS2LRR, NtoS, and FLS2LRR-S390K after no stress or additive stress induction. Surface display cells were treated with 0.1, 1, 3, 10, 50, and 100 µM flg22-TAMRA and median fluorescence intensity was obtained to determine binding affinity. Affinity (KD) was estimated by fitting flg22-TAMRA concentration and median fluorescence intensity with a nonlinear least-squares regression curve showing mean and standard error for n=3 biological replicates per group (each point).

Because the curves did not saturate and the S390K control retained substantial concentration-dependent signal, fitted values from these data should be interpreted only as apparent signal-response parameters, not true equilibrium dissociation constants. Fitting MFI as a proxy for receptor occupancy produced apparent values of approximately 34 µM for FLS2LRR, 43 µM for NtoS, and 73 µM for FLS2LRR-S390K. These values are several orders of magnitude weaker than the nanomolar flg22-binding values reported in plant membrane assays and likely reflect a combination of weak FLS2-associated binding, stress-induced background, assay-format differences, and nonspecific flg22-TAMRA interactions at high ligand concentrations.

To reduce background, we subtracted TAMRA signal associated with FLS2-negative cells from the FLS2-positive population (**Supporting Information 4**). However, substantial background remained, particularly above 1 µM ligand. Therefore, the flow cytometry titration data do not support quantitative affinity determination for yeast-displayed FLS2 under these conditions. Instead, they reveal an important limitation of the assay: additive stress enables concentration-dependent flg22-TAMRA signal, but this signal is not sufficiently specific or saturable to confidently distinguish true FLS2-flg22 binding from false-positive or nonspecific events.

These results refine the interpretation of the earlier flow cytometry experiments. Tunicamycin and thermal stress increase flg22-TAMRA-positive populations, especially for FLS2LRR, but flow cytometry alone cannot establish that these events represent specific receptor-ligand binding. Essentially, these experiments revealed that the relatively low binding affinity between yeast displayed FLS2 and flg22-TAMRA was incompatible with flow cytometry analysis given the known limit of detecting weak (mid-micromolar affinity) interactions (*69,70*).Consequently, additional orthogonal approaches are required to test whether flg22 can selectively enrich FLS2-displaying cells and to distinguish true binding populations from stress-associated background.

#### Magnetic bead enrichment provides orthogonal support for flg22-associated enrichment of FLS2LRR-displaying cells

Because flow cytometry titrations revealed substantial background and did not support quantitative affinity determination, we next used magnetic bead enrichment as an orthogonal assay to test whether flg22 could preferentially recover FLS2LRR-displaying cells from a large excess of non-FLS2-displaying cells. We reasoned that bead-based enrichment could detect weak or low-frequency ligand-associated populations that are difficult to resolve by single-cell flow cytometry, particularly when selections are performed under high ligand-density conditions (*70,71*).

We adapted a previously reported yeast display selection workflow (*70*) using streptavidin-coated magnetic beads that were either unmodified, coated with biotin-IgG, or coated with biotin-flg22 (**Figure S10**). Unmodified beads and biotin-IgG beads were used for negative selections to reduce recovery of cells that bind beads or non-target proteins. Biotin-flg22 beads were then used for positive selection. FLS2LRR-expressing cells and RIXI-expressing cells were induced under either no-stress or additive-stress conditions, mixed at a 1:1000 FLS2LRR:RIXI ratio, and subjected to sequential negative and positive selections. Bead-bound cells were recovered after each round, and clones were genotyped to estimate the fraction of FLS2LRR and RIXI cells in the enriched populations (**Figure S10-S11**).

After the first round of selection, all screened clones from each induction condition were RIXI-positive, indicating that one round of biotin-flg22 bead selection was insufficient to recover detectable FLS2LRR clones from the initial 1:1000 mixture (**Figure S11A**). After a second round of selection, however, FLS2LRR clones were detected in nearly all replicate selection groups. In two independent experiments, additive-stress-induced FLS2LRR cells showed enrichment factors of approximately 330- to 830-fold relative to the starting mixture (**Figure S11B; Supplementary Information 1**). These results indicate that repeated biotin-flg22 bead selection can enrich FLS2LRR-displaying cells from a large excess of RIXI-displaying cells.

These enrichment data provide orthogonal support for the conclusion that at least a subset of yeast-displayed FLS2LRR cells can be preferentially recovered using flg22-coated beads. This finding is consistent with the flow cytometry results showing increased flg22-TAMRA-positive populations under stress conditions, but it should not be interpreted as definitive proof of receptor-level binding specificity. The assay compares FLS2LRR to RIXI, an unrelated non-FLS2 displayed protein, but does not yet include key FLS2-specific negative controls such as FLS2LRR-S390K or competition with free flg22. Therefore, the bead-enrichment results support flg22-associated enrichment of FLS2LRR-displaying cells, while further controls will be needed to determine whether enrichment reflects specific FLS2-flg22 recognition, avidity effects, altered surface properties under stress, or other display-dependent factors.

Overall, magnetic bead enrichment strengthens the evidence that the stress-optimized display conditions produce an enrichment-compatible FLS2LRR population. However, the current data do not yet establish a quantitative affinity-screening platform or fully resolve the molecular basis of the interaction. Future enrichment experiments that include FLS2LRR-S390K, relevant reduced site-specific glycan state variants, excess soluble flg22 competition, scrambled peptide controls, and replicate-level sequencing would more directly test receptor-specific flg22 recognition.

### Structure-guided glycan-site variant analysis identifies that N388S is associated with a promising display phenotype

Tunicamycin-based partial glycosylation increased the fraction of flg22-TAMRA-positive FLS2LRR cells, but tunicamycin globally perturbs N-glycosylation, ER stress, cell physiology, and surface-display behavior. Therefore, the increased flg22-TAMRA signal could not be assigned to specific FLS2 glycosylation sites. To more directly test whether individual predicted N-glycosylation sites influence the flg22-TAMRA-positive display phenotype, we designed structure-guided glycan-site variants.

We modeled yeast high-mannose glycans onto the FLS2 ectodomain crystal structure bound to flg22 (PDB ID 4MN8) (*16*) using the GlycoShape Re-glyco tool (*72*) (**Figure 5A**). The modeled structure was aligned to the original crystal structure in PyMOL (*73*), focusing on FLS2 chain A and flg22 chain C. We then examined the spatial relationship between modeled glycans and the flg22-binding interface, as well as regions where multiple modeled glycans clustered on the FLS2 ectodomain. From this analysis, we identified two classes of candidate sites: proximal sites, where modeled high-mannose glycans approached the flg22 peptide within 5 Å and could plausibly affect ligand accessibility, and aggregated sites, where nearby glycans appeared to crowd local regions of the ectodomain. This analysis identified three proximal sites, N347, N371, and N388, and two aggregated sites, N631 and N704.

**Figure 5.**
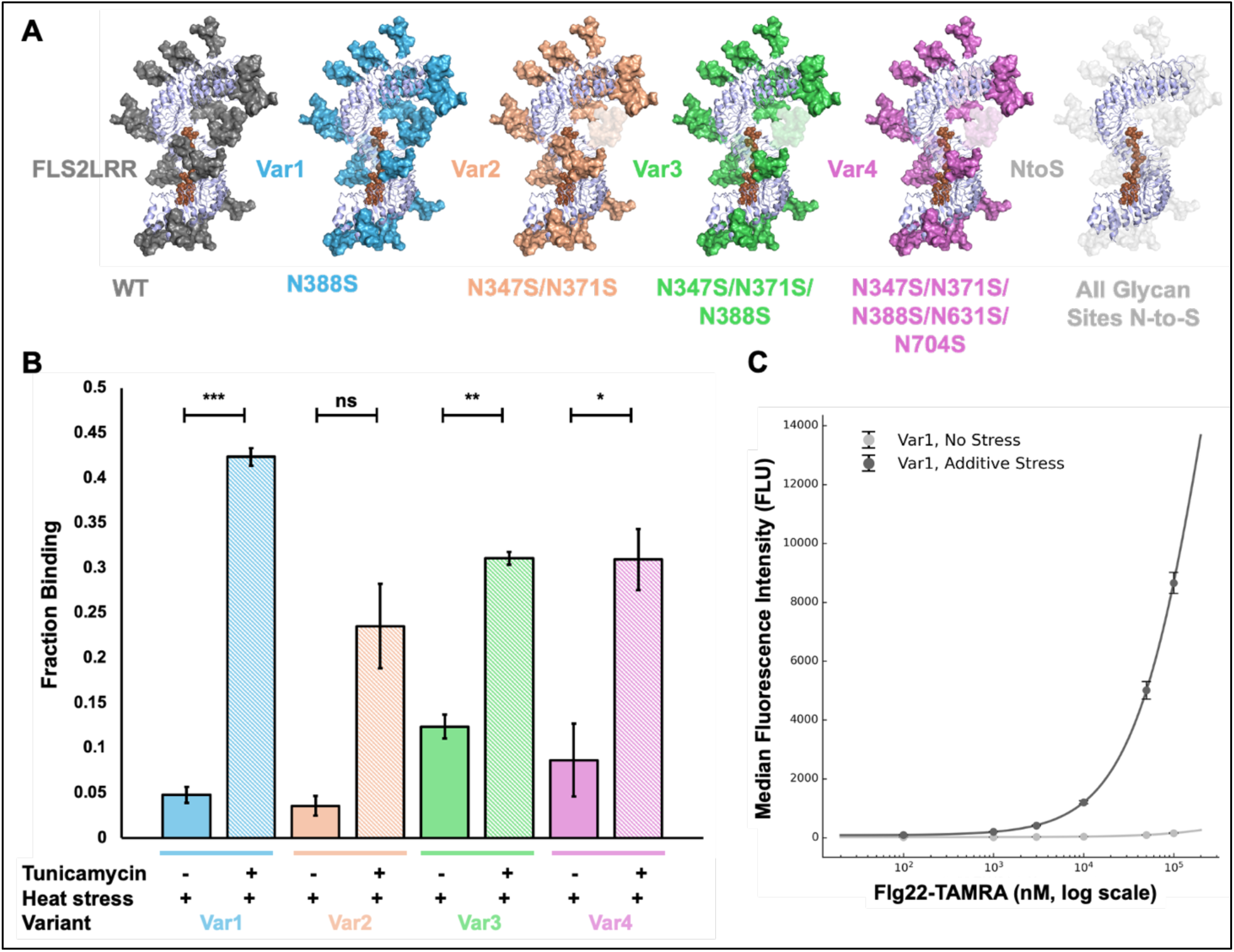
Glycosylation site variants including Var1 (N388S) typically outperform FLS2LRR after additive stress induction. (A) GlycoShape Re-Glyco modeled FLS2 ectodomain crystal structure (PDB ID 4MN8) with high-mannose glycans (GlyTouCan ID G92042VQ). Transparent glycans are modified for that specific FLS2LRR glycan variant. (B) The fraction binding/expression positive populations for four select site NtoS variants after induction of protein expression at either 37 oC (filled bars) or 37 oC with 2 µg/mL tunicamycin (striped bars). (C) The binding affinity of Var1 after additive and no stress induction using the same concentrations and curve fitting parameters as described previously. Cytometry data shown in (B) is the mean and standard error for n=3 biological replicates after background subtraction (see methods) per group (each bar) with Bonferroni Correction factor of 0.0125 (4 comparisons): * < 0.0125; ** < 0.00313; *** < 0.00063.

Based on these positions, we designed four FLS2LRR-derived glycan-site variants: Var1, N388S; Var2, N347S/N371S; Var3, N347S/N371S/N388S; and Var4, N347S/N371S/N388S/N631S/N704S (**Figure 5A**). Each variant was expressed on the yeast surface as an Aga2 fusion, and flg22-TAMRA positivity and surface expression were evaluated by flow cytometry under heat stress alone (37 °C) or additive stress (37 °C plus 2 µg/mL tunicamycin) (**Figure 5B**). Because Var1 showed the clearest additive-stress-associated increase in the double-positive population, we also performed a flg22-TAMRA titration for this variant after no stress and additive-stress induction (**Figure 5C**).

Under heat stress alone, Var3 and the NtoS construct showed the highest fractions of flg22-TAMRA-positive cells, approximately 12.4% and 11.5%, respectively, whereas Var1 and Var2 showed lower fractions, approximately 4.8% and 3.6%, respectively (**Figure 5B**). Raw density plots showed that Var3 produced a double-positive population similar to NtoS under heat stress, suggesting that combined removal of N347, N371, and N388 can increase the flg22-TAMRA-positive phenotype under this condition (**Supporting Information 5c-e**). However, because the NtoS construct also showed this phenotype and because prior titration experiments revealed substantial assay background, these data should be interpreted as changes in the flow-cytometry display phenotype rather than definitive evidence of increased receptor-specific binding.

Under additive stress, flg22-TAMRA-positive events increased for most glycan-site variants, although the increase for Var2 was not statistically significant relative to heat stress alone (**Figure 5B**). Var1 showed a robust and statistically significant increase in binding-positive/expression-positive events under additive stress compared with heat stress alone. In raw density plots, Var1 also showed a larger and more compact double-positive population than Var2 or Var3 under additive stress (**Supporting Information 5c-e**). Additionally, two-factor ANOVA for the significance of flg22-TAMRA positive change across treatments, variants, and between combined treatment and variant groups showed a significant difference across all comparisons (p-value = 1.25x10^-14^, 0.000132, 0.000694, respectively; **Supplementary Table 1**). This indicates that there is a strong effect of treatment, the variants differ overall, and treatment effect depends on the variant, respectively. Overall, this supports a claim where the effect of treatment is not the same for every variant such as how Var1 positive binding phenotype increases substantially compared to other variants between the no stress and additive stress treatment groups. These results identify N388S as a promising variant that increases the flg22-TAMRA-positive display phenotype while avoiding the broader glycan-site removal used in Var3, Var4, or NtoS.

We next asked whether the improved Var1 phenotype reflected saturable flg22-TAMRA binding. However, Var1 did not reach a clear binding plateau over the flg22-TAMRA concentration range compatible with reliable flow cytometry analysis (**Figure 5C**). Therefore, the Var1 titration does not support assignment of a quantitative KD or definitive receptor-level affinity. Instead, the data show that N388S increases the fraction of cells detected as flg22-TAMRA-positive under additive stress, while the underlying molecular basis of this increase remains unresolved.

Collectively, these data identify N388 as a candidate glycosylation site that influences the yeast-display phenotype of FLS2LRR. Because N388 is proximal to the flg22-binding interface in the glycan-modeled structure, one plausible model is that a yeast high-mannose glycan at this site could reduce ligand accessibility. However, the current data do not directly demonstrate glycan occupancy at N388, physical occlusion of flg22, or a specific rescue of receptor-ligand binding upon N388S substitution. N388S could also improve folding, trafficking, surface orientation, glycan heterogeneity, or background-corrected cytometry behavior. Thus, the most evidence-supported conclusion is that N388S increases the flg22-TAMRA-positive/expression-positive population under additive stress and represents a promising candidate for follow-up studies using more direct binding, competition, enrichment, and glycosylation-site occupancy assays.

## Conclusion

We set out to evaluate whether yeast surface display could support functional presentation of the plant pattern recognition receptor FLS2, motivated by the need for higher-throughput approaches to engineer receptors with altered or broadened flagellin recognition. Our results show that the A. thaliana FLS2 ectodomain can be expressed on the surface of S. cerevisiae as an Aga2 fusion, but that standard yeast display conditions produce little or no detectable flg22 binding. This disconnect between surface expression and ligand detection motivated a systematic analysis of how yeast N-linked glycosylation and stress-responsive folding pathways influence the FLS2 display phenotype.

Several conclusions are strongly supported by the current data. First, yeast-displayed FLS2LRR is extensively modified by Endo H-sensitive glycans, as shown by the large molecular-weight shift after Endo H treatment under denaturing conditions. Second, complete removal of all predicted N-glycosylation motifs reduces surface expression and increases intracellular retention, suggesting that some degree of N-glycosylation supports productive folding and/or trafficking of FLS2 in yeast. Together, these findings reveal a central tradeoff: yeast-type glycosylation may interfere with detectable ligand-associated signal, but complete glycan-site removal compromises efficient surface display.

Tunicamycin treatment and thermal stress increased the fraction of cells scored as flg22-TAMRA-positive, particularly for FLS2LRR under tunicamycin-containing conditions and for NtoS under heat stress. However, these effects occurred alongside reduced expression and substantial assay background. Moreover, concentration-dependent flg22-TAMRA signal did not saturate for FLS2LRR, NtoS, or the S390K negative-control variant, even at high ligand concentrations. Therefore, these flow cytometry data should be interpreted as changes in a flg22-TAMRA-positive display phenotype rather than as definitive evidence of restored native-like binding or quantitative receptor affinity.

Magnetic bead enrichment provided orthogonal support that biotin-flg22-coated beads can enrich FLS2LRR-displaying cells from a 1:1000 mixture with RIXI-displaying cells after two rounds of selection. This result suggests that at least a subset of yeast-displayed FLS2LRR cells can be preferentially recovered using flg22-coated beads. However, because the current enrichment assay does not yet include FLS2-specific negative controls such as S390K or soluble flg22 competition, it does not fully resolve receptor-level binding specificity.

Finally, structure-guided glycan-site analysis identified N388S as a promising variant that increases the flg22-TAMRA-positive/expression-positive population under additive stress. Because N388 is proximal to the flg22-binding interface in the glycan-modeled structure, it is a plausible site where yeast high-mannose glycosylation could influence ligand accessibility. However, the current data do not directly demonstrate glycan occupancy at N388, physical occlusion of flg22, or a specific mechanistic rescue of FLS2-flg22 binding. Thus, N388 should be described as a candidate site influencing the yeast-display phenotype, not yet as a proven determinant of ligand access.

Overall, this work does not yet establish a quantitative affinity-screening platform for FLS2 engineering. Instead, it defines key barriers and enabling conditions for yeast display of a large, glycosylated plant immune receptor. The data support a model in which yeast glycosylation, ER stress, and selected glycan-site substitutions shape an enrichment-compatible flg22-responsive phenotype. Future studies incorporating soluble ligand competition, FLS2-specific negative controls, site-specific glycosylation analysis, and sequencing-based enrichment will be needed to distinguish true receptor-ligand binding from stress-associated background and to convert this system into a reliable platform for engineering FLS2 and other difficult plant glycoproteins.

## Materials and Methods

### Materials and Media preparation

We purchased yeast extract, peptone, and dextrose from Fisher for YPD growth media preparation. We purchased SD/-trp packets (clonetech) for SD/-trp selection media preparation. We purchased sodium phosphate dibasic, sodium phosphate monobasic, galactose from Sigma-Aldrich; dextrose and casamino acids from Fisher; and yeast nitrogen base from DOT Scientific for SG-CAA induction media preparation. We purchased LB broth (Miller) for bacterial growth media. For growth or selection agar plate preparation, we purchased agar from fisher. Agar plates were prepared in 500 mL batches including reagents specified for growth or selection media and 8 g agar, then autoclaved, cooled, and poured onto 100x15mm petri dishes from Fisher (∼20-25 mL media + agar per plate). For short-term storage (1-2 months), ampicillin (Sigma-Aldrich) was added to cooled media for a final concentration of 1 mg/mL ampicillin. Bacterial media and agar plates always contained 1 mg/mL ampicillin for plasmid selection.

For western blot wash buffer, 500 mL of 1X TBS was prepared using 10% 10X TBS (24 g Tris, 88 g NaCl). For flow cytometry working buffer (washing, staining), 500 mL of 1X PBS + 0.1% (w/v) BSA was prepared using 10% 10X PBS w/o calcium, w/o magnesium (VWR) and 0.5g bovine serum albumin (Fisher), then sterile filtered.

### Yeast and Bacterial Strains and Plasmids

All surface display constructs were designed in-house, except the plasmid pCT80-FnLoopHp, which is routinely used for our YSD applications. Insert sequences were codon optimized (*S. cerevisiae*) and cloned into pCT80-FnLoopHp by GenScript, except FLS2LRR. pCT80-FnLoopHp is a modified version of the pCTcon2 (*74*) surface display shuttle vector that was used to design FLS2LRR, NtoS, 4 NtoS glycan variants, RIXI, and HaloTag yeast display constructs (**Figure S2, Supplementary Information 2,3**). Briefly, *S. cerevisiae* codon-optimized recombinant FLS2LRR insert sequence with homologous overlapping ends (**Supplementary Table 2**) was inserted via homologous recombination or Gibson cloning at NheI and BamHI restriction enzyme cut sites in the GAL1 open reading frame of restriction enzyme linearized vector. All constructs were transformed into the *EBY100* yeast strain (genotype MATa AGA1::GAL1-AGA1::URA3 ura3-52 trp1 leu2-delta200 his3-delta200 pep4::HIS3 prbd1.6R can1 GAL). For plasmid propagation, NEB5a (NEB) *E. coli* strain was used.

### Yeast outgrowth, storage, transformation, selection, and induction of protein expression

WT *EBY100* yeast passage was performed in YPD media and all stocks were stored at −80 °C in 25% Glycerol. Yeast transformation was performed using the Frozen-EZ Yeast Transformation II Kit (Zymo) or electroporation was performed to acquire recombinant yeast clones by homologous recombination of linearized display vector with recombinant inserts containing homologous overlapping ends (*75*). Transformants were plated on SD/-trp plates for selection. Yeast transformants were grown in SD/-trp selective media at 30 °C/225 rpm/overnight and induced in SG-CAA induction media at 30 °C/225 rpm/overnight. To confirm successful transformation, single colony clones were picked and outgrown in selective media for DNA extraction using the Zymoprep Yeast Plasmid Miniprep II kit (Zymo). Then, genomic extract was transformed by heat shock into NEB5a chemically competent *E. coli* (following High Efficiency Transformation Protocol from NEB) for plasmid propagation and extraction (QIAprep Spin Miniprep Kit, QIAGEN). Plasmid concentration was determined using a UV/Vis Spectrophotometer (NanoDrop One, Fisher). DNA plasmids were submitted to Plasmidsaurus for High Copy Whole Plasmid Sequencing service to confirm successful yeast transformants.

For induction under stress conditions, after outgrowth in SD/-trp, yeast were resuspended in SG media with or without tunicamycin (Sigma-Aldrich) at a starting OD of 0.2- 0.5 and incubated at either 30 or 37 °C/225 rpm/overnight. Due to growth disruption and cell death caused by tunicamycin treatment, it is recommended to grow in a larger volume of induction media for sufficient cell quantities in downstream experiments (e.g. 2-3x induction cultures).

### FLS2 Computational Modeling and Stability Calculation

**Figure S7** summarizes the workflow for identifying amino acids to substitute putative N-glycan asparagines by AlphaFold2 (AF2) modeling of de-glycosylated FLS2 ectodomains and metrics/tools for evaluating prediction performance, modeling atomic deviation, and modeling ΔEnergy. Briefly, AF2 provides an interface predicted template modelling (ipTM) score, which measures how well AlphaFold2-Multimer predicts the overall structural complex (*47*). Using the Rosetta molecular modeling software (*76*), modeled structures are relaxed (Fast Relax) and total_score is calculated using the Rosetta scoring function (REF15), which is compared to the AF2-predicted WT structure total_score as described previously (*48,76*). Root mean-squared deviation (RMSD, Å) is obtained after PyMOL (*73*) structural alignment of relaxed modeled structures to the crystal structure (PDB ID 4MN8, only FLS2 ectodomain (chain A) and flg22 (chain C)). Evaluation metrics (ipTM, RMSD, total_score) were tabulated for WT and substituted modeled structures to determine which amino acid change at all putative N-glycan sites (substituted structures) resulted in the fewest deviations from values for the WT modeled structure (**Figure S7**). This FLS2LRR de-glycosylated variant clone was established in yeast as described in *<u>Yeast outgrowth, storage, transformation, selection, and induction of protein expression</u>*.

For choosing specific-site NtoS variants, FLS2 ectodomain crystal structure (PDB ID 4MN8) was modelled with yeast-type high mannose glycans using GlycoShape (*72*). Prior to choosing N-glycan sites for removal, the modeled structure was aligned to the original crystal structure (only FLS2 ectodomain (chain A) and flg22 (chain C)) in PyMOL. To determine N-glycan sites chosen for removal, we considered the proximity within 5 Å of modelled high-mannose glycan atoms to flg22 (proximal sites) and locations where GlycoShape predicted glycan aggregation (aggregated sites) using PyMOL and commands described previously (*48*). Based on these constraints we chose 3 proximal glycan sites (N388, N347, N371) and 2 aggregated glycan sites (N631, N704S). Four variant constructs were designed using combinations of these identified sites – **Var1**:N388S, **Var2**:N347S/N371S**, Var3**:N388S/N347S/N371S, **Var4**:N388S/N347S/N371S/N631S/N704S. FLS2LRR variant clones were established in yeast as described in *<u>Yeast outgrowth, storage, transformation, selection, and induction of protein expression.</u>*

### Protein Extraction, SDS-PAGE, and Western Blot Assays

To obtain protein extracts containing surface displayed protein from EBY100 transformant clones – after overnight induction of protein expression – cell cultures were treated with 2 mM DTT (GOLDBIO) for 45 minutes at 30 °C/225 rpm. DTT disrupts disulfide bridges formed between Aga1 and Aga2 that allow for yeast surface display (*77*). For obtaining cytosolic extracts, after surface extraction with DTT, we performed horn-type sonication of a high cell density mixture as described previously (*78*). Briefly, sonication per sample was performed 20 s on, 30 s off, for 10 minutes. After DTT treatment and sonication, cells were pelleted at 4000 xg/ 10 minutes and supernatant was collected for Amicon Ultra 15-30K concentration. Concentrated supernatant protein density was determined by Bradford Assay (Quick Start Bradford Protein Assay, Bio-Rad) against freshly prepared BSA standard dilution panel using 595 nm setting on a spectrophotometer. After determining protein concentration, samples were snap frozen and stored at −80 °C as 50-100 uL 0.5-2 mg/mL aliquots in 10% glycerol and 1X Protease Inhibitor solution (Halt Protease Inhibitor Cocktail (100X), Fisher).

For protein extracts treated with endoglycosidase H (Endo H, NEB) or PNGase F (NEB), 10-20 µg concentrated supernatant protein was processed following the manufacturer recommended instructions for denaturing (PNGase F only) and non-denaturing Endo H treatment. Briefly, non-denaturing reaction samples were incubated with Glyco 3 Buffer and Endo H at 37 °C overnight, prior to SDS-PAGE and Western Blot. Non-denaturing incubations did not include a the initial incubation with denaturing buffer.

To prepare samples for SDS-PAGE, 10-20 µg of concentrated supernatant protein was prepared with 2.5 uL LDS (4X LDS Sample Buffer, GenScript) and 1 uL 500 mM DTT, then denatured at 100 °C/ 10 minutes. Pre-made 4-12% Bis-Tris gel (SurePAGE, Bis-Tris, 10x8, 4-12%, 12 well, GenScript) was prepared in polyacrylamide gel electrophoresis box with Tris-MOPS-SDS Running Buffer (GenScript). Samples and PageRuler Prestained protein ladder (Fisher) were loaded onto the gel so that sample protein quantity was equivalent for each sample regardless of prior treatments (e.g. loading 20 uL endo-H treated sample vs 10 uL un-treated sample both contain 10 µg of concentrated supernatant protein from the same initial batch extraction).

Electrophoresis was conducted at 200V/40min. After SDS-PAGE, gel was prepared for wet transfer using the sandwich method (*79*) in 0.1% SDS, 1X Transfer (Towbin) Buffer (10% v/v 10X Transfer Buffer (Fisher), 20% v/v Methanol (Sigma-Aldrich)). Briefly, sandwich was prepared for gel to activated PVDF membrane transfer and loaded into Criterion Blotter (Bio-Rad) for wet transfer at 4 °C/ 90 V/ 60 minutes. After wet transfer, membrane was carefully removed and washed twice with 1X TBS, then blocked with 1X TBS + 3% (w/v) BSA for 60 minutes/room temperature (RT)/slow shaking. After blocking, membrane was washed once with 1X TBS, then twice with 1X TBS-T (0.1% v/v Tween-20), before incubation in primary antibody solution (1:1000 anti-HA-biotin (3F10, Millipore Sigma), 1X TBS + 3% BSA) overnight/4 °C/slow shaking. After primary antibody incubation, membrane was washed twice with 1X TBS-T before incubation in secondary antibody solution (1:10000 streptavidin-AlexaFluor647 (Fisher), 1X TBS + 3% BSA) for 60 minutes/RT/slow shaking. After secondary antibody incubation, membrane was washed four times with 1X TBS-T and detected with laser compatible for AlexaFluor647 detection using a ChemiDoc Imager (BioRad).

### Flow Cytometry Assays

To prepare samples for flow cytometry analysis, induced cells optical density was measured using a spectrophotometer set to measure OD600. For each sample replicate, 4x10^6^ were transferred to microfuge tubes and washed with 1X PBS + 0.1% BSA. After washing, each sample was incubated with primary antibody at a final concentration of 5 µg/mL in 1X PBS + 0.1% BSA for 20 minutes at room temperature. After primary antibody incubation, samples were washed and resuspended in secondary antibody solution at a final concentration of 10 µg/mL in 1X PBS + 0.1% BSA for 20 minutes at room temperature. Finally, after secondary antibody incubation, samples were washed and resuspended in 1-10 µM TAMRA-flg22 (BIOMATIK) in 1X PBS + 0.1% BSA solution with concentration consistent between experiments unless otherwise specified. Incubation with TAMRA-flg22 was conducted for 20 minutes at room temperature. After final incubation, samples were washed once and resuspended in 100 or 500 uL 1X PBS + 0.1% BSA for flow cytometry analysis using either the Accuri C6 (BD Biosciences) or Attune Cytpix (Fisher). The median fluorescence intensity from flow cytometry measurements for FLS2LRR and variant populations was used to calculate the binding affinity (K_D_) as described previously (*80,81*). FCS Express 7 (De Novo Software) was used for statistical analyses, data extraction, and figure preparation of cytometry data.

### Magnetic Bead Enrichment of FLS2LRR-displaying cells

**Figure S9** diagrams the workflow for the first round of enrichment, subsequent rounds of enrichment, and enrichment analysis using a modified approach described in Ackerman et al. 2009 (*70*). Briefly, modifications to this approach include: using Dynabeads Biotin Binder (Fisher) magnetic streptavidin-coated beads. Beads are coated with negative (biotin conjugated goat-IgG, Rockland) and positive (biotin-flg22, GenScript) selection agent prior to incubation with cells (direct approach). Selections are conducted for 1.5 hrs/4 °C/rotating before bead binding cells are transferred for enrichment analysis. Prior to induction of positive selection cells before subsequent rounds, beads from previous round are removed. To wash, remove unbound cells, and remove previous round beads, the bead/cell mixture was placed on a DynaMag-2 (Fisher) magnetic tube rack for 2-5 minutes before removal of suspension without disrupting beads. Bead/cell mixtures are washed three times between selections using 1X PBS + 0.1% BSA.

### Statistical Analysis and Reproducibility

We determined the statistical significance with two-tailed T-test with unequal variance for all cytometry data and used Bonferroni correction (corrected α provided in figure descriptions) when >1 comparisons were made. All collected data were used for the calculation of means and standard error. The fraction binding for cytometry data bar plots represents the mean of n=3 biological replicates after background subtraction performed as described in **Supplementary Information 4**.

## Abbreviations

FLS2: FLAGELLIN SENSING 2
PRR: Pattern Recognition Receptor
ERQC: Endoplasmic Reticulum Quality Control
LLO: Lipid-linked Oligosaccharide
OST: Oligosaccharyltransferase
DTT: Dithiothreitol
UPR: Unfolded Protein Response
TAMRA: Tetramethylrhodamine
MFI: Median Fluorescence Intensity
YSD: Yeast Surface Display

## Author Information

### Author Contributions

B.D. and D.R.W. contributed equally to the conception, design, and interpretation of the study.

B.D. and S.S. contributed to the execution of experiments. All authors contributed to writing review, and approval of the final manuscript.

### Notes

The authors declare no competing financial interest.

## Acknowledgement

We thank Benjamin Orlando for his contribution to conceptualization of the FLS2LRR specific site glycosylation variant panel. We are thankful to Jens Schmidt for suggestions on analyzing flow cytometry data. We thank Eric Patterson and Michael Feig for critical assessment of findings throughout. We are thankful to the Michigan State University Flow Cytometry Core for training and super-user access to flow cytometry instrumentation used in the experiments described. Lastly, we thank Joelle Eaves, Annie Needs, and Logan Garland for providing critical feedback during the writing process. B.D. and D.R.W. acknowledge funding from USDA-NIFA-AFRI-009003 (GRANT13700968). B.D. also acknowledges partial support by a fellowship from Michigan State University under the Training Program in Plant Biotechnology for Health and Sustainability (T32-GM152798).

## Supporting Information

Additional experimental details, materials, and methods, including relevant schemes, tables, and raw cytometry data.

## Notes

### Competing Interest Statement

The authors have declared no competing interest.

### Summary of Updates

In this revised version, we have repeated and expanded the flg22-TAMRA titration experiments to 100 uM, added no-stress controls, included the FLS2LRR-S390K negative-control variant, and evaluated the N388S variant by titration. These new data clarified that the current flow cytometry assay does not support reliable KD determination because the curves do not saturate and the S390K control retains concentration-dependent signal. We therefore revised the manuscript to avoid overclaiming a validated quantitative affinity-screening platform and instead frame the work as identifying conditions and variants that produce an enrichment-compatible flg22-responsive display phenotype. We also revised the manuscript to more clearly motivate partial glycosylation modulation, explain why the fully deglycosylated NtoS construct is not the preferred engineering scaffold, and distinguish between observed flg22-TAMRA-positive phenotypes and definitive receptor-level binding. In addition, we clarified the interpretation of the magnetic bead enrichment assay, added two-factor ANOVA analysis for Figure 5, improved figure labeling and axis consistency, combined the supplemental materials into a single document, and revised the abstract, transitions, figure legends, and supporting information for clarity.

