## Supporting Information for "Development of approaches and identifying limitations towards functional yeast surface display of FLS2"

**Additional experimental details, materials, and methods, including relevant schemes and tables**

**Title:**

**Table of Contents:**

**S1 |** Illustration of Aga2-HA-FLS2 surface displayed on *S. cerevisiae*

**S2 |** Linear yeast surface display plasmid map including the Aga2-HA-FLS2 ectodomain open reading frame

**S3 |** Surface displayed Aga2-HA-FLS2 ectodomain shows good expression with negligible flg22-TAMRA binding

**S4 |** FLS2 glycosylation sites *(*NXS/T*)* are highly-conserved among select flowering dicotyledons*.*

**S5 |** The size and saccharide composition of eukaryotic N-linked glycans varies between lineages

**S6 |** Quantified high-mannose N-glycan mass range aligns with de-glycosylated FLS2 size difference for 22 putative N-linked glycosylation sites

**S7 |** Alphafold2 predictions and Rosetta energy calculation for Non-glycosylated FLS2LRR design

**S8 |** Modeling high-mannose glycosylation on the FLS2 ectodomain may be informative for identifying an intermediate glycosylation state variant

**S9 |** Tunicamycin concentration effects the lipid-linked oligosaccharide (LLO) pool available for transfer by oligosaccharyltransferase (OST) complex subunits onto FLS2LRR bound for surface display

**S10 |** Magnetic bead sorting allows enrichment of >1 uM affinity binders

**S11 |** Magnetic bead enrichment of FLS2LRR cells by biotin-flg22 coated beads from a starting 1:1000 (binder:non-binder) ratio

**SI1 |** Conditional differences that influence interpretation of magnetic bead enrichment experiments

**ST1 |** Two-factor ANOVA for Figure 5 across treatments, variants, and combined treatment and variant groups

**ST2 |** Primer sequences for molecular cloning

**SI2 |** DNA sequences for yeast surface display inserts

**SI3 |** pCT80 template DNA expression vector

**SI4 |** Example gating and calculation of background subtracted fraction binding

**S12 |** Gating strategy for flow cytometry experiments

**SI5a-q |** Raw flow cytometry density plots

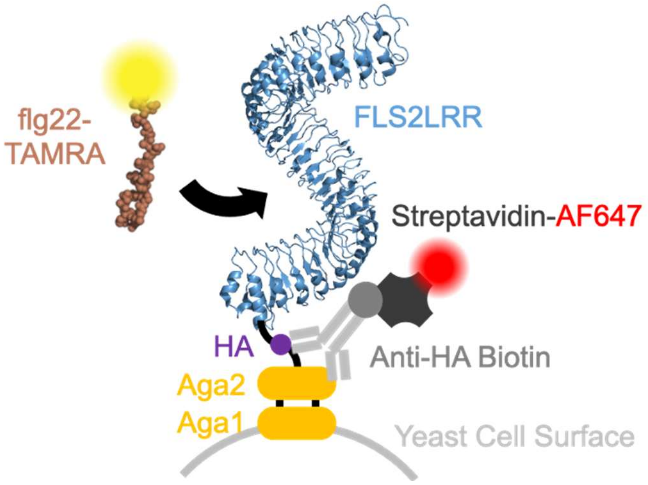

**Supplementary Figure 1: Illustration of Aga2-HA-FLS2 surface displayed on *S. cerevisiae*.** N-terminal Aga2 interacts with Aga1 using disulfide bridges to immobilize C-terminal surface displayed protein on the yeast cell surface, here showing the FLS2 ectodomain (PDB ID 4MN8, Chain A (1)). Protein expression is detected using anti-HA-biotin primary and streptavidin-AlexaFluor647 secondary antibodies. Binding of flg22 is detected using fluorescence from TAMRA N-terminally conjugated to flg22. flg22-TAMRA was chosen as the fluorophore-conjugated peptide for binding assays since it has been previously validated for molecular imaging (2).

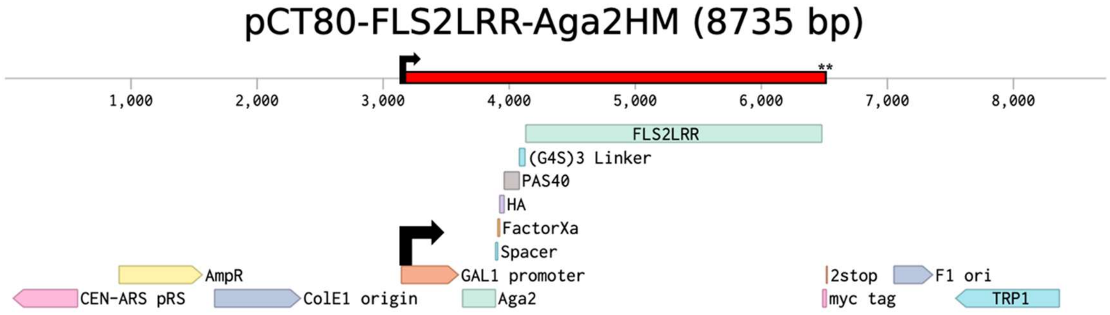

**Supplementary Figure 2: Linear yeast surface display plasmid map including the Aga2-HA-FLS2 ectodomain open reading frame.** Galactose inducible expression utilizes the GAL1 promoter. At the N-terminus, the A-agglutinin-binding subunit (AGA2) binds to the yeast cell wall A-agglutinin-anchorage subunit (AGA1) using two disulfide bonds. Downstream, AGA2 is followed by a spacer sequence buffer, Factor Xa cut site, Human Influenza Hemagglutinin (HA)-tag, and multiple linkers before the FLS2 ectodomain. At the C-terminus, for flow cytometry and western blot staining, myc-tag cannot be labelled due to a Kex2 (KR) cut site at the C-terminal end of the FLS2 ectodomain, which was mutated to KK to allow myc-tag labelling (data not shown) if necessary. TRP1 is the auxotrophic marker used for yeast surface display plasmid-containing cell selection after transformation. Red boxed region indicates the span of the Aga2-HA-FLS2 open reading frame. Two asterisks indicate the location of both stop codons.

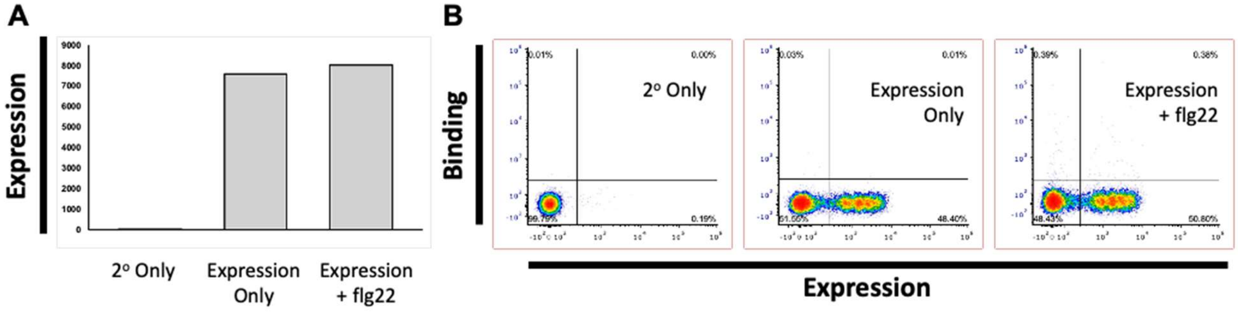

**Supplementary Figure 3: Surface displayed Aga2-HA-FLS2 ectodomain shows good expression with negligible flg22-TAMRA binding.** FLS2LRR was displayed on the surface of yeast and stained to detect expression (anti-HA-biotin (1º) + strep-AF647 (2º)) and flg22 binding (flg22-TAMRA purified peptide) for flow cytometry screening. **(A)** Y-axis shows right half quadrants expression positive event count for – 2º Only Staining, Expression staining, All detection reagent staining with 10 uM flg22-TAMRA. **(B)** Density plots for binding (flg22-TAMRA) and expression (HA-tag staining, AlexaFluor647) show negligible flg22-TAMRA binding despite high FLS2 ectodomain surface expression. At lower concentration flg22-TAMRA treatments (100 nM, 1 uM), negligible flg22-TAMRA binding is also observed (data not shown).

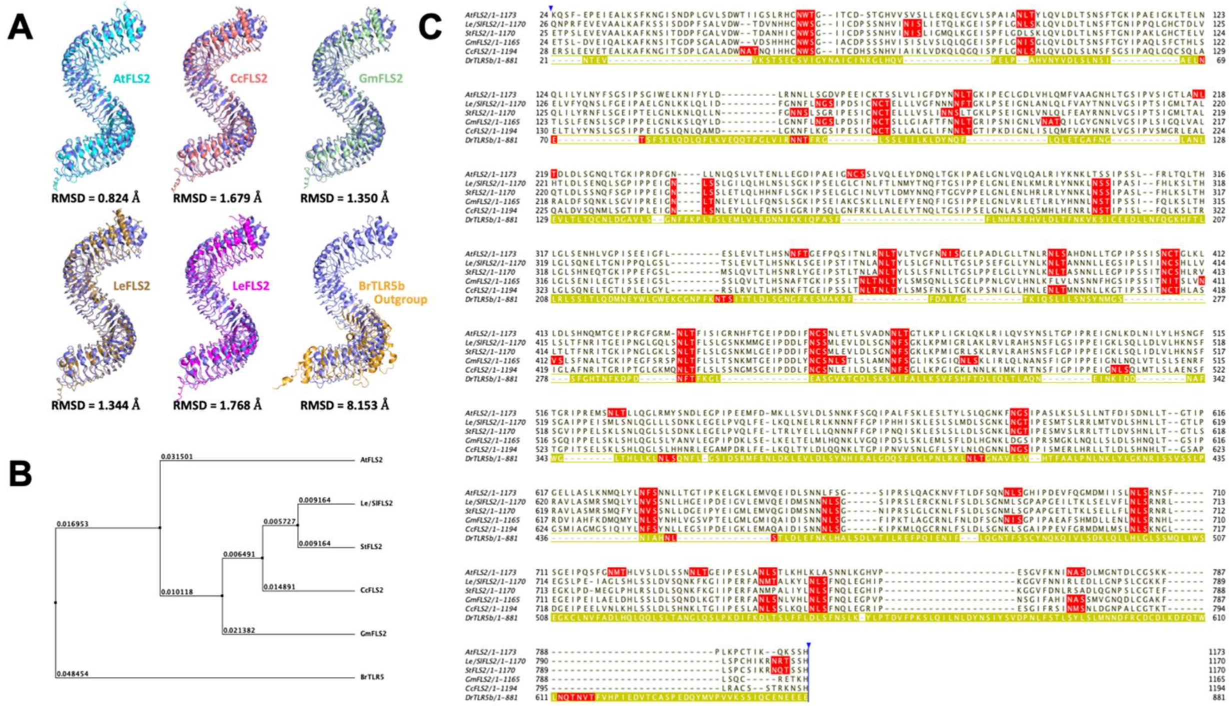

**Supplementary Figure 4. FLS2 glycosylation sites (NXS/T) are highly-conserved among select flowering dicotyledons.** Selected magnoliopsid (dicotyledons) class plant FLS2 ectodomains relevant to FLS2 research and sustainable agriculture were aligned in Jalview (3) using MAFFT (4) including the LRR-RLK outgroup Toll-Like Receptor 5b (TLR5b) from *Brachydanio rerio* (Zebrafish). **(A)** Using the Alphafold Server (5), five FLS2 and one TLR5b structures were modeled, then superimposed to the FLS2 crystal structure (PDB ID 4MN8) in PyMOL (6). **(B)** Using Jalview, phylogenetic tree was calculated using average distances and sequence feature similarity (using NXS/T as the feature). **(C)** The MAFFT alignment (default settings) of FLS2s and TLR5b (yellow highlight) including only FLS2 ectodomains shows high NXS/T site (red highlight) conservation between FLS2 orthologs. CcFLS2: Citrus x clementina (clementine); GmFLS2: Glycine max (soybean); LeFLS2: Solanum lycopersicum (garden tomato); StFLS2: Solanum tuberosum (irish potato); AtFLS2: Arabidopsis thaliana (mouse-ear cress).

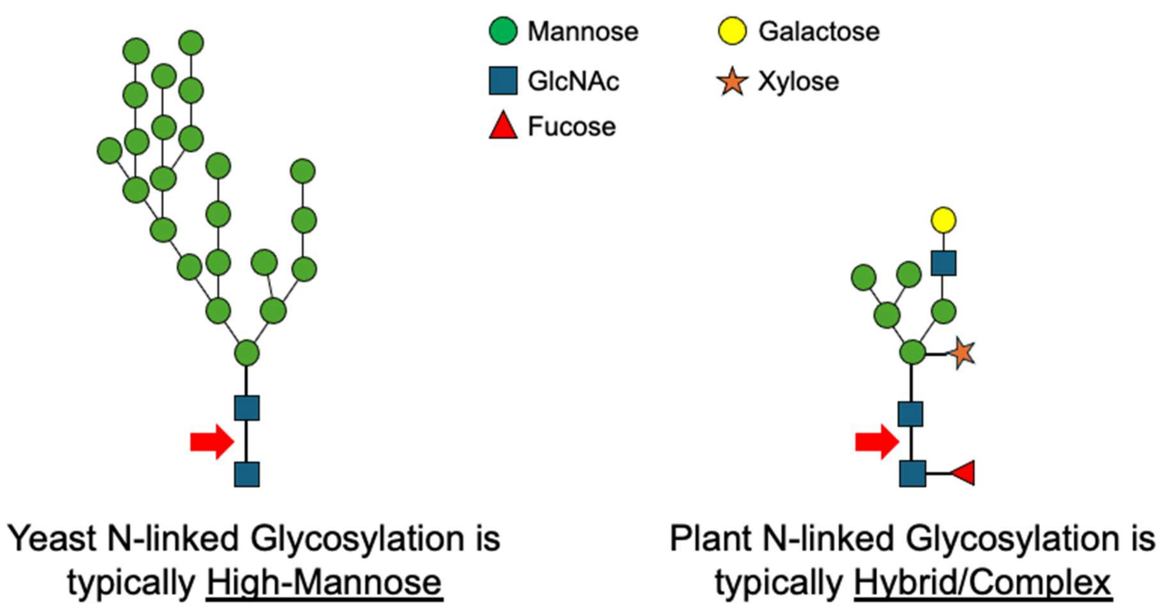

**Supplementary Figure 5: The size and saccharide composition of eukaryotic N-linked glycans varies between lineages.** *Saccharomyces cerevisiae* (baker's yeast) is used as the surface display expression system throughout all experiments and typical yeast glycoproteins have high-mannose N-linked glycans (7). In many plants, high-mannose glycosylation is rare and instead glycoproteins are decorated with hybrid/complex N-linked glycans. To release glycan structures from terminal GlcNAc-Asn of glycoproteins, the enzyme endoglycosidase H (Endo H) cuts after the terminal GlcNAc (red arrow).

A B

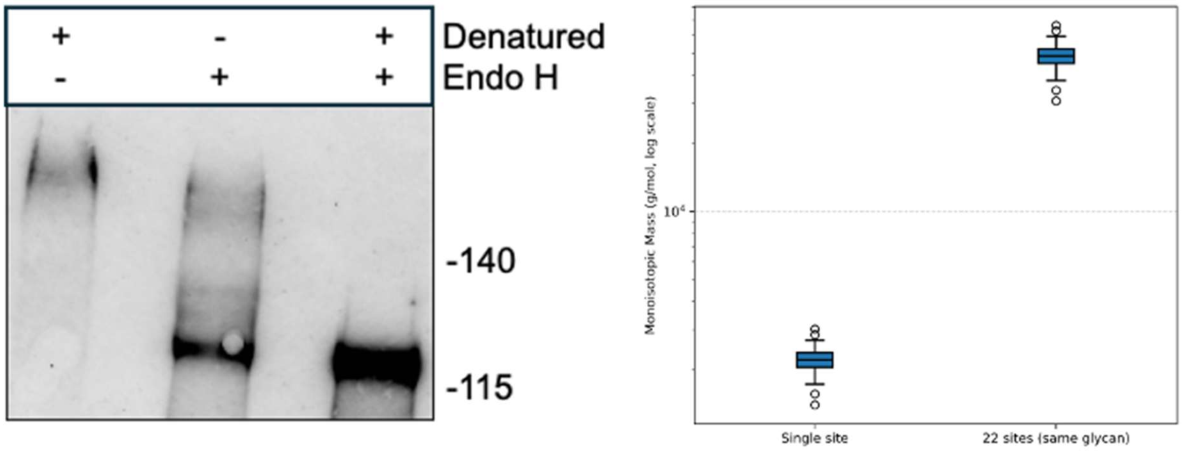

C

| Monoisotopic Mass (g/mol) | Minimum | Maximum | Mean |
|---|---|---|---|
| ***Single Entries*** | 1396.49 | 3014.89 | 2218.96 |
| ***22 Putative Sites*** | 30722.78 | 66327.58 | 48817.17 |

**Supplementary Figure 6: Quantified high-mannose N-glycan mass range aligns with de-glycosylated FLS2 size difference for 22 putative N-linked glycosylation sites. (A)** Western blot analysis of denatured FLS2 before and after endoglycosidase H treatment shows >30 kDa mass difference. Using the GlyCosmos database (https://glycosmos.org/), "Saccharomyces cerevisiae" was used as search input to identify entries in GlyCosmos Glycans. Entries were downloaded as CSV and datasheet was filtered by the "N-Glycan high mannose" label in the Motifs column using Microsoft Excel. A filtered list containing Monoisotopic Mass column entries was used to calculate an additional column where each entry is multiplied by 22, **(B)** then plotted to visualize single site and 22 site (same glycan) mass range. **(C)** Values in these column were used to calculate the minimum, maximum, and mean monoisotopic mass for single entries and 22 putative sites.

A

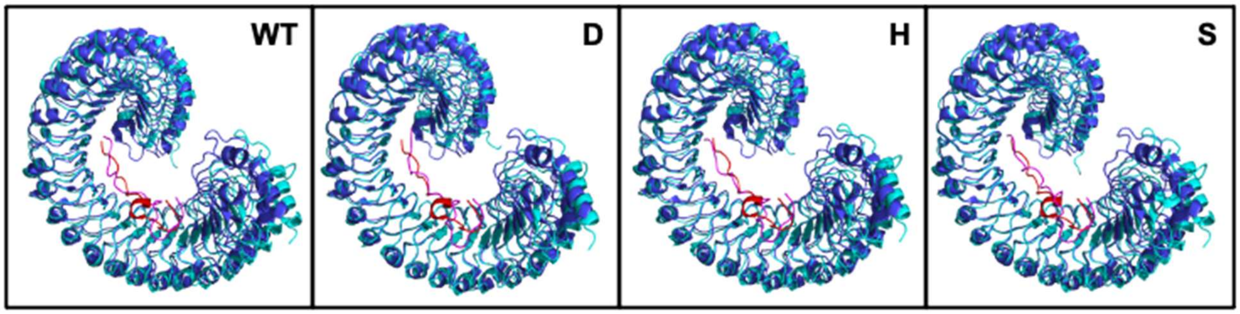

B

| Substitution at all NxS/T | RMSD (Å) | ipTM | Rosetta total_score (REU) |
|---|---|---|---|
| WT | 2.148 | 0.88 | -2671 |
| D (Aspartic Acid) | 1.447 | 0.87 | -2680 |
| H (Histidine) | 1.819 | 0.90 | -2699 |
| S (Serine) | 1.601 | 0.90 | -2672 |
| P (Proline) | 1.951 | 0.90 | -1342 |
| W (Tryptophan) | 2.192 | 0.90 | -1745 |

**Supplementary Figure 7: Alphafold2 predictions and Rosetta energy calculation for Non-glycosylated FLS2LRR design.** Amino acids were chosen to replace asparagine at NxS/T motifs based on the frequency of substitution (BLOSUM62). **(A)** The crystal structure of *A. thaliana* FLS2 ectodomain (PDB ID: 4MN8) was used to predict structures where NxS/T asparagine was modified to either aspartic acid (D), histidine (H), and serine (S). Color coding for structures is as follows: 4MN8 FLS2 crystal structure (Blue), AlphaFold predicted FLS2 (Teal), 4MN8 flg22 crystal structure (Red), AlphaFold predicted flg22 (Magenta). **(B)** Predicted structures were compared to 4MN8 crystal structure using RMSD from PyMOL structural alignment. AlphaFold ipTM score was obtained from Alphafold2 Multimer (8) predictive modeling. The total_score was calculated from Rosetta molecular modeling software (9) following a Fast Relax. Mutation to proline and tryptophan were used as negative controls.

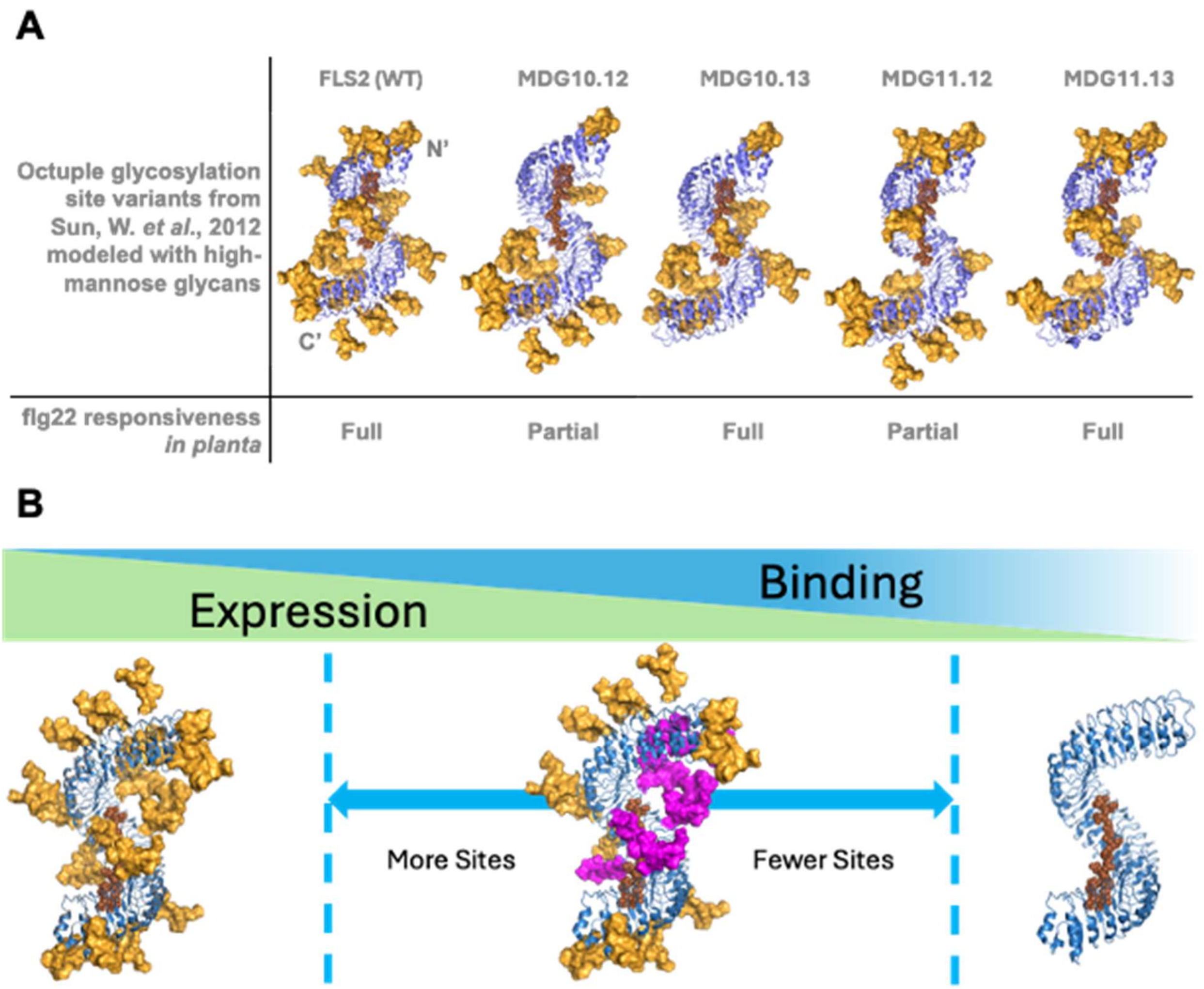

**Supplementary Figure 8. Modeling high-mannose glycosylation on the FLS2 ectodomain may be informative for identifying an intermediate glycosylation state variant.** The octuple glycosylation site variants from Sun,W. et al., 2012 (10) were **(A)** modeled with yeast high-mannose glycans (GlyTouCan ID G92042VQ) using the GlycoShape Re-Glyco tool (11). Notably, more glycosylation-scarce gaps around the flg22 binding interface and adjacent solvent-exposed, convex regions of the FLS2 ectodomain may explain partial flg22 responsiveness for variants MDG10.12 and MDG 11.12. Interestingly, predicted out-facing glycans near the C-terminus are absent from variants with full flg22 responsiveness. **(B)** Optimal FLS2 yeast display expression is dependent on the presence of glycosylation, however the specific glycosylated sites which interfere with flg22 binding in yeast may exist across a range of sites at the concave face or sterically occluding the flg22 binding interface (magenta).

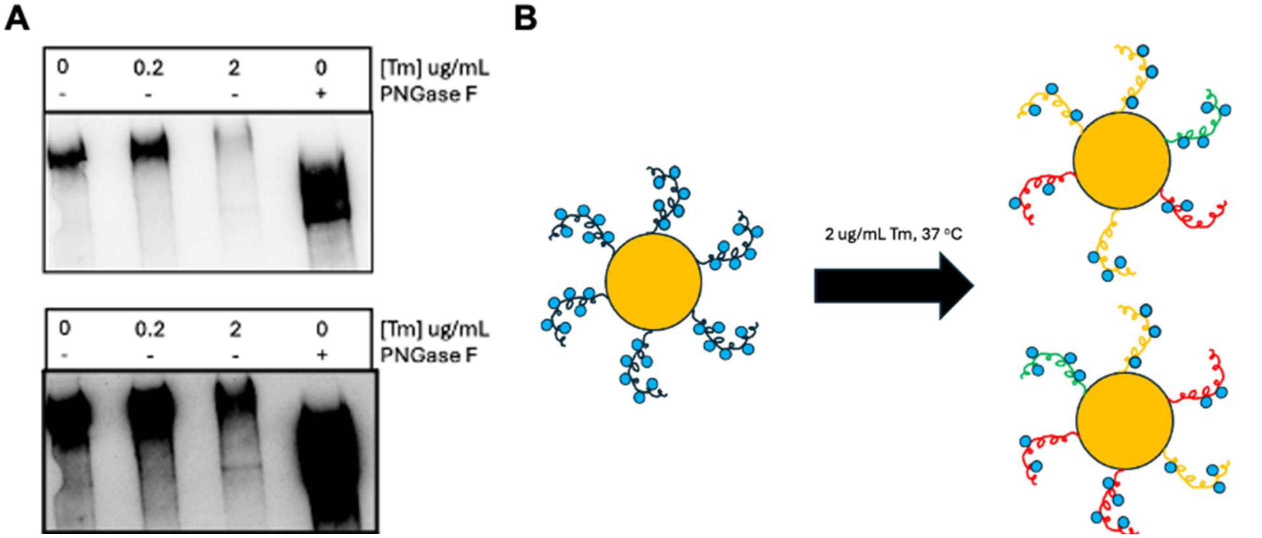

**Supplementary Figure 9: Tunicamycin concentrations effects the lipid-linked oligosaccharide (LLO) pool available for transfer by oligosaccharyltransferase (OST) complex subunits onto FLS2LRR bound for surface display. (A)** Western blot analysis of DTT-extracted FLS2LRR protein (HA-tag staining, AF647) after various concentrations of tunicamycin treatment during induction or endoglycosidase treatment. Due to decreased expression after tunicamycin treatment, modified contrast duplicate of top image is shown to emphasize presence of lower molecular weight FLS2LRR. **(B)** Hypothetical diagram of the FLS2LRR displaying cells and glycan heterogeneity of each FLS2LRR after 2 ug/mL tunicamycin, 37 $^{o}$C induction. Each individual cell is made up of 1-10e4 displaying protein that vary in the number of glycans present for each protein. Thus, if 10-20% of cytometry events are expression and flg22 binding positive after background subtraction (Green), we expect that this population is a fraction of displaying protein per cell across cells with FLS2LRR expression (Green, Orange, Red). Colors for categorizing the displaying protein with varying number of glycans and corresponding phenotype: **Green** – sufficient N-glycosylated sites/optimal binding/optimal expression; **Orange** – insufficient N-glycosylated sites/non-optimal binding/optimal expression; **Red** – insufficient N-glycosylated sites/non-optimal binding/non-optimal expression. In this hypothetical example, sufficient N-glycosylated sites represents a displaying protein with enough glycosylated sites to maintain proper folding and not interfere with flg22 binding in yeast.

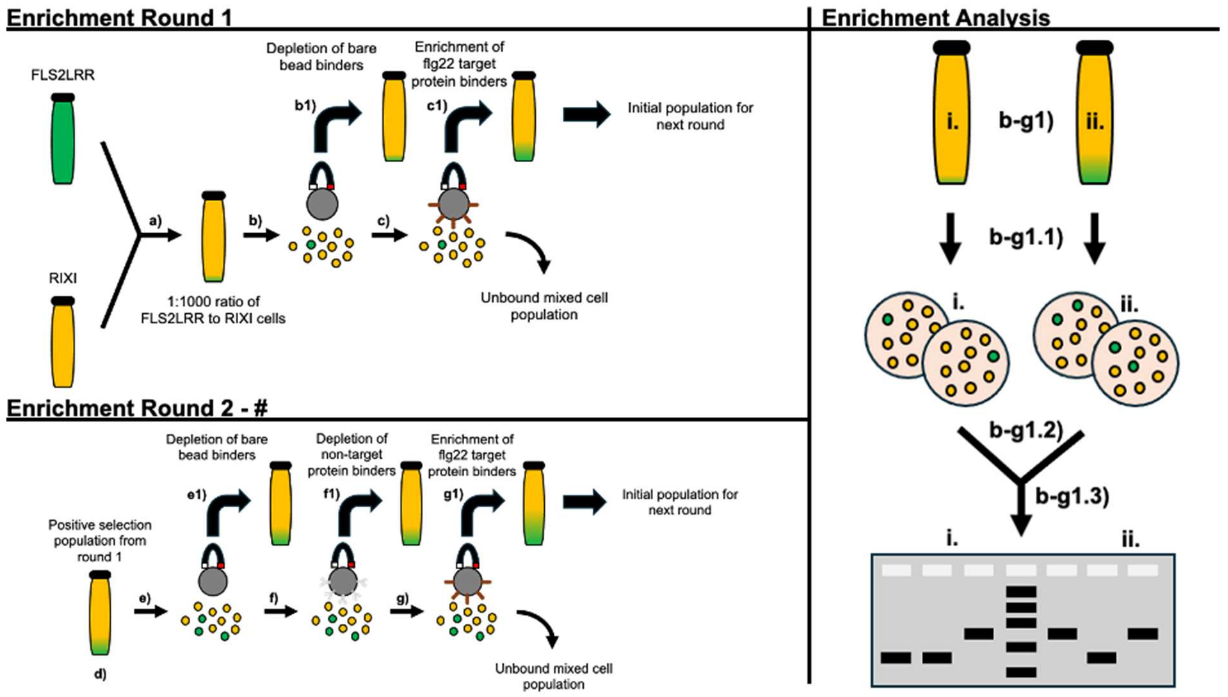

**Supplementary Figure 10. Magnetic bead sorting allows enrichment of >1 uM affinity binders.** FLS2LRR and RIXI cells are combined (a) after induction at additive or no stress conditions for two replicates per induction condition group. During enrichment round 1, bare streptavidin-bead negative selection (b) and biotin-flg22 coated streptavidin-bead positive selection (c) are performed on the cell mixtures. After each selection (b-c1), bead and cell mixtures are collected and processed for enrichment analysis (right). Only the positive selection mixture from each round is carried over to the following round (d), where growth and induction is repeated before the following round of enrichment. During enrichment round 2 and subsequent rounds, bare streptavidin-bead negative selection (e), biotin-IgG coated streptavidin-bead negative selection (f), and biotin-flg22 coated streptavidin-bead positive selection (g) are performed on the cell mixtures. After each selection (e-g1), bead and cell mixtures are collected and processed for enrichment analysis. To determine enrichment of FLS2-flg22 binding clones, respective cell mixtures (i-ii) are plated on tryptophan-deficient selection plates (b-g1.1). Then, 2-3 single colony clones per replicate are picked and independently grown for DNA extraction to obtain FLS2 ectodomain- or RIXI-containing surface display plasmids (b-g1.2). Finally, amplification around the FLS2LRR/RIXI insert region for each clone is performed and evaluated using DNA gel electrophoresis (b-g1.3). The expected nucleotide length for RIXI and FLS2LRR amplicons is ~1,100 bp and ~2,700 bp, respectively, allowing to distinguish between clones compared to a 1kb DNA ladder.

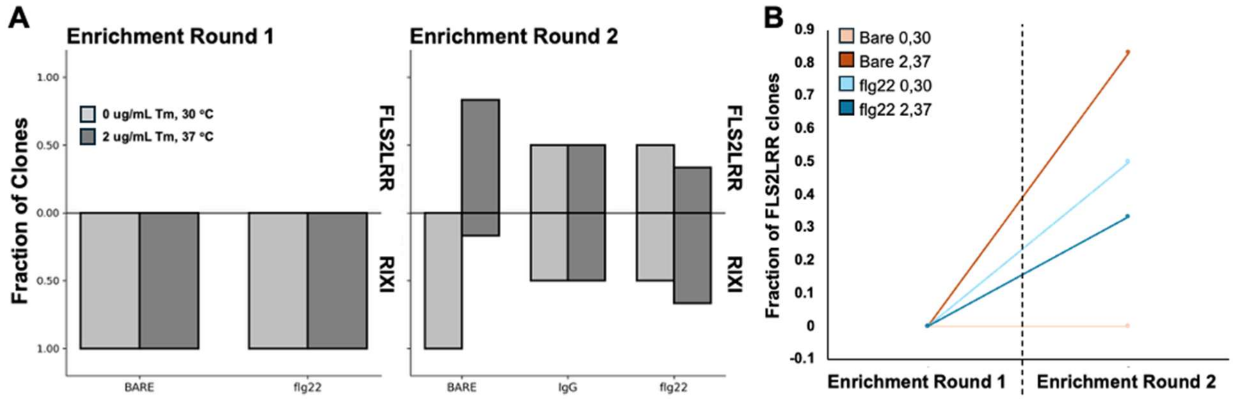

**Supplementary Figure 11: Magnetic bead enrichment of FLS2LRR cells by biotin-flg22 coated beads from a starting 1:1000 (binder:non-binder) ratio.** Following the protocol described in **Figure S10**, **(A)** FLS2LRR cells are enriched after outgrowth of round one positive selection cells. Cells plated on auxotrophic (tryptophan-deficient) selection plates are grown out for enrichment analysis after negative and positive bead selection. After round two selections there is at least 330-fold enrichment (33% of clones) of FLS2LRR cells except for bare bead negative selection of no stress induction clones. Using positive selection cells for round two increases the likelihood of FLS2LRR selection in all induction treatment and selection groups (**Supplementary Information 1**). However, **(B)** bare bead selection in round two only shows FLS2LRR enrichment in additive stress induction group, supporting FLS2LRR gain of specificity for flg22 aided by additive stress conditions. Bead saturation with cells in both rounds increases the likelihood of non-binder enrichment in round two (**Supplementary Information 1**). However, despite 1:1000 starting ratio of binder:non-binder in round one, FLS2LRR is enriched in almost all groups in round two.

**Supplementary Information 1. Conditional differences that influence interpretation of magnetic bead enrichment experiments from figures S10-11.** In a standard magnetic bead sort with yeast surface display (e.g., Ackerman et al., Biotechnol Prog 2009 (12)), researchers may start from a mixture with ~1:1000 binders:non-binders and use sequential depletion on bare beads and non-target–coated beads, followed by enrichment on target-coated beads. Because all "binder" cells typically display the same protein with a uniform glycosylation pattern, this scheme enriches target-specific binders on target beads while depleting non-specific binders using bare and non-target beads, and very few target-specific binders are captured during the depletion steps. In our modified FLS2 display system, after induction under tunicamycin and thermal stress, that assumption breaks down: each FLS2-expressing yeast cell displays a heterogeneous mixture of FLS2 glycoforms because tunicamycin partially inhibits N-linked glycosylation at the 22 putative NxS/T sites in a stochastic fashion. As a result, even within a single "binder" cell, some FLS2 molecules may carry different numbers and patterns of glycans (e.g. 12 vs. 15 modified sites, or different combinations of sites), yielding a spectrum of surface variants that can differ in both flg22 binding and non-specific stickiness.

Practically, after overnight additive and no stress induction, we began with a 1:1000 mixture of FLS2LRR (binder) and RIXI (non-binder) yeast on Day 1, depleted cells that bound bare beads, and then enriched those captured by flg22-coated beads. On Day 2, we re-induced the flg22-enriched population under additive stress and no stress, then performed sequential selection: depletion on bare beads, depletion on non-target–coated beads, and enrichment on flg22-coated beads, and quantified FLS2LRR vs. RIXI in each bead-bound fraction. After the second round, we observed strong enrichment of FLS2LRR (≥330-fold, to ~33% of clones) in almost all induction and selection conditions, including bare-bead and flg22-bead sorts, except for the bare-bead negative selection of the no-stress induction group. Using the flg22-positive fraction from round one as input to round two further increased the likelihood of recovering FLS2LRR across conditions. Round-two bare-bead selections showed FLS2LRR enrichment only under additive stress conditions, consistent with stress-dependent changes in FLS2LRR surface properties that both increase nonspecific stickiness and maintain flg22 binding. Bead saturation by cells in both rounds likely increased the chance of non-binder carryover, and our small sampling numbers (n=2 or n=3) add uncertainty to the measurements. Nonetheless, the robust FLS2LRR enrichment in round two despite a 1:1000 starting ratio, together with the increased "stickiness" of stressed cells seen by flow cytometry, is consistent with a model in which tunicamycin-induced heterogeneous glycosylation and stress-induced surface changes allow FLS2-expressing cells to be captured on multiple bead types, rather than cleanly separating binders and non-binders as in a classical magnetic bead sorting campaign.

**Supplementary Table 1. Two-Factor ANOVA with replication for Figure 5 across treatments, variants, and combined treatment and variants groups.** Background subtracted fraction binding for all sample group replicates was used for calculating significant differences using two-factor ANOVA with replication. Calculations were performed using Microsoft Excel.

| Anova: Two-Factor With Replication | | | | | | | |
|---|---|---|---|---|---|---|---|
| SUMMARY | FL | Var 1 | Var 3 | Var 5 | Var 7 | NtoS | Total |
| *0, 37* | | | | | | | |
| Count | 3 | 3 | 3 | 3 | 3 | 3 | 18 |
| Sum | 0.13178205 | 0.144571478 | 0.37145437 | 0.260024612 | 0.10720311 | 0.34612059 | 1.36115621 |
| Average | 0.04392735 | 0.048190493 | 0.12381812 | 0.086674871 | 0.03573437 | 0.11537353 | 0.07561979 |
| Variance | 0.00016674 | 0.000226177 | 0.00052632 | 0.004885911 | 0.00035601 | 0.00054528 | 0.00209016 |
| *2, 37* | | | | | | | |
| Count | 3 | 3 | 3 | 3 | 3 | 3 | 18 |
| Sum | 0.59391665 | 1.271191592 | 0.93307045 | 0.929332307 | 0.70648135 | 0.8447722 | 5.27876455 |
| Average | 0.19797222 | 0.423730531 | 0.31102348 | 0.309777436 | 0.23549378 | 0.28159073 | 0.2932647 |
| Variance | 0.00113275 | 0.000287835 | 0.00015235 | 0.003445763 | 0.00661737 | 0.00033332 | 0.00673119 |
| *Total* | | | | | | | |
| Count | 6 | 6 | 6 | 6 | 6 | 6 | |
| Sum | 0.72569869 | 1.41576307 | 1.30452483 | 1.18935692 | 0.81368445 | 1.19089279 | |
| Average | 0.12094978 | 0.235960512 | 0.2174208 | 0.198226153 | 0.13561408 | 0.19848213 | |
| Variance | 0.00763874 | 0.042514701 | 0.01078522 | 0.018265096 | 0.0147605 | 0.00863989 | |
| ANOVA | | | | | | | |
| *urce of Variati* | *SS* | *df* | *MS* | *F* | *P-value* | *F crit* | |
| Sample | 0.42632375 | 1 | 0.42632375 | 273.9308234 | 1.251E-14 | 4.25967727 | Treatment |
| Columns | 0.06326589 | 5 | 0.01265318 | 8.130195265 | 0.00013245 | 2.62065415 | Variant |
| Interaction | 0.04934533 | 5 | 0.00986907 | 6.34128772 | 0.00069393 | 2.62065415 | Variant x Treatment |
| Within | 0.03735166 | 24 | 0.00155632 | | | | |
| Total | 0.57628663 | 35 | | | | | |

**Supplementary Table 2.** Primer sequences for molecular cloning.

| Purpose | Sequence (5' -> 3') |
|---|---|
| Homologous end overlapping primers for amplification of FLS2LRR for Gibson cloning into pCT80 | F: TGGTGGGGGCGGATCTGCTAGCAAACAATCATTTGAGCCAGAGATCG<br>R: GAAATAAGCTTTTGTTCGGATCCACGAGTCCTCTTACTGAAGTGG |

**Supplementary Information 2. DNA sequences for yeast surface display inserts**. For the select site FLS2LRR variants, the position with an asparagine to serine substitution is highlighted in red.

*S. cerevisiae codon optimized FLS2LRR insert sequence:*

AAACAATCATTTGAGCCAGAGATCGAAGCACTTAAGTCCTTCAAAAATGGCATAAGCAACGACCCCCTG
GGAGTCTTAAGCGATTGGACCATTATTGGGAGTCTTCGTCATTGCAACTGGACCGGTATTACGTGCGAC
AGCACTGGTCACGTCGTGTCTGTTTCACTTTTGGAGAAGCAGTTGGAGGGAGTCTTGAGTCCAGCGATC
GCTAACCTTACTTATTTACAAGTCTTGGACCTTACAAGTAATAGTTTTACAGGGAAGATACCGGCGGAAA
TTGGTAAACTTACGGAGCTTAATCAGTTGATTCTGTATCTTAACTACTTCAGCGGATCTATTCCATCAGGA
ATCTGGGAACTGAAGAATATATTTTATTTAGACTTGCGTAATAACCTACTATCCGGTGATGTCCCAGAGG
AGATATGTAAGACCTCCAGCTTAGTTCTTATTGGGTTTGACTACAATAATCTTACAGGCAAGATCCCGGA
GTGTTTAGGGGACCTTGTGCATCTGCAGATGTTTGTCGCCGCAGGGAATCATTTGACCGGAAGTATACC
TGTGAGTATTGGGACATTGGCAAATCTGACCGATCTAGATCTTTCAGGCAATCAGTTGACTGGAAAAAT
ACCGAGAGATTTCGGTAACTTACTTAATCTTCAGTCTCTTGTACTGACCGAGAATCTGCTAGAGGGAGAC
ATACCGGCTGAGATAGGAAACTGCTCTTCCTTAGTCCAGTTAGAATTATATGACAACCAACTGACAGGCA
AAATCCCTGCTGAACTAGGAAACCTAGTACAACTACAAGCGTTGCGTATTTACAAAAACAAACTAACATC
TTCAATACCATCAAGCCTGTTTCGTCTGACTCAGCTAACTCATCTAGGTCTTAGCGAGAACCACCTGGTCG
GACCTATCAGTGAAGAGATCGGTTTCCTTGAAAGCCTTGAAGTTCTTACCTTACATTCCAACAATTTCACC
GGGGAGTTCCCACAGAGTATCACAAATTTACGTAACTTGACTGTGTTGACTGTTGGCTTCAACAATATCA
GTGGCGAATTGCCTGCTGATCTAGGGCTGTTAACGAATCTGCGTAATTTAAGCGCGCATGATAATTTGCT
TACAGGCCCTATACCGTCCAGTATCTCAAATTGTACAGGCCTTAAATTATTGGACCTGTCTCACAATCAAA
TGACCGGCGAAATCCCACGTGGATTTGGCAGGATGAACCTTACGTTCATATCAATAGGCCGTAACCATTT
CACCGGCGAAATACCCGATGATATCTTTAATTGTAGTAATTTGGAGACCTTATCTGTGGCGGATAACAAC
CTTACTGGTACCCTTAAACCCCTAATCGGAAAGCTGCAAAAGCTTAGAATCCTACAGGTCTCCTACAATT
CATTAACCGGGCCTATCCCCAGGGAGATAGGGAACTTGAAGGACTTAAACATCCTTTATCTACACTCAAA
TGGTTTCACAGGTAGGATACCCAGAGAGATGAGCAATTTGACATTGCTACAAGGGTTGAGAATGTACTC

CAATGACTTGGAAGGCCCGATTCCCGAGGAAATGTTCGACATGAAGCTGTTGTCTGTGCTGGATCTTAG
CAATAATAAGTTTTCAGGCCAAATCCCAGCATTGTTTTCTAAATTAGAGTCACTTACATATCTGTCTCTTCA
AGGTAACAAGTTTAATGGCAGTATACCTGCAAGCTTGAAATCCTTGTCTTTATTGAACACATTCGATATA
AGTGACAATCTATTAACGGGAACGATTCCCGGTGAACTTCTTGCAAGCCTAAAGAACATGCAACTATATC
TGAACTTCTCTAACAACTTATTAACGGGGACTATTCCTAAGGAACTAGGCAAATTAGAGATGGTGCAAG
AAATAGACTTATCAAACAACTTGTTTTCAGGATCAATACCGAGGTCCTTGCAGGCATGTAAAAACGTCTT
TACGCTAGATTTTAGCCAGAACAACTTGTCCGGTCACATTCCTGACGAAGTTTTCCAGGGTATGGACATG
ATAATTTCCTTAAACTTGTCCAGAAACTCATTCTCCGGGGAGATCCCTCAGTCCTTTGGTAATATGACGCA
TCTAGTGAGCTTGGATCTTTCATCTAATAACCTAACTGGAGAGATCCCCGAGTCATTAGCAAATTTGAGC
ACGTTAAAACATCTAAAGCTTGCAAGCAATAATCTTAAAGGACATGTACCGGAGTCCGGAGTTTTTAAG
AATATTAACGCCAGTGACTTAATGGGTAACACGGATCTTTGTGGCAGTAAGAAGCCTCTAAAGCCATGC
ACCATAAAACAGAAATCCAGCCACTTCAGTAAGAGGACTCGT

*S. cerevisiae codon optimized NtoS insert sequence:*

AAACAATCTTTCGAACCAGAGATTGAAGCTTTGAAGTCTTTCAAGAACGGTATTTCTAACGACCCATTGG
GTGTTTTGTCCGACTGGACTATCATTGGTTCTTTAAGACACTGTTCTTGGACCGGTATCACTTGCGATTCT
ACTGGTCACGTTGTCTCCGTTTCCCTGTTGGAAAAGCAACTGGAAGGTGTTTTGTCTCCAGCCATTGCTTC
CCTTACCTACTTGCAAGTCTTGGACCTCACCTCCAATTCCTTCACTGGAAAAATTCCAGCCGAAATTGGTA
AATTGACCGAATTGAACCAATTGATTTTGTATTTGAACTACTTCTCTGGTTCTATTCCATCTGGTATCTGG
GAACTCAAGAATATTTTCTACTTGGACTTGAGAAATAACTTGTTGTCTGGTGACGTCCCAGAAGAAATCT
GTAAGACCTCTTCTTTAGTATTGATTGGTTTCGACTACAACTCATTGACTGGTAAGATACCAGAATGTCTC
GGTGATTTAGTCCACTTACAAATGTTCGTTGCTGCTGGTAACCATTTGACGGGTTCCATCCCAGTTTCCAT
CGGTACTCTGGCTTCTTTGACTGATTTGGATTTGTCTGGGAACCAATTAACTGGTAAGATTCCAAGAGAT
TTTGGTAACCTGCTAAACTTGCAATCTTTGGTCTTGACTGAAAACTTATTGGAAGGTGAAATCCCAGCTG
AAATCGGTTCTTGTTCCTCTTTGGTTCAATTGGAATTGTACGATAACCAATTGACCGGTAAGATCCCTGCT
GAACTCGGTAATTTGGTTCAATTGCAAGCTTTGCGTATTTACAAGAACAAGTTGACTTCTTCTATCCCAAG
TTCGTTGTTCAGATTGACTCAATTGACTCATTTGGGTTTGTCTGAAAACCACCTTGTTGGTCCAATTTCCG
AAGAAATCGGTTTCTTGGAATCTTTAGAAGTCTTGACTTTGCACTCTAACTCATTCACCGGTGAATTCCCA
CAATCTATCACCAACTTGAGATCCTTGACTGTTTTGACCGTCGGTTTCAACTCTATCTCCGGTGAATTGCC
AGCTGATTTGGGTTTGTTGACCAACTTGAGATCTTTGTCCGCTCACGACAATCTTTTGACAGGTCCAATTC
CATCCTCTATCTCTTCTTGCACTGGTCTAAAATTGCTGGACTTATCTCACAACCAAATGACCGGTGAAATT
CCGAGAGGTTTCGGTCGTATGTCTCTCACTTTCATCTCCATCGGTAGAAACCACTTCACCGGTGAAATCCC
AGACGATATCTTCTCCTGTTCCAACTTGGAAACCCTTTCTGTCGCTGACAACTCTTTGACTGGTACCCTAA
AGCCATTGATCGGTAAGTTGCAAAAGTTGAGAATTTTGCAAGTTTCTTACAACTCGTTGACCGGTCCAAT
TCCAAGAGAAATTGGTAACCTAAAGGATTTAAACATCTTGTATTTGCACTCGAACGGTTTTACCGGTCGT
ATTCCAAGAGAAATGTCCTCCTTGACTTTGTTGCAAGGTCTAAGAATGTACTCTAACGATTTGGAAGGTC

CAATCCCTGAAGAAATGTTTGACATGAAGTTGTTATCTGTTTTGGATTTGTCCAATAACAAGTTCTCTGGT
CAAATCCCAGCTTTGTTTTCTAAGTTGGAATCCTTGACTTACTTGTCTTTGCAAGGTAACAAATTTTCCGG
CTCCATCCCAGCCTCTTTGAAATCTTTATCTTTGTTGAACACCTTCGATATTTCTGACAACCTATTAACAGG
TACCATCCCAGGTGAGCTATTGGCTAGTTTGAAGAACATGCAATTATACCTGTCTTTCTCTAACAACTTAC
TGACCGGTACCATTCCAAAGGAATTGGGTAAGCTAGAAATGGTCCAAGAAATCGACCTTTCCAACAACC
TGTTTTCCGGTTCCATTCCAAGATCACTACAAGCTTGTAAGAACGTTTTCACTTTGGACTTCTCTCAAAAC
AGTTTATCTGGTCACATTCCAGATGAAGTTTTCCAAGGTATGGACATGATTATCTCTTTGTCCTTGAGCAG
AAACTCTTTCTCTGGTGAAATTCCTCAATCCTTCGGTTCCATGACCCACTTGGTCAGCTTAGACCTTTCCTC
CAACAGCTTGACCGGTGAAATTCCAGAATCTTTGGCCTCCTTATCCACCTTAAAGCACTTAAAATTGGCC
AGTAACAATTTAAAGGGTCATGTTCCCGAAAGCGGTGTCTTCAAGAACATTTCTGCTTCTGACTTGATGG
GTAACACTGATTTGTGTGGTTCCAAGAAGCCCTTGAAGCCATGTACTATCAAGCAAAAGTCCTCTCATGC
ACCATAAAACAGAAATCCAGCCACTTCAGTAAGAGGACTCGT

*S. cerevisiae codon optimized FLS2LRR-Var1 insert sequence:*

AAACAATCATTTGAGCCAGAGATCGAAGCACTTAAGTCCTTCAAAAATGGCATAAGCAACGACCCCCTG
GGAGTCTTAAGCGATTGGACCATTATTGGGAGTCTTCGTCATTGCAACTGGACCGGTATTACGTGCGAC
AGCACTGGTCACGTCGTGTCTGTTTCACTTTTGGAGAAGCAGTTGGAGGGAGTCTTGAGTCCAGCGATC
GCTAACCTTACTTATTTACAAGTCTTGGACCTTACAAGTAATAGTTTTACAGGGAAGATACCGGCGGAAA
TTGGTAAACTTACGGAGCTTAATCAGTTGATTCTGTATCTTAACTACTTCAGCGGATCTATTCCATCAGGA
ATCTGGGAACTGAAGAATATATTTTATTTAGACTTGCGTAATAACCTACTATCCGGTGATGTCCCAGAGG
AGATATGTAAGACCTCCAGCTTAGTTCTTATTGGGTTTGACTACAATAATCTTACAGGCAAGATCCCGGA
GTGTTTAGGGGACCTTGTGCATCTGCAGATGTTTGTCGCCGCAGGGAATCATTTGACCGGAAGTATACC
TGTGAGTATTGGGACATTGGCAAATCTGACCGATCTAGATCTTTCAGGCAATCAGTTGACTGGAAAAAT
ACCGAGAGATTTCGGTAACTTACTTAATCTTCAGTCTCTTGTACTGACCGAGAATCTGCTAGAGGGAGAC
ATACCGGCTGAGATAGGAAACTGCTCTTCCTTAGTCCAGTTAGAATTATATGACAACCAACTGACAGGCA
AAATCCCTGCTGAACTAGGAAACCTAGTACAACTACAAGCGTTGCGTATTTACAAAAACAAACTAACATC
TTCAATACCATCAAGCCTGTTTCGTCTGACTCAGCTAACTCATCTAGGTCTTAGCGAGAACCACCTGGTCG
GACCTATCAGTGAAGAGATCGGTTTCCTTGAAAGCCTTGAAGTTCTTACCTTACATTCCAACAATTTCACC
GGGGAGTTCCCACAGAGTATCACAAATTTACGTAACTTGACTGTGTTGACTGTTGGCTTCAACAATATCA
GTGGCGAATTGCCTGCTGATCTAGGGCTGTTAACGAATCTGCGTTCTTTAAGCGCGCATGATAATTTGCT
TACAGGCCCTATACCGTCCAGTATCTCAAATTGTACAGGCCTTAAATTATTGGACCTGTCTCACAATCAAA
TGACCGGCGAAATCCCACGTGGATTTGGCAGGATGAACCTTACGTTCATATCAATAGGCCGTAACCATTT
CACCGGCGAAATACCCGATGATATCTTTAATTGTAGTAATTTGGAGACCTTATCTGTGGCGGATAACAAC
CTTACTGGTACCCTTAAACCCCTAATCGGAAAGCTGCAAAAGCTTAGAATCCTACAGGTCTCCTACAATT
CATTAACCGGGCCTATCCCCAGGGAGATAGGGAACTTGAAGGACTTAAACATCCTTTATCTACACTCAAA
TGGTTTCACAGGTAGGATACCCAGAGAGATGAGCAATTTGACATTGCTACAAGGGTTGAGAATGTACTC

CAATGACTTGGAAGGCCCGATTCCCGAGGAAATGTTCGACATGAAGCTGTTGTCTGTGCTGGATCTTAG
CAATAATAAGTTTTCAGGCCAAATCCCAGCATTGTTTTCTAAATTAGAGTCACTTACATATCTGTCTCTTCA
AGGTAACAAGTTTAATGGCAGTATACCTGCAAGCTTGAAATCCTTGTCTTTATTGAACACATTCGATATA
AGTGACAATCTATTAACGGGAACGATTCCCGGTGAACTTCTTGCAAGCCTAAAGAACATGCAACTATATC
TGAACTTCTCTAACAACTTATTAACGGGGACTATTCCTAAGGAACTAGGCAAATTAGAGATGGTGCAAG
AAATAGACTTATCAAACAACTTGTTTTCAGGATCAATACCGAGGTCCTTGCAGGCATGTAAAAACGTCTT
TACGCTAGATTTTAGCCAGAACAACTTGTCCGGTCACATTCCTGACGAAGTTTTCCAGGGTATGGACATG
ATAATTTCCTTAAACTTGTCCAGAAACTCATTCTCCGGGGAGATCCCTCAGTCCTTTGGTAATATGACGCA
TCTAGTGAGCTTGGATCTTTCATCTAATAACCTAACTGGAGAGATCCCCGAGTCATTAGCAAATTTGAGC
ACGTTAAAACATCTAAAGCTTGCAAGCAATAATCTTAAAGGACATGTACCGGAGTCCGGAGTTTTTAAG
AATATTAACGCCAGTGACTTAATGGGTAACACGGATCTTTGTGGCAGTAAGAAGCCTCTAAAGCCATGC
ACCATAAAACAGAAATCCAGCCACTTCAGTAAGAGGACTCGT

*S. cerevisiae codon optimized FLS2LRR-Var2 insert sequence:*

AAACAATCATTTGAGCCAGAGATCGAAGCACTTAAGTCCTTCAAAAATGGCATAAGCAACGACCCCCTG
GGAGTCTTAAGCGATTGGACCATTATTGGGAGTCTTCGTCATTGCAACTGGACCGGTATTACGTGCGAC
AGCACTGGTCACGTCGTGTCTGTTTCACTTTTGGAGAAGCAGTTGGAGGGAGTCTTGAGTCCAGCGATC
GCTAACCTTACTTATTTACAAGTCTTGGACCTTACAAGTAATAGTTTTACAGGGAAGATACCGGCGGAAA
TTGGTAAACTTACGGAGCTTAATCAGTTGATTCTGTATCTTAACTACTTCAGCGGATCTATTCCATCAGGA
ATCTGGGAACTGAAGAATATATTTTATTTAGACTTGCGTAATAACCTACTATCCGGTGATGTCCCAGAGG
AGATATGTAAGACCTCCAGCTTAGTTCTTATTGGGTTTGACTACAATAATCTTACAGGCAAGATCCCGGA
GTGTTTAGGGGACCTTGTGCATCTGCAGATGTTTGTCGCCGCAGGGAATCATTTGACCGGAAGTATACC
TGTGAGTATTGGGACATTGGCAAATCTGACCGATCTAGATCTTTCAGGCAATCAGTTGACTGGAAAAAT
ACCGAGAGATTTCGGTAACTTACTTAATCTTCAGTCTCTTGTACTGACCGAGAATCTGCTAGAGGGAGAC
ATACCGGCTGAGATAGGAAACTGCTCTTCCTTAGTCCAGTTAGAATTATATGACAACCAACTGACAGGCA
AAATCCCTGCTGAACTAGGAAACCTAGTACAACTACAAGCGTTGCGTATTTACAAAAACAAACTAACATC
TTCAATACCATCAAGCCTGTTTCGTCTGACTCAGCTAACTCATCTAGGTCTTAGCGAGAACCACCTGGTCG
GACCTATCAGTGAAGAGATCGGTTTCCTTGAAAGCCTTGAAGTTCTTACCTTACATTCCAACTCTTTCACC
GGGGAGTTCCCACAGAGTATCACAAATTTACGTAACTTGACTGTGTTGACTGTTGGCTTCAACTCTATCA
GTGGCGAATTGCCTGCTGATCTAGGGCTGTTAACGAATCTGCGTAATTTAAGCGCGCATGATAATTTGCT
TACAGGCCCTATACCGTCCAGTATCTCAAATTGTACAGGCCTTAAATTATTGGACCTGTCTCACAATCAAA
TGACCGGCGAAATCCCACGTGGATTTGGCAGGATGAACCTTACGTTCATATCAATAGGCCGTAACCATTT
CACCGGCGAAATACCCGATGATATCTTTAATTGTAGTAATTTGGAGACCTTATCTGTGGCGGATAACAAC
CTTACTGGTACCCTTAAACCCCTAATCGGAAAGCTGCAAAAGCTTAGAATCCTACAGGTCTCCTACAATT
CATTAACCGGGCCTATCCCCAGGGAGATAGGGAACTTGAAGGACTTAAACATCCTTTATCTACACTCAAA
TGGTTTCACAGGTAGGATACCCAGAGAGATGAGCAATTTGACATTGCTACAAGGGTTGAGAATGTACTC

CAATGACTTGGAAGGCCCGATTCCCGAGGAAATGTTCGACATGAAGCTGTTGTCTGTGCTGGATCTTAG
CAATAATAAGTTTTCAGGCCAAATCCCAGCATTGTTTTCTAAATTAGAGTCACTTACATATCTGTCTCTTCA
AGGTAACAAGTTTAATGGCAGTATACCTGCAAGCTTGAAATCCTTGTCTTTATTGAACACATTCGATATA
AGTGACAATCTATTAACGGGAACGATTCCCGGTGAACTTCTTGCAAGCCTAAAGAACATGCAACTATATC
TGAACTTCTCTAACAACTTATTAACGGGGACTATTCCTAAGGAACTAGGCAAATTAGAGATGGTGCAAG
AAATAGACTTATCAAACAACTTGTTTTCAGGATCAATACCGAGGTCCTTGCAGGCATGTAAAAACGTCTT
TACGCTAGATTTTAGCCAGAACAACTTGTCCGGTCACATTCCTGACGAAGTTTTCCAGGGTATGGACATG
ATAATTTCCTTAAACTTGTCCAGAAACTCATTCTCCGGGGAGATCCCTCAGTCCTTTGGTAATATGACGCA
TCTAGTGAGCTTGGATCTTTCATCTAATAACCTAACTGGAGAGATCCCCGAGTCATTAGCAAATTTGAGC
ACGTTAAAACATCTAAAGCTTGCAAGCAATAATCTTAAAGGACATGTACCGGAGTCCGGAGTTTTTAAG
AATATTAACGCCAGTGACTTAATGGGTAACACGGATCTTTGTGGCAGTAAGAAGCCTCTAAAGCCATGC
ACCATAAAACAGAAATCCAGCCACTTCAGTAAGAGGACTCGT

*S. cerevisiae codon optimized FLS2LRR-Var3 insert sequence:*

AAACAATCATTTGAGCCAGAGATCGAAGCACTTAAGTCCTTCAAAAATGGCATAAGCAACGACCCCCTG
GGAGTCTTAAGCGATTGGACCATTATTGGGAGTCTTCGTCATTGCAACTGGACCGGTATTACGTGCGAC
AGCACTGGTCACGTCGTGTCTGTTTCACTTTTGGAGAAGCAGTTGGAGGGAGTCTTGAGTCCAGCGATC
GCTAACCTTACTTATTTACAAGTCTTGGACCTTACAAGTAATAGTTTTACAGGGAAGATACCGGCGGAAA
TTGGTAAACTTACGGAGCTTAATCAGTTGATTCTGTATCTTAACTACTTCAGCGGATCTATTCCATCAGGA
ATCTGGGAACTGAAGAATATATTTTATTTAGACTTGCGTAATAACCTACTATCCGGTGATGTCCCAGAGG
AGATATGTAAGACCTCCAGCTTAGTTCTTATTGGGTTTGACTACAATAATCTTACAGGCAAGATCCCGGA
GTGTTTAGGGGACCTTGTGCATCTGCAGATGTTTGTCGCCGCAGGGAATCATTTGACCGGAAGTATACC
TGTGAGTATTGGGACATTGGCAAATCTGACCGATCTAGATCTTTCAGGCAATCAGTTGACTGGAAAAAT
ACCGAGAGATTTCGGTAACTTACTTAATCTTCAGTCTCTTGTACTGACCGAGAATCTGCTAGAGGGAGAC
ATACCGGCTGAGATAGGAAACTGCTCTTCCTTAGTCCAGTTAGAATTATATGACAACCAACTGACAGGCA
AAATCCCTGCTGAACTAGGAAACCTAGTACAACTACAAGCGTTGCGTATTTACAAAAACAAACTAACATC
TTCAATACCATCAAGCCTGTTTCGTCTGACTCAGCTAACTCATCTAGGTCTTAGCGAGAACCACCTGGTCG
GACCTATCAGTGAAGAGATCGGTTTCCTTGAAAGCCTTGAAGTTCTTACCTTACATTCCAACTCTTTCACC
GGGGAGTTCCCACAGAGTATCACAAATTTACGTAACTTGACTGTGTTGACTGTTGGCTTCAACTCTATCA
GTGGCGAATTGCCTGCTGATCTAGGGCTGTTAACGAATCTGCGTTCTTTAAGCGCGCATGATAATTTGCT
TACAGGCCCTATACCGTCCAGTATCTCAAATTGTACAGGCCTTAAATTATTGGACCTGTCTCACAATCAAA
TGACCGGCGAAATCCCACGTGGATTTGGCAGGATGAACCTTACGTTCATATCAATAGGCCGTAACCATTT
CACCGGCGAAATACCCGATGATATCTTTAATTGTAGTAATTTGGAGACCTTATCTGTGGCGGATAACAAC
CTTACTGGTACCCTTAAACCCCTAATCGGAAAGCTGCAAAAGCTTAGAATCCTACAGGTCTCCTACAATT
CATTAACCGGGCCTATCCCCAGGGAGATAGGGAACTTGAAGGACTTAAACATCCTTTATCTACACTCAAA
TGGTTTCACAGGTAGGATACCCAGAGAGATGAGCAATTTGACATTGCTACAAGGGTTGAGAATGTACTC

CAATGACTTGGAAGGCCCGATTCCCGAGGAAATGTTCGACATGAAGCTGTTGTCTGTGCTGGATCTTAG
CAATAATAAGTTTTCAGGCCAAATCCCAGCATTGTTTTCTAAATTAGAGTCACTTACATATCTGTCTCTTCA
AGGTAACAAGTTTAATGGCAGTATACCTGCAAGCTTGAAATCCTTGTCTTTATTGAACACATTCGATATA
AGTGACAATCTATTAACGGGAACGATTCCCGGTGAACTTCTTGCAAGCCTAAAGAACATGCAACTATATC
TGAACTTCTCTAACAACTTATTAACGGGGACTATTCCTAAGGAACTAGGCAAATTAGAGATGGTGCAAG
AAATAGACTTATCAAACAACTTGTTTTCAGGATCAATACCGAGGTCCTTGCAGGCATGTAAAAACGTCTT
TACGCTAGATTTTAGCCAGAACAACTTGTCCGGTCACATTCCTGACGAAGTTTTCCAGGGTATGGACATG
ATAATTTCCTTAAACTTGTCCAGAAACTCATTCTCCGGGGAGATCCCTCAGTCCTTTGGTAATATGACGCA
TCTAGTGAGCTTGGATCTTTCATCTAATAACCTAACTGGAGAGATCCCCGAGTCATTAGCAAATTTGAGC
ACGTTAAAACATCTAAAGCTTGCAAGCAATAATCTTAAAGGACATGTACCGGAGTCCGGAGTTTTTAAG
AATATTAACGCCAGTGACTTAATGGGTAACACGGATCTTTGTGGCAGTAAGAAGCCTCTAAAGCCATGC
ACCATAAAACAGAAATCCAGCCACTTCAGTAAGAGGACTCGT

*S. cerevisiae codon optimized FLS2LRR-Var4 insert sequence:*

AAACAATCATTTGAGCCAGAGATCGAAGCACTTAAGTCCTTCAAAAATGGCATAAGCAACGACCCCCTG
GGAGTCTTAAGCGATTGGACCATTATTGGGAGTCTTCGTCATTGCAACTGGACCGGTATTACGTGCGAC
AGCACTGGTCACGTCGTGTCTGTTTCACTTTTGGAGAAGCAGTTGGAGGGAGTCTTGAGTCCAGCGATC
GCTAACCTTACTTATTTACAAGTCTTGGACCTTACAAGTAATAGTTTTACAGGGAAGATACCGGCGGAAA
TTGGTAAACTTACGGAGCTTAATCAGTTGATTCTGTATCTTAACTACTTCAGCGGATCTATTCCATCAGGA
ATCTGGGAACTGAAGAATATATTTTATTTAGACTTGCGTAATAACCTACTATCCGGTGATGTCCCAGAGG
AGATATGTAAGACCTCCAGCTTAGTTCTTATTGGGTTTGACTACAATAATCTTACAGGCAAGATCCCGGA
GTGTTTAGGGGACCTTGTGCATCTGCAGATGTTTGTCGCCGCAGGGAATCATTTGACCGGAAGTATACC
TGTGAGTATTGGGACATTGGCAAATCTGACCGATCTAGATCTTTCAGGCAATCAGTTGACTGGAAAAAT
ACCGAGAGATTTCGGTAACTTACTTAATCTTCAGTCTCTTGTACTGACCGAGAATCTGCTAGAGGGAGAC
ATACCGGCTGAGATAGGAAACTGCTCTTCCTTAGTCCAGTTAGAATTATATGACAACCAACTGACAGGCA
AAATCCCTGCTGAACTAGGAAACCTAGTACAACTACAAGCGTTGCGTATTTACAAAAACAAACTAACATC
TTCAATACCATCAAGCCTGTTTCGTCTGACTCAGCTAACTCATCTAGGTCTTAGCGAGAACCACCTGGTCG
GACCTATCAGTGAAGAGATCGGTTTCCTTGAAAGCCTTGAAGTTCTTACCTTACATTCCAACTCTTTCACC
GGGGAGTTCCCACAGAGTATCACAAATTTACGTAACTTGACTGTGTTGACTGTTGGCTTCAACTCTATCA
GTGGCGAATTGCCTGCTGATCTAGGGCTGTTAACGAATCTGCGTTCTTTAAGCGCGCATGATAATTTGCT
TACAGGCCCTATACCGTCCAGTATCTCAAATTGTACAGGCCTTAAATTATTGGACCTGTCTCACAATCAAA
TGACCGGCGAAATCCCACGTGGATTTGGCAGGATGAACCTTACGTTCATATCAATAGGCCGTAACCATTT
CACCGGCGAAATACCCGATGATATCTTTAATTGTAGTAATTTGGAGACCTTATCTGTGGCGGATAACAAC
CTTACTGGTACCCTTAAACCCCTAATCGGAAAGCTGCAAAAGCTTAGAATCCTACAGGTCTCCTACAATT
CATTAACCGGGCCTATCCCCAGGGAGATAGGGAACTTGAAGGACTTAAACATCCTTTATCTACACTCAAA
TGGTTTCACAGGTAGGATACCCAGAGAGATGAGCAATTTGACATTGCTACAAGGGTTGAGAATGTACTC

CAATGACTTGGAAGGCCCGATTCCCGAGGAAATGTTCGACATGAAGCTGTTGTCTGTGCTGGATCTTAG
CAATAATAAGTTTTCAGGCCAAATCCCAGCATTGTTTTCTAAATTAGAGTCACTTACATATCTGTCTCTTCA
AGGTAACAAGTTTAATGGCAGTATACCTGCAAGCTTGAAATCCTTGTCTTTATTGAACACATTCGATATA
AGTGACAATCTATTAACGGGAACGATTCCCGGTGAACTTCTTGCAAGCCTAAAGAACATGCAACTATATC
TGTCTTTCTCTAACAACTTATTAACGGGGACTATTCCTAAGGAACTAGGCAAATTAGAGATGGTGCAAGA
AATAGACTTATCAAACAACTTGTTTTCAGGATCAATACCGAGGTCCTTGCAGGCATGTAAAAACGTCTTT
ACGCTAGATTTTAGCCAGAACAACTTGTCCGGTCACATTCCTGACGAAGTTTTCCAGGGTATGGACATGA
TAATTTCCTTATCTTTGTCCAGAAACTCATTCTCCGGGGAGATCCCTCAGTCCTTTGGTAATATGACGCAT
CTAGTGAGCTTGGATCTTTCATCTAATAACCTAACTGGAGAGATCCCCGAGTCATTAGCAAATTTGAGCA
CGTTAAAACATCTAAAGCTTGCAAGCAATAATCTTAAAGGACATGTACCGGAGTCCGGAGTTTTTAAGA
ATATTAACGCCAGTGACTTAATGGGTAACACGGATCTTTGTGGCAGTAAGAAGCCTCTAAAGCCATGCA
CCATAAAACAGAAATCCAGCCACTTCAGTAAGAGGACTCGT

**Supplementary information 3. pCT80-FnLoopHP surface display expression vector DNA sequence**. This sequence was originally described in Stern, L.A., et al. 2017 (13). Highlighted regions within the open reading frame are colored as follows: red for start codon; yellow for Aga2; magenta for HA-tag; Green for the NheI restriction enzyme cut site; Blue for the BamHI restriction enzyme cut site; grey for two stop codons. Any above insert sequence can be Gibson cloned between the NheI and BamHI cut sites using the homologous overlapping primers provided in **Supplmentary Table 1**.

**PCT80-FnLoopHp Vector:**

ACGAAAGGGCCTCGTGATACGCCTATTTTTATAGGTTAATGTCATGATAATAATGGTTTCTTAGACGGAT
CGCTTGCCTGTAACTTACACGCGCCTCGTATCTTTTAATGATGGAATAATTTGGGAATTTACTCTGTGTTT
ATTTATTTTTATGTTTTGTATTTGGATTTTAGAAAGTAAATAAAGAAGGTAGAAGAGTTACGGAATGAAG
AAAAAAAAATAAACAAAGGTTTAAAAAATTTCAACAAAAAGCGTACTTTACATATATATTTATTAGACAA
GAAAAGCAGATTAAATAGATATACATTCGATTAACGATAAGTAAAATGTAAAATCACAGGATTTTCGTGT
GTGGTCTTCTACACAGACAAGATGAAACAATTCGGCATTAATACCTGAGAGCAGGAAGAGCAAGATAA

AAGGTAGTATTTGTTGGCGATCCCCCTAGAGTCTTTTACATCTTCGGAAAACAAAAACTATTTTTTCTTTA
ATTTCTTTTTTTACTTTCTATTTTTAATTTATATATTTATATTAAAAAATTTAAATTATAATTATTTTTATAGC
ACGTGATGAAAAGGACCCAGGTGGCACTTTTCGGGGAAATGTGCGCGGAACCCCTATTTGTTTATTTTTC
TAAATACATTCAAATATGTATCCGCTCATGAGACAATAACCCTGATAAATGCTTCAATAATATTGAAAAA
GGAAGAGTATGAGTATTCAACATTTCCGTGTCGCCCTTATTCCCTTTTTTGCGGCATTTTGCCTTCCTGTTT
TTGCTCACCCAGAAACGCTGGTGAAAGTAAAAGATGCTGAAGATCAGTTGGGTGCACGAGTGGGTTAC
ATCGAACTGGATCTCAACAGCGGTAAGATCCTTGAGAGTTTTCGCCCCGAAGAACGTTTTCCAATGATGA
GCACTTTTAAAGTTCTGCTATGTGGCGCGGTATTATCCCGTATTGACGCCGGGCAAGAGCAACTCGGTC
GCCGCATACACTATTCTCAGAATGACTTGGTTGAGTACTCACCAGTCACAGAAAAGCATCTTACGGATGG
CATGACAGTAAGAGAATTATGCAGTGCTGCCATAACCATGAGTGATAACACTGCGGCCAACTTACTTCT
GACAACGATCGGAGGACCGAAGGAGCTAACCGCTTTTTTGCACAACATGGGGGATCATGTAACTCGCCT
TGATCGTTGGGAACCGGAGCTGAATGAAGCCATACCAAACGACGAGCGTGACACCACGATGCCTGTAG
CAATGGCAACAACGTTGCGCAAACTATTAACTGGCGAACTACTTACTCTAGCTTCCCGGCAACAATTAAT
AGACTGGATGGAGGCGGATAAAGTTGCAGGACCACTTCTGCGCTCGGCCCTTCCGGCTGGCTGGTTTAT
TGCTGATAAATCTGGAGCCGGTGAGCGTGGGTCTCGCGGTATCATTGCAGCACTGGGGCCAGATGGTA
AGCCCTCCCGTATCGTAGTTATCTACACGACGGGGAGTCAGGCAACTATGGATGAACGAAATAGACAGA
TCGCTGAGATAGGTGCCTCACTGATTAAGCATTGGTAACTGTCAGACCAAGTTTACTCATATATACTTTA
GATTGATTTAAAACTTCATTTTTAATTTAAAAGGATCTAGGTGAAGATCCTTTTTGATAATCTCATGACCA
AAATCCCTTAACGTGAGTTTTCGTTCCACTGAGCGTCAGACCCCGTAGAAAAGATCAAAGGATCTTCTTG
AGATCCTTTTTTTCTGCGCGTAATCTGCTGCTTGCAAACAAAAAAACCACCGCTACCAGCGGTGGTTTGTT
TGCCGGATCAAGAGCTACCAACTCTTTTTCCGAAGGTAACTGGCTTCAGCAGAGCGCAGATACCAAATA
CTGTTCTTCTAGTGTAGCCGTAGTTAGGCCACCACTTCAAGAACTCTGTAGCACCGCCTACATACCTCGCT
CTGCTAATCCTGTTACCAGTGGCTGCTGCCAGTGGCGATAAGTCGTGTCTTACCGGGTTGGACTCAAGAC
GATAGTTACCGGATAAGGCGCAGCGGTCGGGCTGAACGGGGGGTTCGTGCACACAGCCCAGCTTGGA
GCGAACGACCTACACCGAACTGAGATACCTACAGCGTGAGCTATGAGAAAGCGCCACGCTTCCCGAAG
GGAGAAAGGCGGACAGGTATCCGGTAAGCGGCAGGGTCGGAACAGGAGAGCGCACGAGGGAGCTTC
CAGGGGGAAACGCCTGGTATCTTTATAGTCCTGTCGGGTTTCGCCACCTCTGACTTGAGCGTCGATTTTT
GTGATGCTCGTCAGGGGGGCGGAGCCTATGGAAAAACGCCAGCAACGCGGCCTTTTTACGGTTCCTGG
CCTTTTGCTGGCCTTTTGCTCACATGTTCTTTCCTGCGTTATCCCCTGATTCTGTGGATAACCGTATTACCG
CCTTTGAGTGAGCTGATACCGCTCGCCGCAGCCGAACGACCGAGCGCAGCGAGTCAGTGAGCGAGGAA
GCGGAAGAGCGCCCAATACGCAAACCGCCTCTCCCCGCGCGTTGGCCGATTCATTAATGCAGCTGGCAC
GACAGGTTTCCCGACTGGAAAGCGGGCAGTGAGCGCAACGCAATTAATGTGAGTTAGCTCACTCATTAG
GCACCCCAGGCTTTACACTTTATGCTTCCGGCTCGTATGTTGTGTGGAATTGTGAGCGGATAACAATTTC
ACACAGGAAACAGCTATGACCATGATTACGCCAAGCTCGAAATTAACCCTCACTAAAGGGAACAAAAGC
TGGTACCAATTCCTTGAATTTTCAAAAATTCTTACTTTTTTTTTGGATGGACGCAAAGAAGTTTAATAATC
ATATTACATGGCATTACCACCATATACATATCCATATCTAATCTTACTTATATGTTGTGGAAATGTAAAGA
GCCCCATTATCTTAGCCTAAAAAAACCTTCTCTTTGGAACTTTCAGTAATACGCTTAACTGCTCATTGCTAT
ATTGAAGTACGGATTAGAAGCCGCCGAGCGGGTGACAGCCCTCCGAAGGAAGACTCTCCTCCGTGCGT

CCTCGTCTTCACCGGTCGCGTTCCTGAAACGCAGATGTGCCTCGCGCCGCACTGCTCCGAACAATAAAGA
TTCTACAATACTAGCTTTTATGGTTATGAAGAGGAAAAATTGGCAGTAACCTGGCCCCACAAACCTTCAA
ATGAACGAATCAAATTAACAACCATAGGATGATAATGCGATTAGTTTTTTAGCCTTATTTCTGGGGTAAT
TAATCAGCGAAGCGATGATTTTTGATCTATTAACAGATATATAAATGCAAAAACTGCATAACCACTTTAA
CTAATACTTTCAACATTTTCGGTTTGTATTACTTCTTATTCAAATGTAATAAAAGTATCAACAAAAAATTGT
TAATATACCTCTATACTTTAACGTCAAGGAGAAAAAACCCCGGATCGAATTCCCTACTTCATACATTTTCA
ATTAAGATGCAGTTACTTCGCTGTTTTTCAATATTTTCTGTTATTGCTTCAGTTTTAGCACAGGAACTGAC
AACTATATGCGAGCAAATCCCCTCACCAACTTTAGAATCGACGCCGTACTCTTTGTCAACGACTACTATTT
TGGCCAACGGGAAGGCAATGCAAGGAGTTTTTGAATATTACAAATCAGTAACGTTTGTCAGTAATTGCG
GTTCTCACCCCTCAACAACTAGCAAAGGCAGCCCCATAAACACACAGTATGTTTTTAAGGACAATAGCTC
GACGATTGAAGGTAGATACCCATACGACGTTCCAGACTACGCTCTGCAGGCTAGTGCCTCTCCAGCTGC
ACCTGCTCCAGCAAGCCCTGCTGCACCAGCTCCGTCTGCTCCTGCTGCCTCTCCAGCTGCACCTGCTCCAG
CTTCTCCAGCAGCTCCTGCACCTAGTGCTCCTGCTGGGGGTGGAGGCTCTGGCGGAGGTGGGTCTGGTG
GGGGCGGATCTGCTAGCTCCTCCGACTCTCCGCGTAACCTGGAGGTTACCAACGCAACTCCGAACTCTCT
GACTATTTCTTGGCCATGGATCTGCCCCGGGATGTACCATATGCCAATCAGCATCAATTATCGCACCGAA
ATCGACAAACCGTCTCAGGGATCCGAACAAAAGCTTATTTCTGAAGAGGACTTGTAATAGCTCGAGATC
TGATAACAACAGTGTAGATGTAACAAAATCGACTTTGTTCCCACTGTACTTTTAGCTCGTACAAAATACA
ATATACTTTTCATTTCTCCGTAAACAACATGTTTTCCCATGTAATATCCTTTTCTATTTTTCGTTCCGTTACC
AACTTTACACATACTTTATATAGCTATTCACTTCTATACACTAAAAAACTAAGACAATTTTAATTTTGCTGC
CTGCCATATTTCAATTTGTTATAAATTCCTATAATTTATCCTATTAGTAGCTAAAAAAAGATGAATGTGAA
TCGAATCCTAAGAGAATTGAGCTCCAATTCGCCCTATAGTGAGTCGTATTACAATTCACTGGCCGTCGTT
TTACAACGTCGTGACTGGGAAAACCCTGGCGTTACCCAACTTAATCGCCTTGCAGCACATCCCCCTTTCG
CCAGCTGGCGTAATAGCGAAGAGGCCCGCACCGATCGCCTTTCCCAACAGTTGCGCAGCCTGAATGGCG
AATGGACGCGCCCTGTAGCGGCGCATTAAGCGCGGCGGGTGTGGTGGTTACGCGCAGCGTGACCGCTA
CACTTGCCAGCGCCCTAGCGCCCGCTCCTTTCGCTTTCTTCCCTTCCTTTCTCGCCACGTTCGCCGGCTTTC
CCCGTCAAGCTCTAAATCGGGGGCTCCCTTTAGGGTTCCGATTTAGTGCTTTACGGCACCTCGACCCCAA
AAAACTTGATTAGGGTGATGGTTCACGTAGTGGGCCATCGCCCTGATAGACGGTTTTTCGCCCTTTGACG
TTGGAGTCCACGTTCTTTAATAGTGGACTCTTGTTCCAAACTGGAACAACACTCAACCCTATCTCGGTCTA
TTCTTTTGATTTATAAGGGATTTTGCCGATTTCGGCCTATTGGTTAAAAAATGAGCTGATTTAACAAAAAT
TTAACGCGAATTTTAACAAAATATTAACGCTTACAATTTCCTGATGCGGTATTTTCTCCTTACGCATCTGT
GCGGTATTTCACACCGCATAGATCGGCAAGTGCACAAACAATACTTAAATAAATACTACTCAGTAATAAC
CTATTTCTTAGCATTTTTGACGAAATTTGCTATTTTGTTAGAGTCTTTTACACCATTTGTCTCCACACCTCC
GCTTACATCAACACCAATAACGCCATTTAATCTAAGCGCATCACCAACATTTTCTGGCGTCAGTCCACCAG
CTAACATAAAATGTAAGCTTTCGGGGCTCTCTTGCCTTCCAACCCAGTCAGAAATCGAGTTCCAATCCAA
AAGTTCACCTGTCCCACCTGCTTCTGAATCAAACAAGGGAATAAACGAATGAGGTTTCTGTGAAGCTGC
ACTGAGTAGTATGTTGCAGTCTTTTGGAAATACGAGTCTTTTAATAACTGGCAAACCGAGGAACTCTTGG
TATTCTTGCCACGACTCATCTCCATGCAGTTGGACGATATCAATGCCGTAATCATTGACCAGAGCCAAAA
CATCCTCCTTAGGTTGATTACGAAACACGCCAACCAAGTATTTCGGAGTGCCTGAACTATTTTTATATGCT

TTTACAAGACTTGAAATTTTCCTTGCAATAACCGGGTCAATTGTTCTCTTTCTATTGGGCACACATATAAT
ACCCAGCAAGTCAGCATCGGAATCTAGAGCACATTCTGCGGCCTCTGTGCTCTGCAAGCCGCAAACTTTC
ACCAATGGACCAGAACTACCTGTGAAATTAATAACAGACATACTCCAAGCTGCCTTTGTGTGCTTAATCA
CGTATACTCACGTGCTCAATAGTCACCAATGCCCTCCCTCTTGGCCCTCTCCTTTTCTTTTTTCGACCGAAT
TAATTCTTAATCGGCAAAAAAAGAAAAGCTCCGGATCAAGATTGTACGTAAGGTGACAAGCTATTTTTCA
ATAAAGAATATCTTCCACTACTGCCATCTGGCGTCATAACTGCAAAGTACACATATATTACGATGCTGTTC
TATTAAATGCTTCCTATATTATATATATAGTAATGTCGTGATCTATGGTGCACTCTCAGTACAATCTGCTCT
GATGCCGCATAGTTAAGCCAGCCCCGACACCCGCCAACACCCGCTGACGCGCCCTGACGGGCTTGTCTG
CTCCCGGCATCCGCTTACAGACAAGCTGTGACCGTCTCCGGGAGCTGCATGTGTCAGAGGTTTTCACCGT
CATCACCGAAACGCGCGAG

**Supplementary Information 4. Example gating and calculation of background subtracted fraction binding.** Using the example gating strategy (left) in FCS Express 7, event count values are extracted and used for calculating the fraction binding (x) value described as the mean of triplicates change in conditional probabilities. In other words, this calculation determines how much more often binding occurs among expressing cells than non-expressing cells.

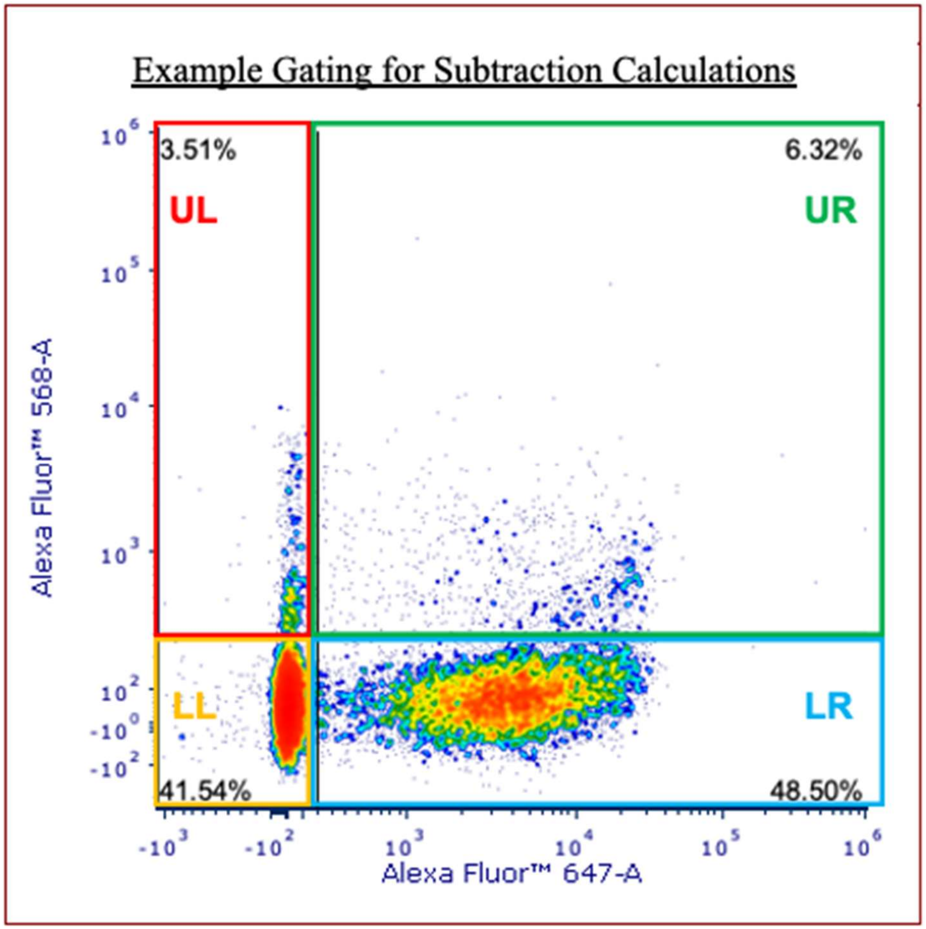

To determine specific ligand binding normalized to expression and subtract background derived from non-expressing cells, calculations are derived from the following formula:

$$\Delta\ in\ conditional\ probabilities = P(568+\mid 647+) - P(568+\mid 647-)$$

Where the conditional probability of ligand positive (568+) among non-expressing cells (647-) is subtracted from the conditional probability of ligand positive among expressing cells (647+). This is used to determine how much more often binding occurs among expressing cells than among non-expressing cells. Using bivariate density plots after flow cytometry gated based on unstained controls, the formula can be applied as follows:

$$\Delta\ in\ conditional\ probabilities = \left(\frac{UR}{UR+LR}\right) - \left(\frac{UL}{UL+LL}\right) = x$$

Where UR is the 568+/647+ event count, LR is the 568-/647+ event count, UL is the 568+/647- event count, and LL is the 568-/647- event count. For each replicate (n=3), x is calculated, and the mean of replicates is plotted as bar plots for each sample group. The standard deviation of replicate x values for each sample group is used to calculate the standard error to obtain error bars.

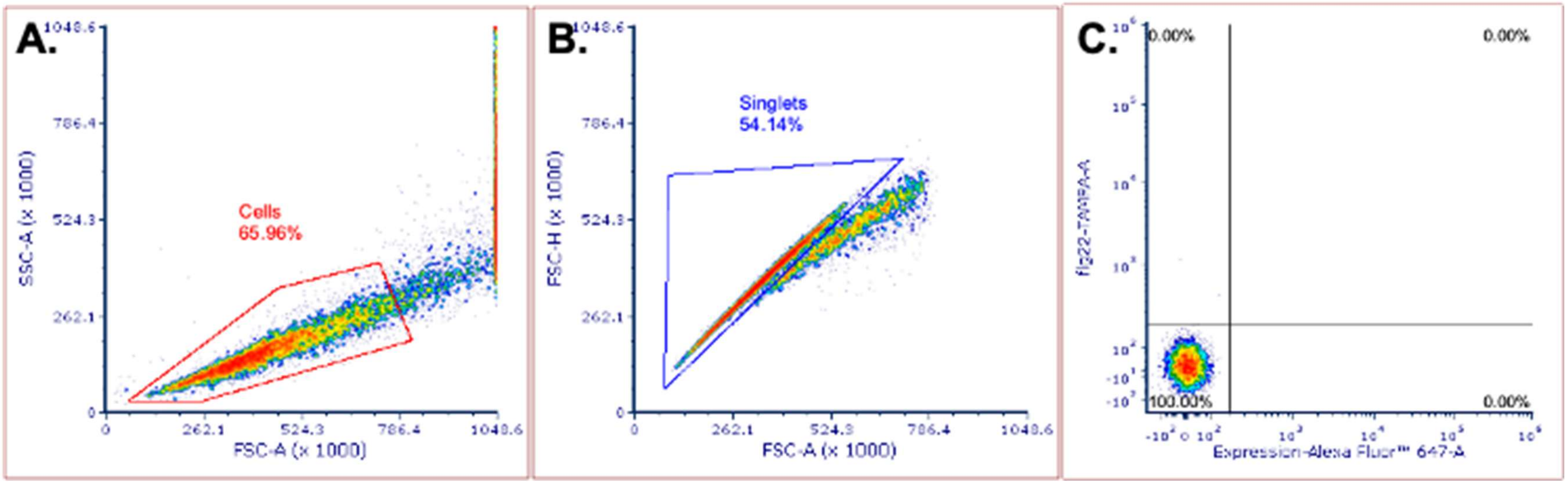

**Supplementary Figure 12: Gating strategy for flow cytometry experiments.** Flow cytometry was performed using either the Accuri C6 or the Attune Cytpix. Analysis of flow cytometry data was performed using FCS Express 7 Research. For each experiment, gating was prepared using the unstained sample. First, (A) density plot is prepared where Y-axis is set to Side Scatter – Area (SSC-A) and X-axis is set to Forward Scatter - Area (FSC-A), then polygon gate is drawn to select cells (Red). (B) ”Cells” gate is then copied to new density plot where Y-axis is set to Forward Scatter – Height (FSC-H) and X-axis is kept the same, then polygon gate is drawn to select for singlet cells (Blue). (C) The final density plot is prepared by copying ”Singlets” to new density plot where Y-axis is set to laser capable of detecting TAMRA with biexponential scaling and X-axis is set to laser capable of detecting Alexa Fluor 647 with biexponential scaling. Quadrant gates are drawn so that the double negative (unstained) population is in the lower left quadrant and spill over into other quadrants is below 0.1%.

**Supplementary Information 5a-q. Raw flow cytometry density plots.** Each raw density plot below is acquired using the gating strategy described in **Figure S12.** Each page of plots is organized in relation to what figure the plots are associated with from the main text and sample group (e.g. FLS2LRR). Some pages include multiple sample groups. Each sub-header is organized by figure number/panels, sample group, and treatment conditions from left to right.

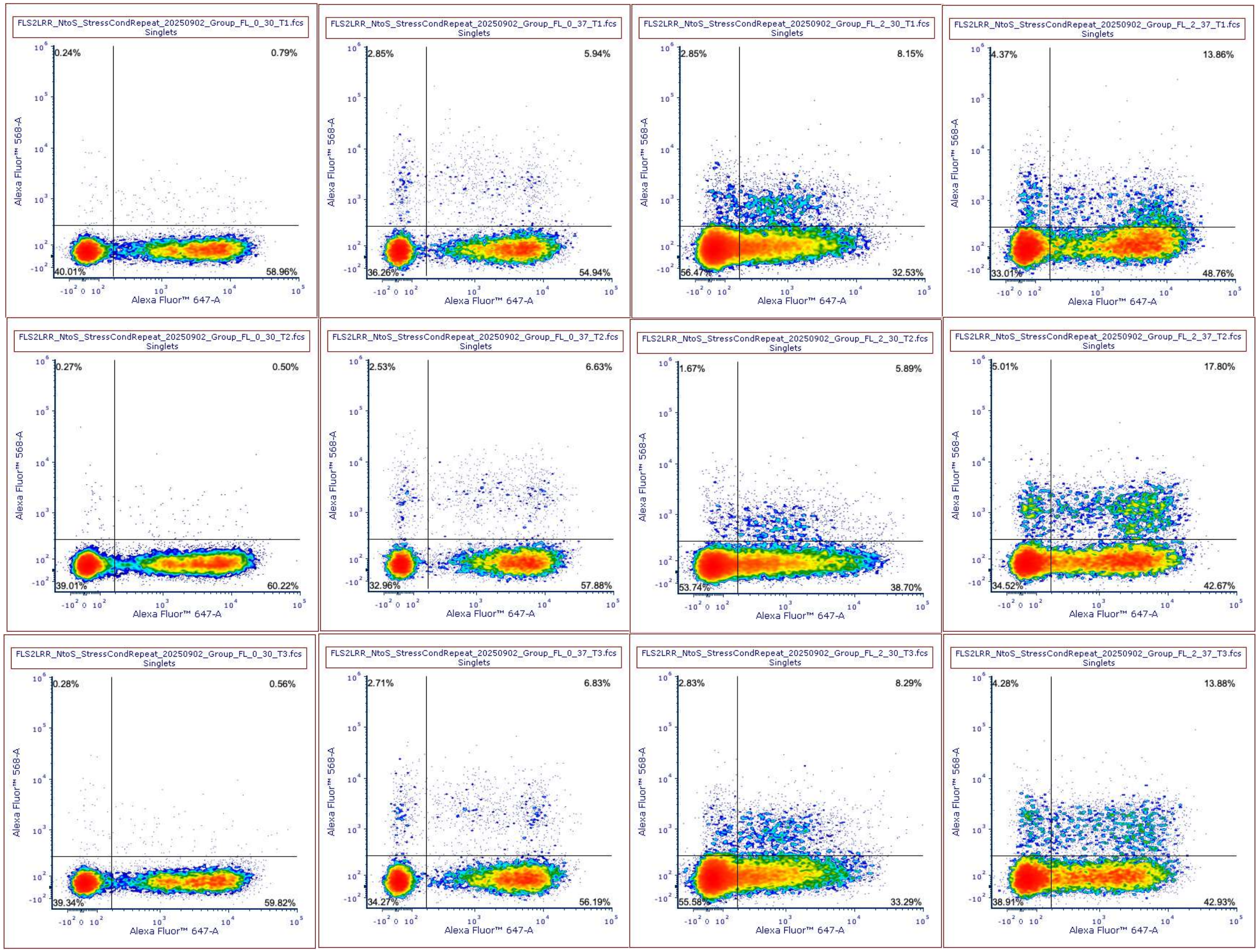

**Suppl. Inf. 5a - Figure 3a-b: FLS2LRR (0,30; 0,37; 2,30; 2,37 column order from left to right)**

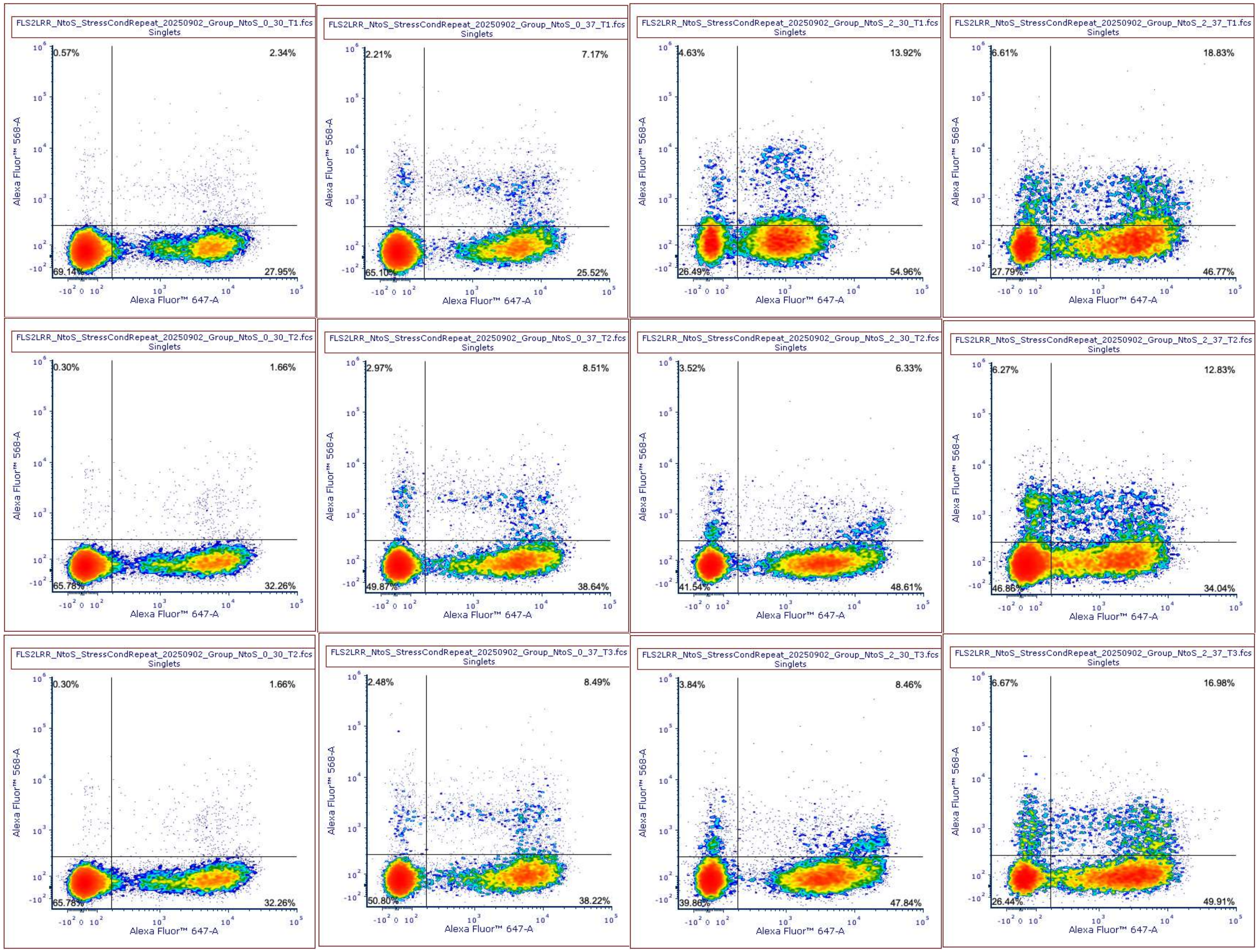

**Suppl. Inf. 5b - Figure 3c-d: NtoS (0,30; 0,37; 2,30; 2,37 column order from left to right)**

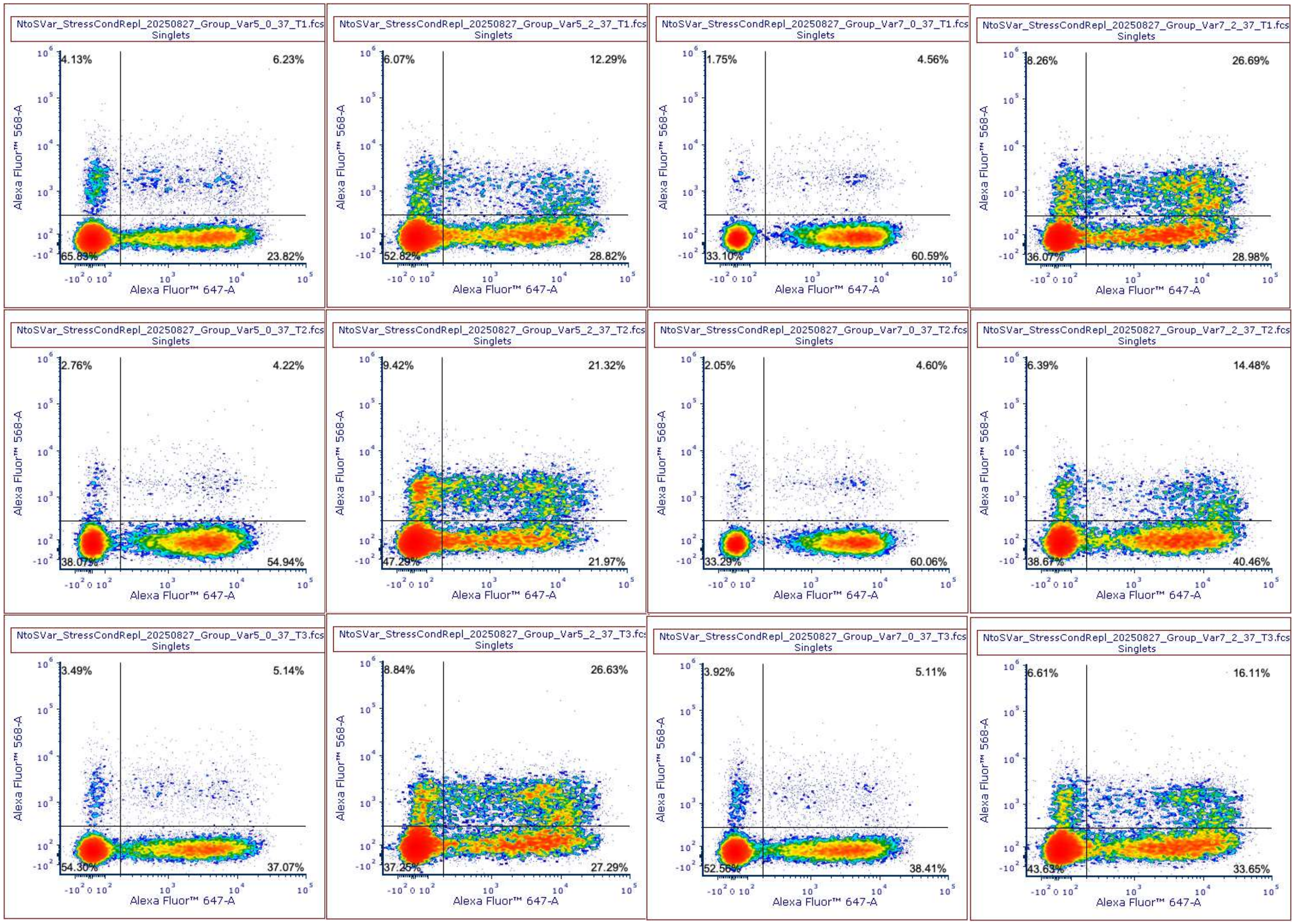

**Suppl. Inf. 5c - Figure 5: Var3 (first two columns) and Var4 (last two columns) - (0,37; 2,37 column order from left to right)**

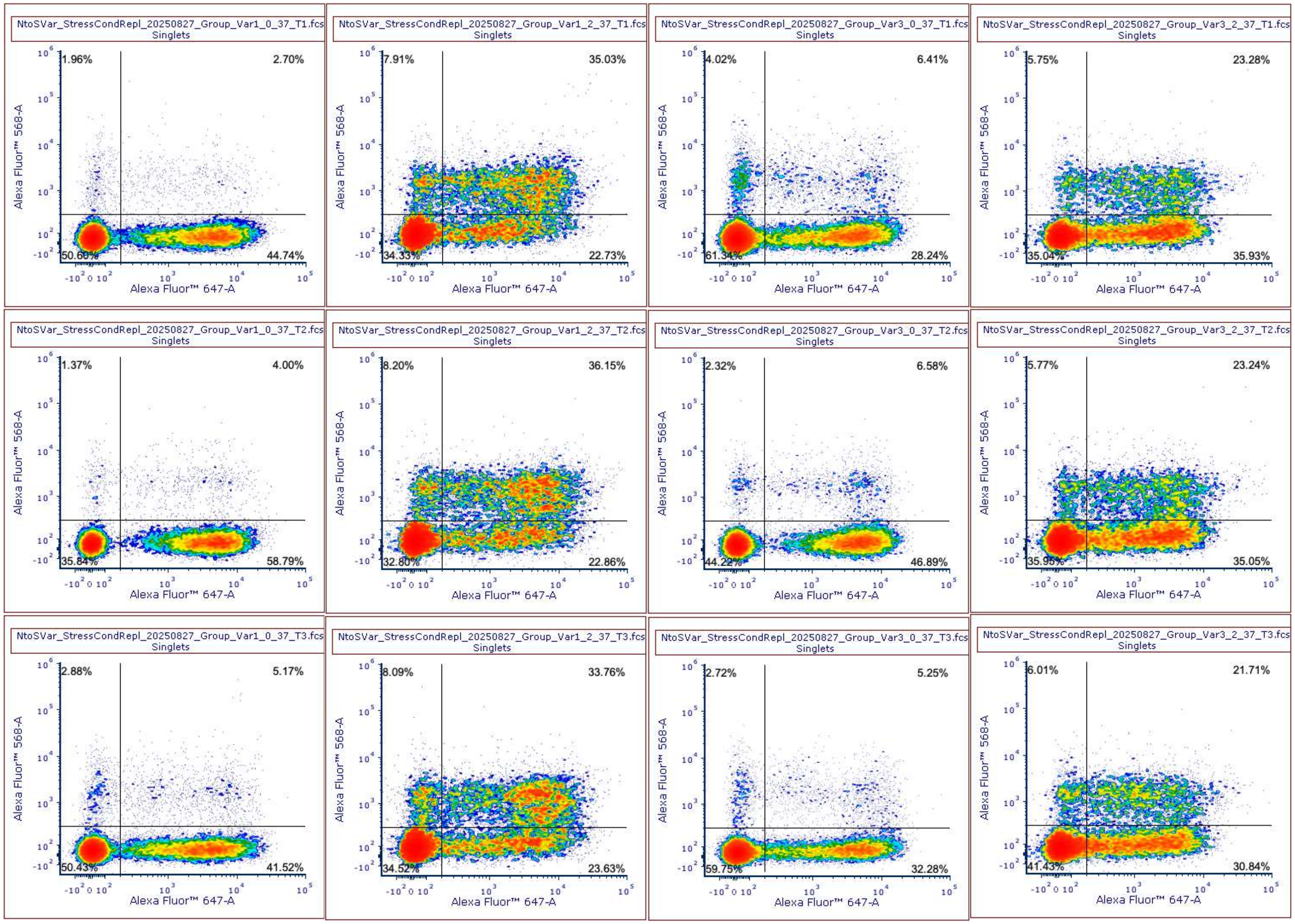

**Suppl. Inf. 5d - Figure 5: Var1 (first two columns) and Var2 (last two columns) - (0,37; 2,37 column order from left to right)**

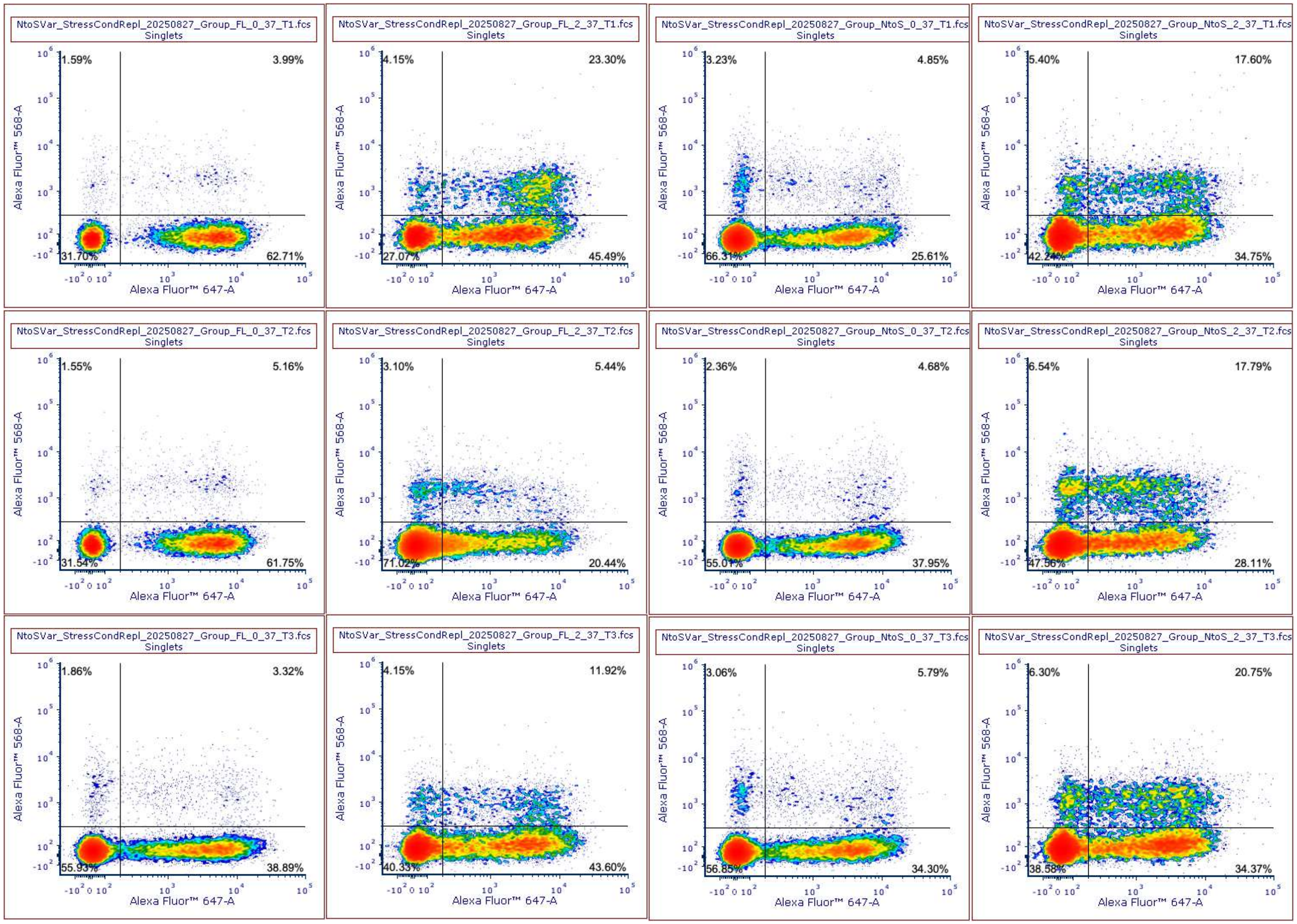

**Suppl. Inf. 5e - Figure 5 Related: FLS2LRR (first two columns) and NtoS (last two columns) - (0,37; 2,37 column order from left to right)**

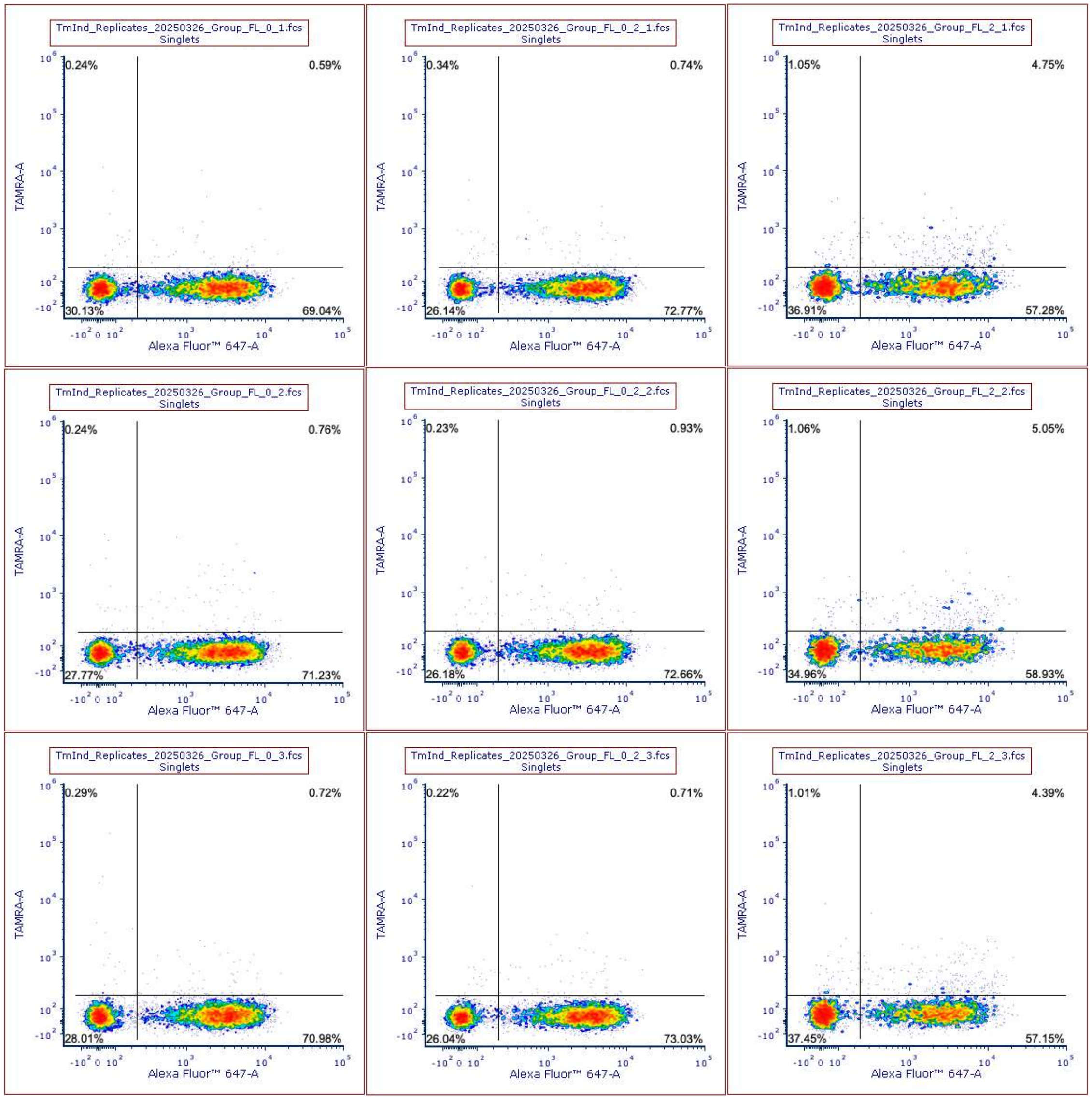

**Suppl. Inf. 5f - Figure 2a: FLS2LRR - (0; 0.2; 2 ug/mL tunicamycin column order from left to right)**

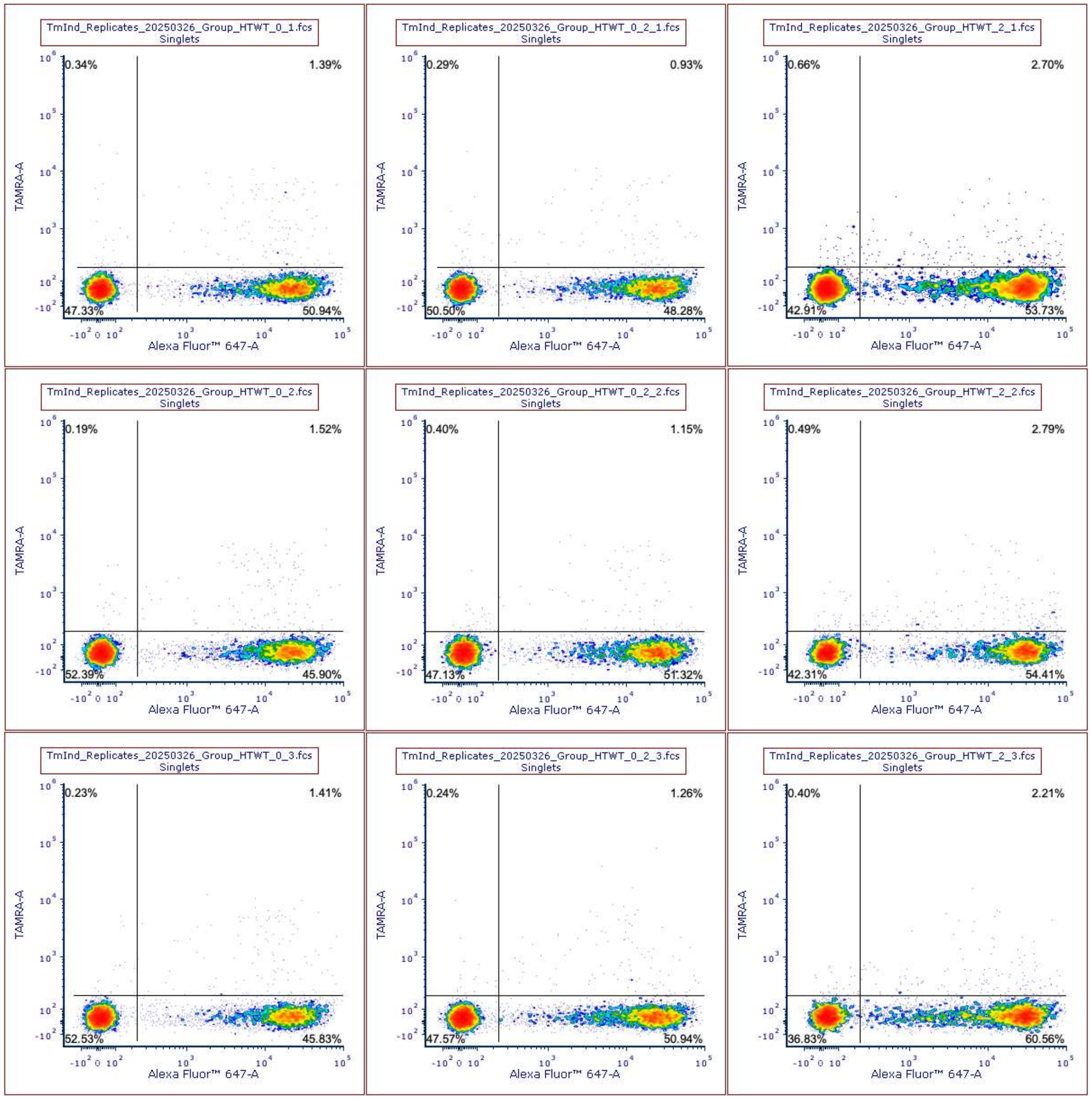

**Suppl. Inf. 5g - Figure 2a: HTWT - (0; 0.2; 2 ug/mL tunicamycin column order from left to right)**

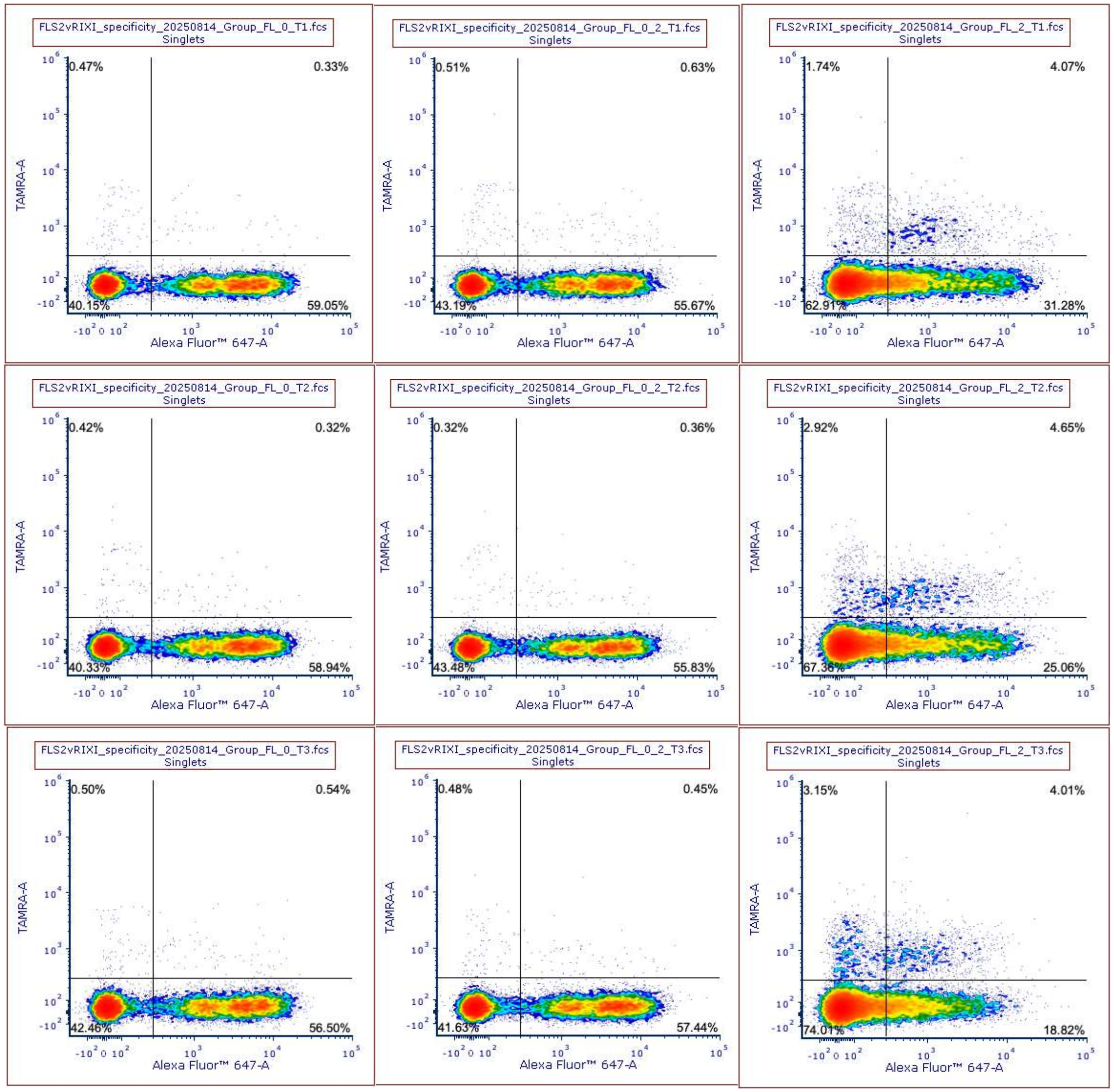

**Suppl. Inf. 5h - Figure 2b: FLS2LRR - (0; 0.2; 2 ug/mL tunicamycin column order from left to right)**

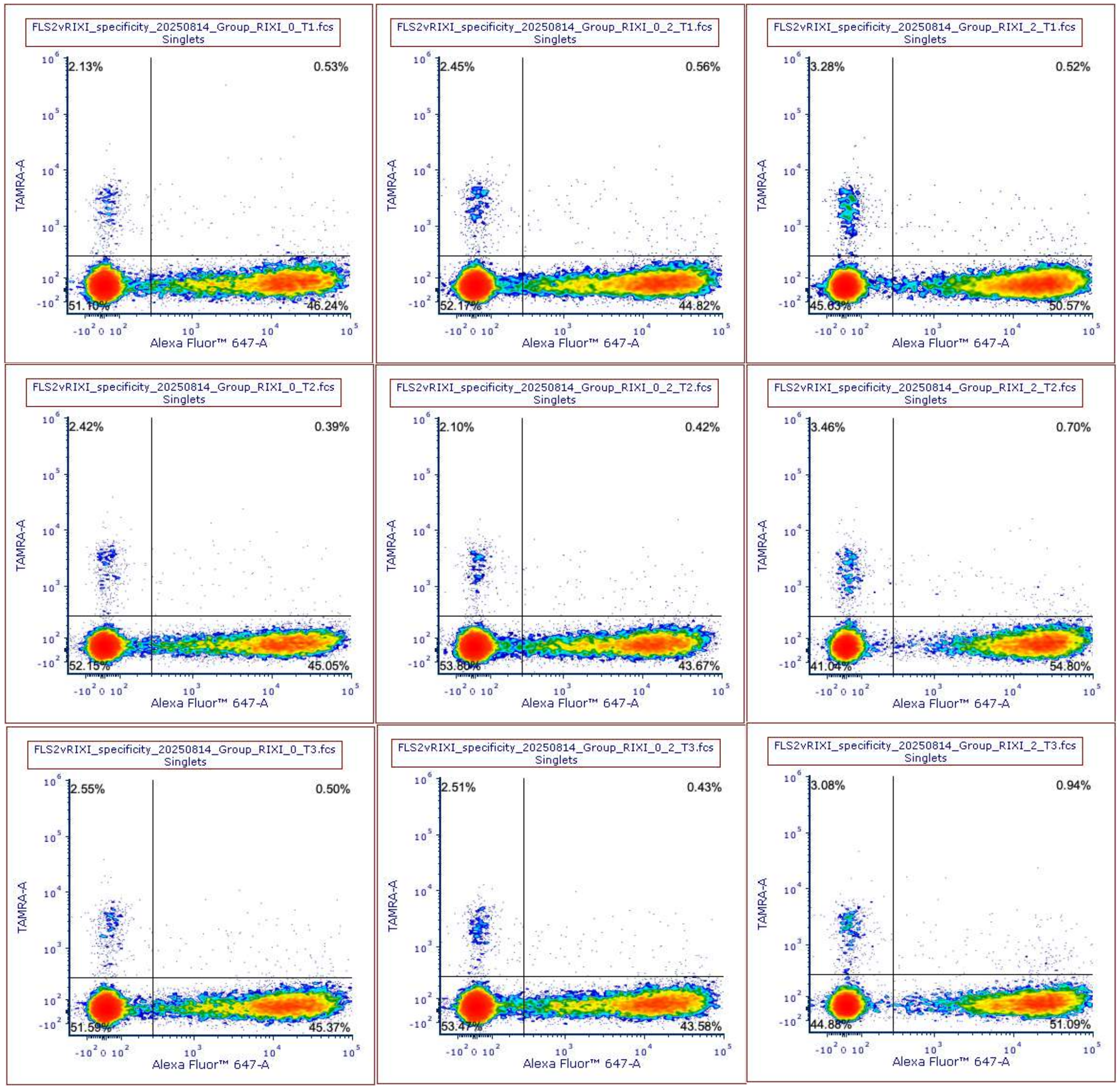

**Suppl. Inf. 5i - Figure 2b: RIXI - (0; 0.2; 2 u g /mL tunicamycin column order from left to right)**

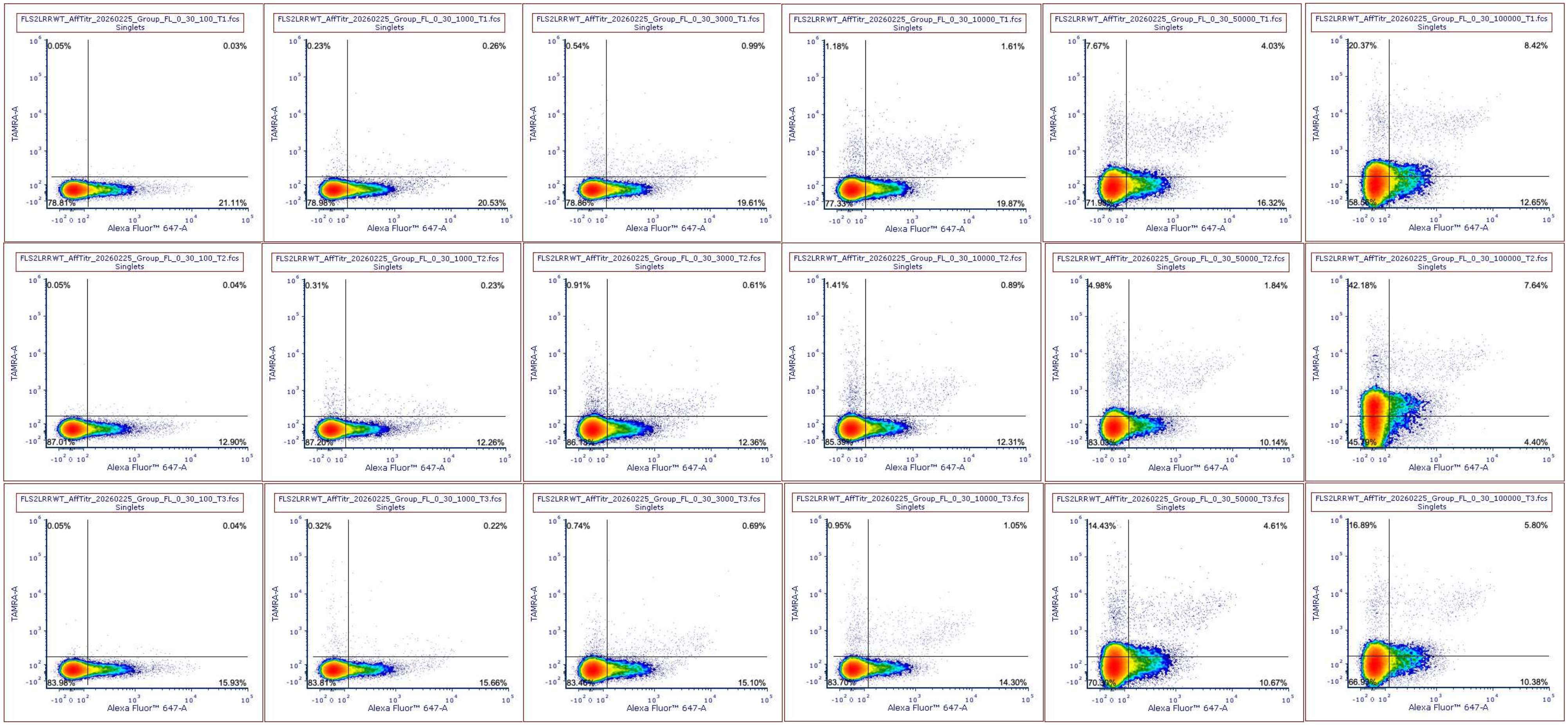

**Suppl. Inf. 5j - Figure 4: FLS2LRR - (No Stress – 0.1, 1, 3, 10, 50, 100 µM flg22-TAMRA column order from left to right)**

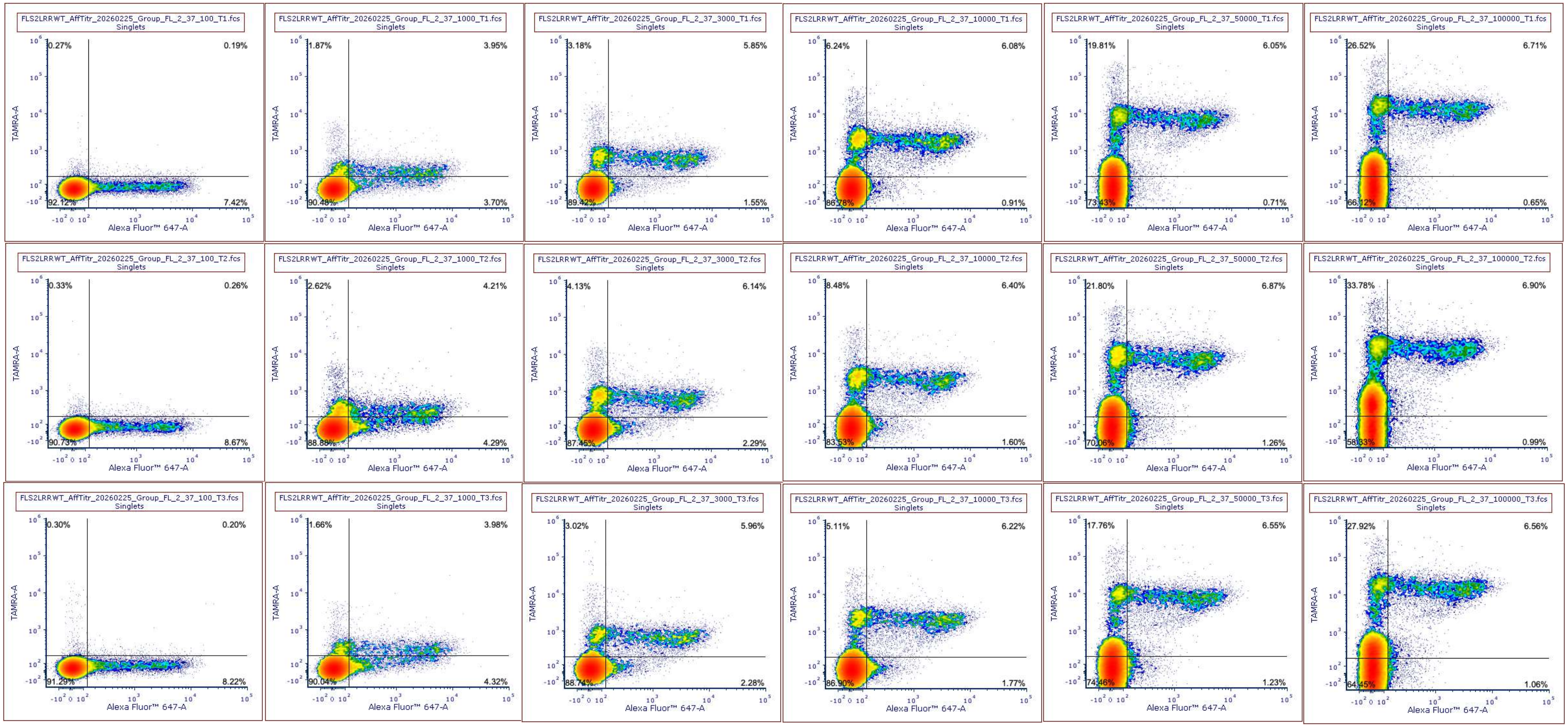

**Suppl. Inf. 5k - Figure 4: FLS2LRR - (Additive Stress – 0.1, 1, 3, 10, 50, 100 µM flg22-TAMRA column order from left to right)**

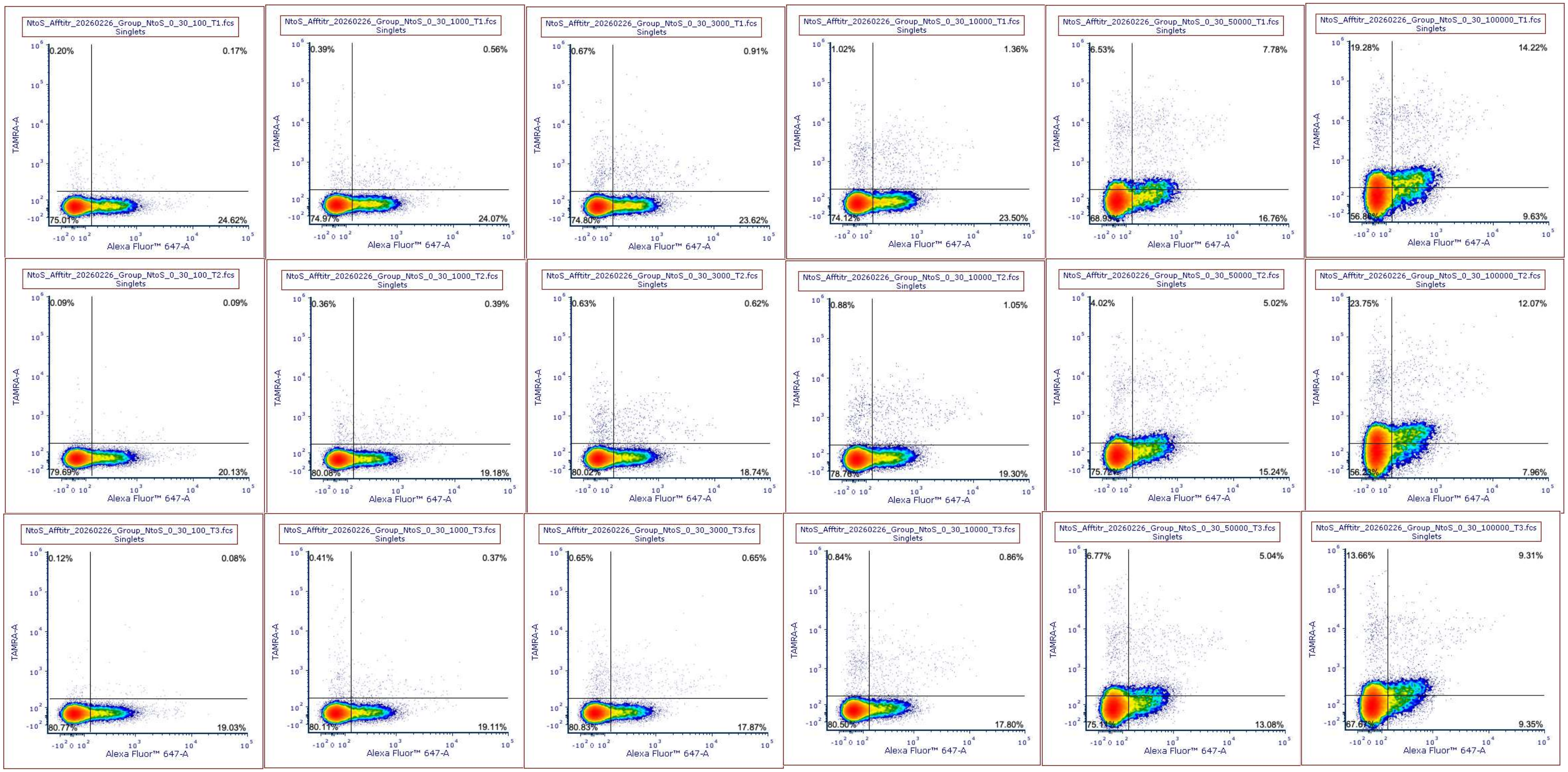

**Suppl. Inf. 5l - Figure 4: NtoS - (No Stress – 0.1, 1, 3, 10, 50, 100 µM flg22-TAMRA column order from left to right)**

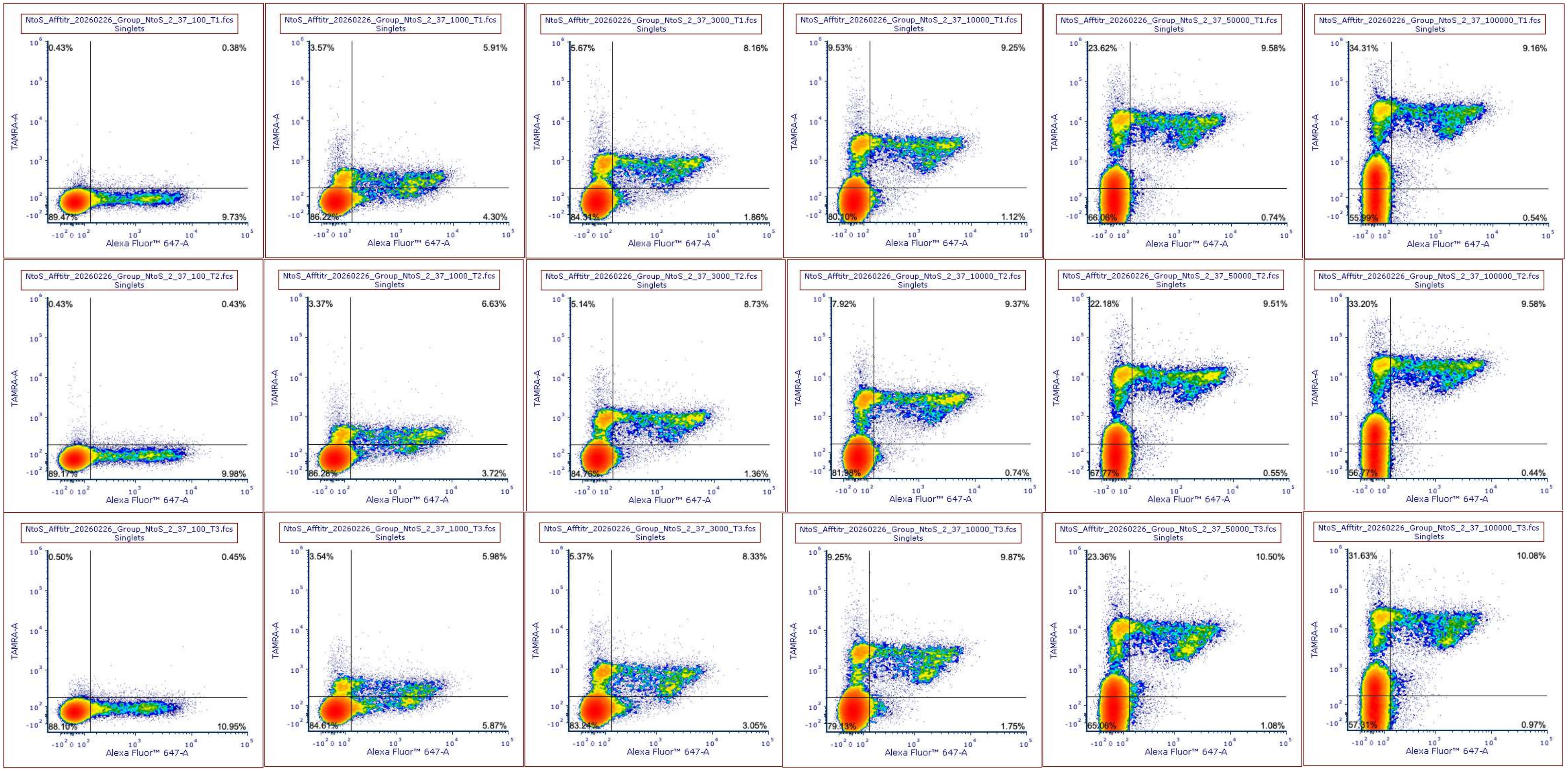

**Suppl. Inf. 5m - Figure 4: NtoS - (Additive Stress – 0.1, 1, 3, 10, 50, 100 µM flg22-TAMRA column order from left to right)**

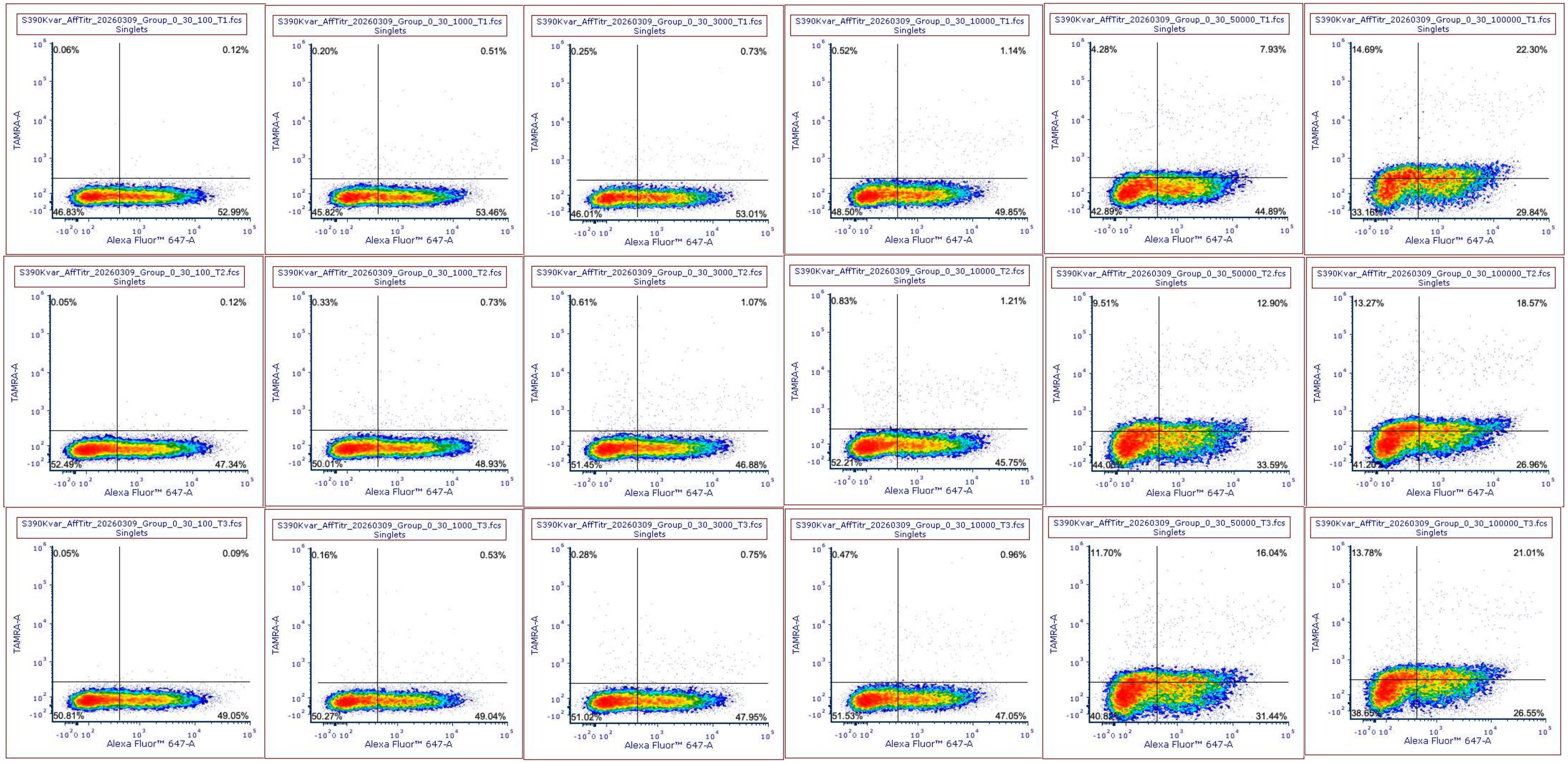

**Suppl. Inf. 5n - Figure 4: FLS2LRR-S390K - (No Stress – 0.1, 1, 3, 10, 50, 100 µM flg22-TAMRA column order from left to right)**

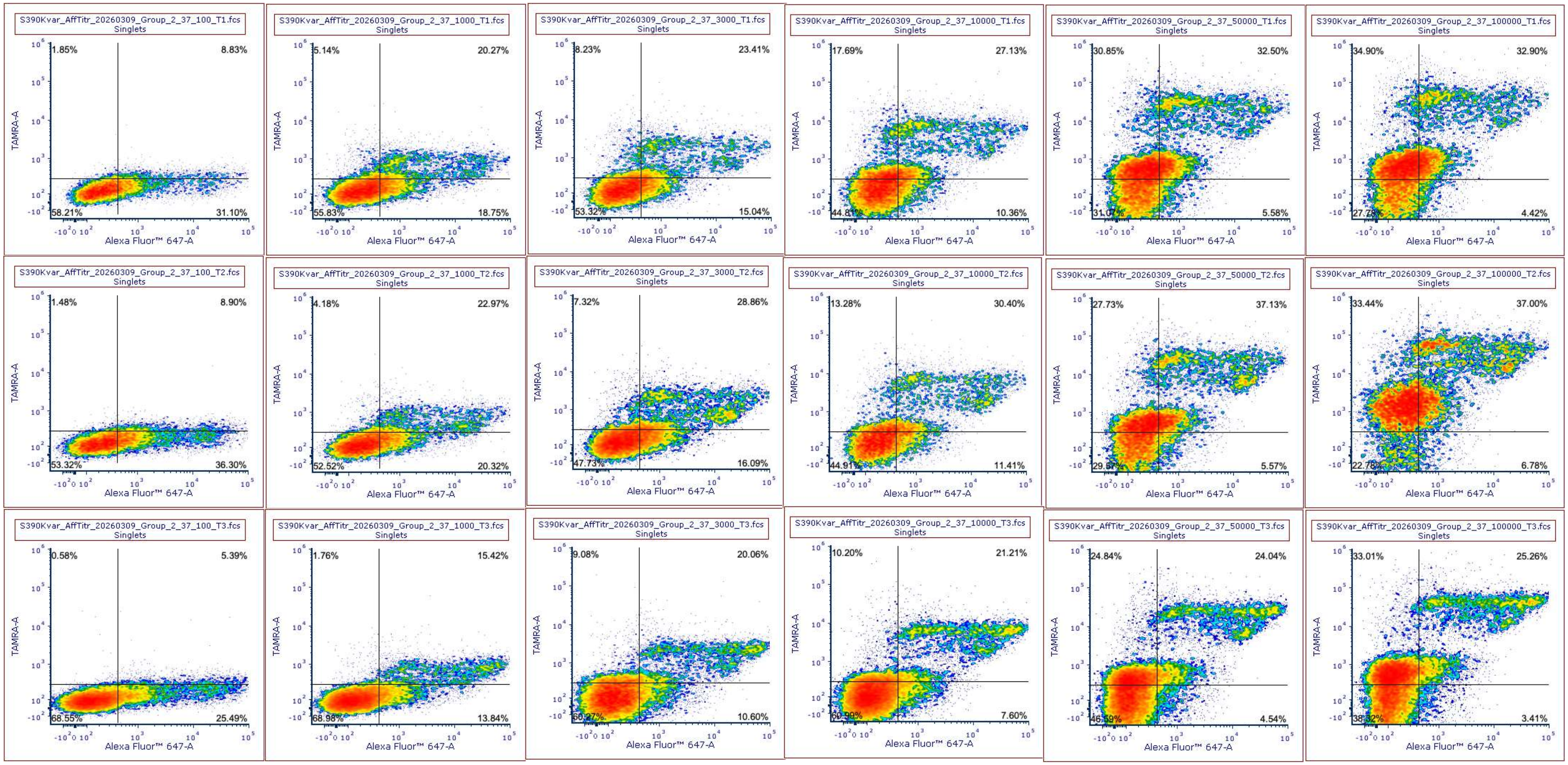

**Suppl. Inf. 5o - Figure 4: FLS2LRR-S390K - (Additive Stress – 0.1, 1, 3, 10, 50, 100 µM flg22-TAMRA column order from left to right)**

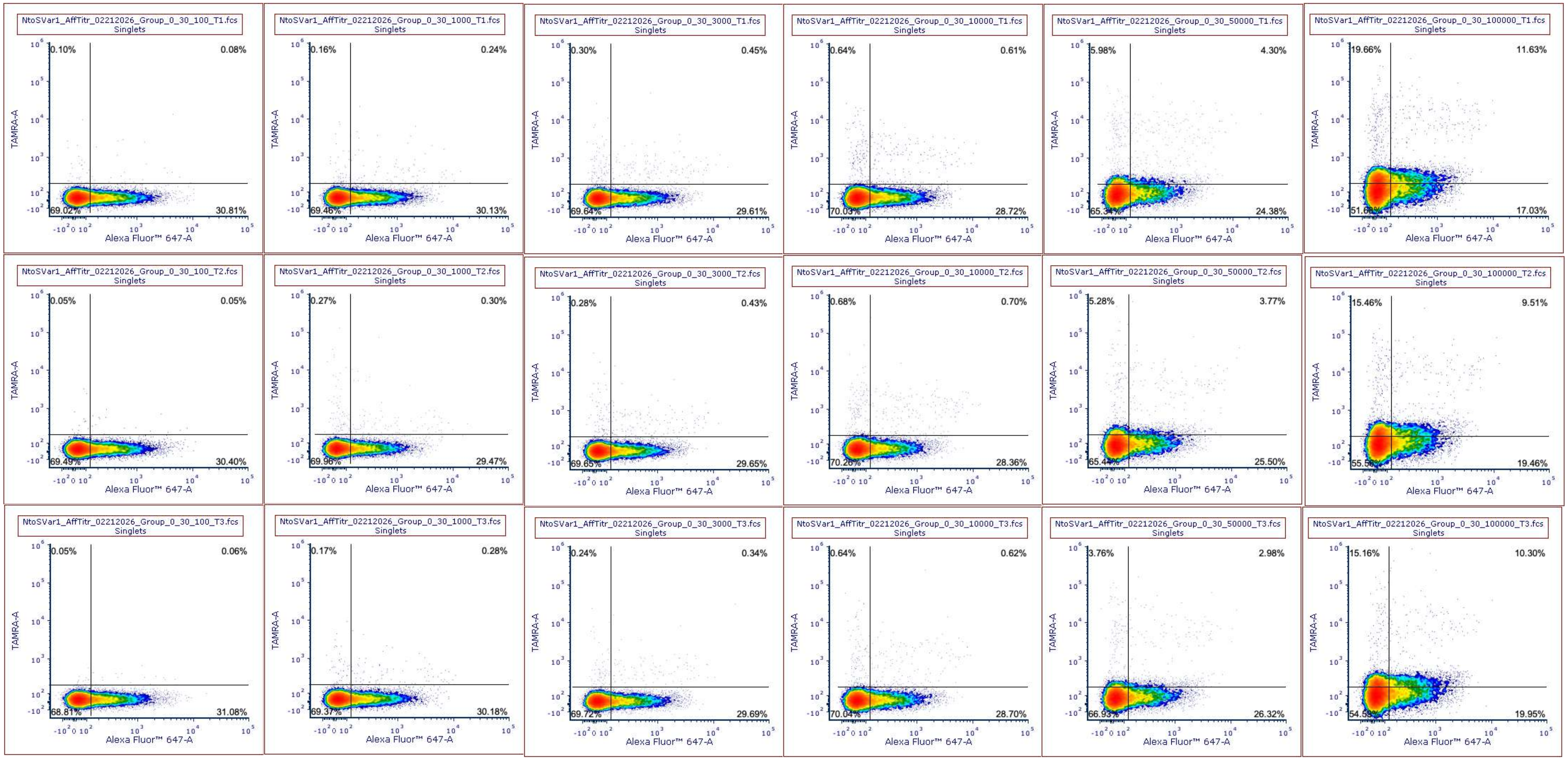

**Suppl. Inf. 5p - Figure 5C: Var1 - (No Stress – 0.1, 1, 3, 10, 50, 100 µM flg22-TAMRA column order from left to right)**

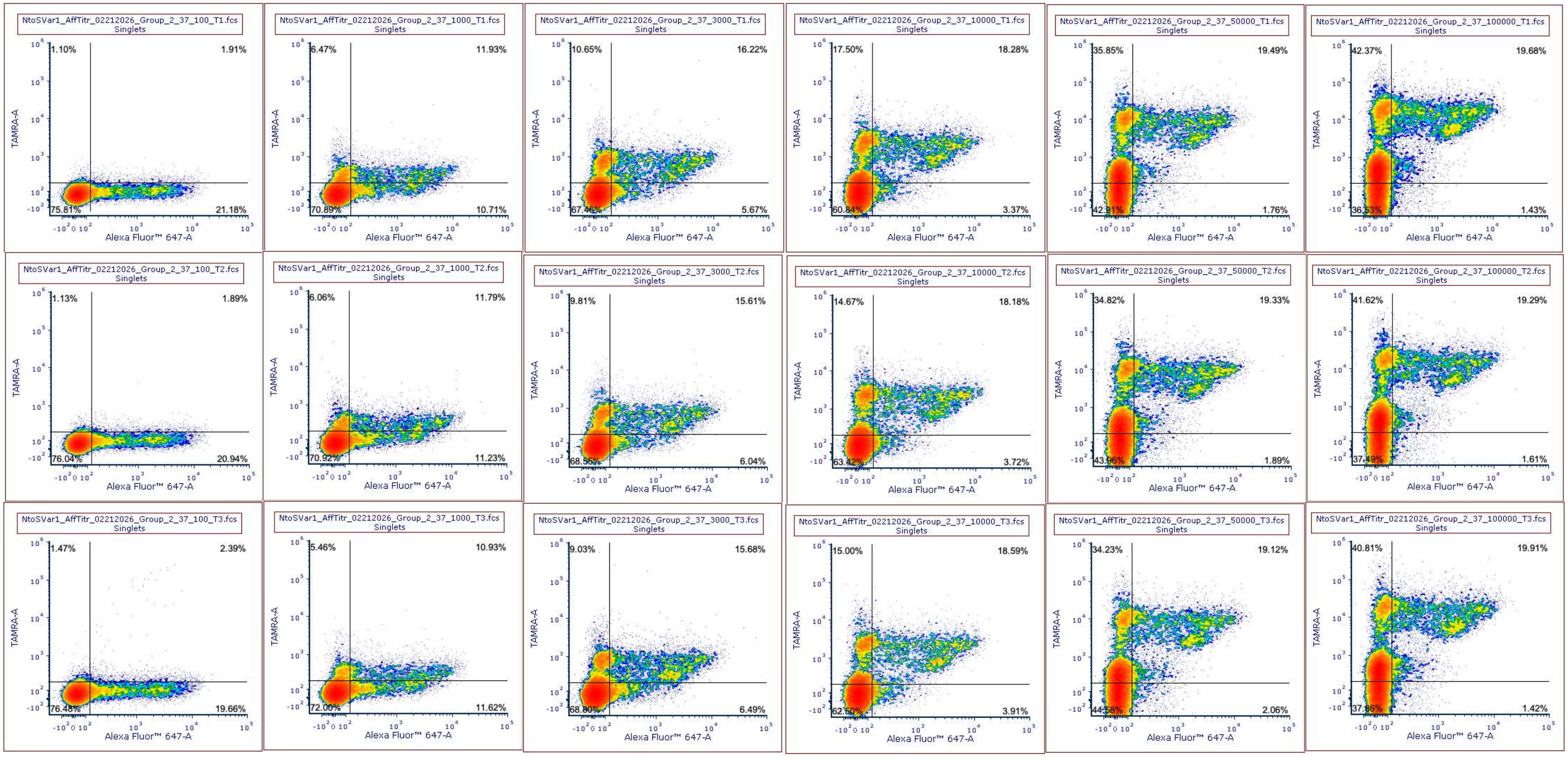

**Suppl. Inf. 5q - Figure 5C: Var1 - (Additive Stress – 0.1, 1, 3, 10, 50, 100 µM flg22-TAMRA column order from left to right)**
